# Democratizing three-dimensional surface phenotyping: an open structured-light platform reveals and removes the projection bias in biological imaging

**DOI:** 10.64898/2026.08.30.748077

**Authors:** Gregor J. Gentsch, Matthew Guo, Adrian Platz, Gunnar Brehm, J. Christopher Hennings, Christian A. Hübner, Andreas W. Stark, Christian Franke

## Abstract

Surface phenotyping underpins plant science, preclinical animal research and entomology, yet across all three the measurement is almost always a photograph, which records a projection and not the surface itself. Here we present the *Gentschinator3000*, an open structured-light platform that brings high-end metric surface measurement within reach of laboratories with no optics expertise, combining documented open hardware, open reconstruction software and analysis workflows for under 4000 Euro in components. It resolves a planar reference to 45 µm local flatness, registers full rotations to a loop closure of 156 µm, and performs stably across acquisition ranges that we define. Applying one workflow to a leaf before and after desiccation, to murine anatomy and to a spread lepidopteran, we find that projection underestimates surface area by 11 to 41 %. That error grows with the condition under study, with the evaluation scale and with the direction of view, so it can confound phenotype comparisons dramatically. In murine limbs a 15-degree change of viewing direction shifts a projected inter-segment angle by up to 23.2 degrees, while the three-dimensional angle does not move. Projection geometry can therefore contribute as much to a measured phenotype as the biology it is meant to quantify.

## Introduction

Many traits that biologists measure are properties of specimen surfaces. Leaf geometry influences boundary-layer exchange and light interception, and changes substantially during dehydration^1-3^. Three-dimensional (3D) measurement approaches have recently quantified functionally relevant variation in the hearing organs of geometrid moths^4^ and shown substantial projection bias in measurements of lepidopteran appendages^5^, while wing shape and size in spread Lepidoptera are commonly quantified from planar images^6^. In mammals, the geometry of the rodent vibrissal array determines which region of space an animal samples by touch, and we reconstructed that geometry with an early prototype of the platform described here to reveal the overlap between the tactile search space and the visual field represented in cortex^7^. Motor phenotyping in rodent disease models likewise rests on angles read from single video frames^8,9^. In each of these very different cases the quantity of interest belongs to a surface in space, and in each case, it is estimated from a projection of that surface.

The resulting bias is neither small nor random. A surface element inclined to the optical axis contributes less area to an image than it occupies in space, so any projected estimate of area, length, curvature or roughness is biased low by an amount that depends on the shape and pose of the specimen. Correcting for it would require knowing the shape, which is what the measurement is meant to establish. A single projection also cannot distinguish a surface from what lies behind it, so self-occluded geometry is absent rather than merely distorted. The practical consequence is that surface phenotyping often stops at ranked or categorical description, not because the underlying questions are soft but because the measurement cannot support more. The same limitation is recognised in high-throughput plant phenotyping, where projected proxies for leaf area and architecture depend on pose and canopy geometry as much as on the plant^10^.

Fluorescence microscopy passed through a comparable transition. Super-resolution methods are remembered for nanometre resolution, but arguably their more consequential effect was to make fluorescence measurements quantitative, because in single-molecule localisation microscopy the primary datum is a list of coordinates rather than an image, and quantitative spatial analysis and three-dimensional localisation followed from that change of representation as much as from the gain in resolution^11-13,17^. The nuclear pore complex illustrates the shift: its molecular-scale architecture was established by correlative super-resolution fluorescence and electron microscopy^14^, its eightfold symmetry has since become a reference standard for labelling efficiency, localisation precision and protein counting^15^, and the periodic actin-spectrin lattice of axons was revealed because localisation coordinates support periodicity analysis in a way a conventional image does not^16^. A calibrated three-dimensional point cloud stands in the same relation to a photograph. It is metric, it carries a confidence value for every point, and it enables operations an image does not: integration of surface area and curvature, roughness resolved across spatial scales, correspondence between specimens, and learning applied directly to unordered coordinates^18,19^.

Structured-light stereophotogrammetry produces such data by well-established means. Two calibrated cameras observe a scene illuminated by a projected pattern, correspondences between the two images are established by correlation, and each correspondence is triangulated to a metric coordinate. Over several decades these methods have matured into established metrology tools, with measurement precision ranging from sub-millimetre to micrometres depending on field size, optical geometry and implementation^20-23^. The obstacles to their use in biology lie elsewhere. Commercial high-end scanners are costly and supplied as sealed hardware and software, a poor match to specimens that are rarely standard. Open structured-light systems have already demonstrated that acquisition hardware and reconstruction software can be made accessible^24,25^, but their emphasis has largely been on geometry acquisition and engineering applications. Closer to biology, specialised platforms have digitised small organisms and put the resulting models to use: scAnt reconstructs arthropods by multi-view photogrammetry on off-the-shelf hardware^26^, such models have supported downstream tasks including the generation of annotated training data for animal detection and pose estimation^27^, and an automated all-side imaging device showed that whole-insect digitisation could be standardised^28^. Other efforts release open protocols that defer reconstruction to third-party software^29,30^. What is missing is a reproducible path from documented, modularizable hardware through reconstruction to biologically interpretable analysis, distributed in full, that reports where a reconstruction can be trusted and does not require the specimen to carry its own surface texture, the property that separates active pattern projection from the passive photogrammetry these platforms depend on^26,28^.

Open microscopy has shown how such obstacles fall. UC2, the OpenFlexure microscope and the miCube established that instruments of research quality can be assembled from commercially available components when the hardware design, the control software and the documentation are developed as a single deliverable^31-34^. None introduced a new physical or measurement principle. They changed who could make the measurement.

Here we apply that model to structured-light metrology. The *Gentschinator3000* (*G^3000^*) is an open, modular platform operated through a graphical toolbox and distributed with the open analysis notebooks used to produce every result below. We describe the build and the workflow a new user follows, characterise the instrument against a planar reference and report the acquisition ranges over which its performance is stable, which is rarely reported and which we found to be decisive for a non-specialist, and register multi-angle and full rotational views from a calibrated stage axis. We then apply the complete workflow to a leaf measured fresh and after desiccation, to murine surface anatomy across four animals, and to a spread lepidopteran, and show the same platform reconfigured for deep-ultraviolet illumination and for telecentric imaging at isotropic micrometre resolution^5,35^. **Figure 1** sets out the range of specimens the platform covers, the sampling achieved on each, and the projection error each of them carries.

**Fig. 1.**
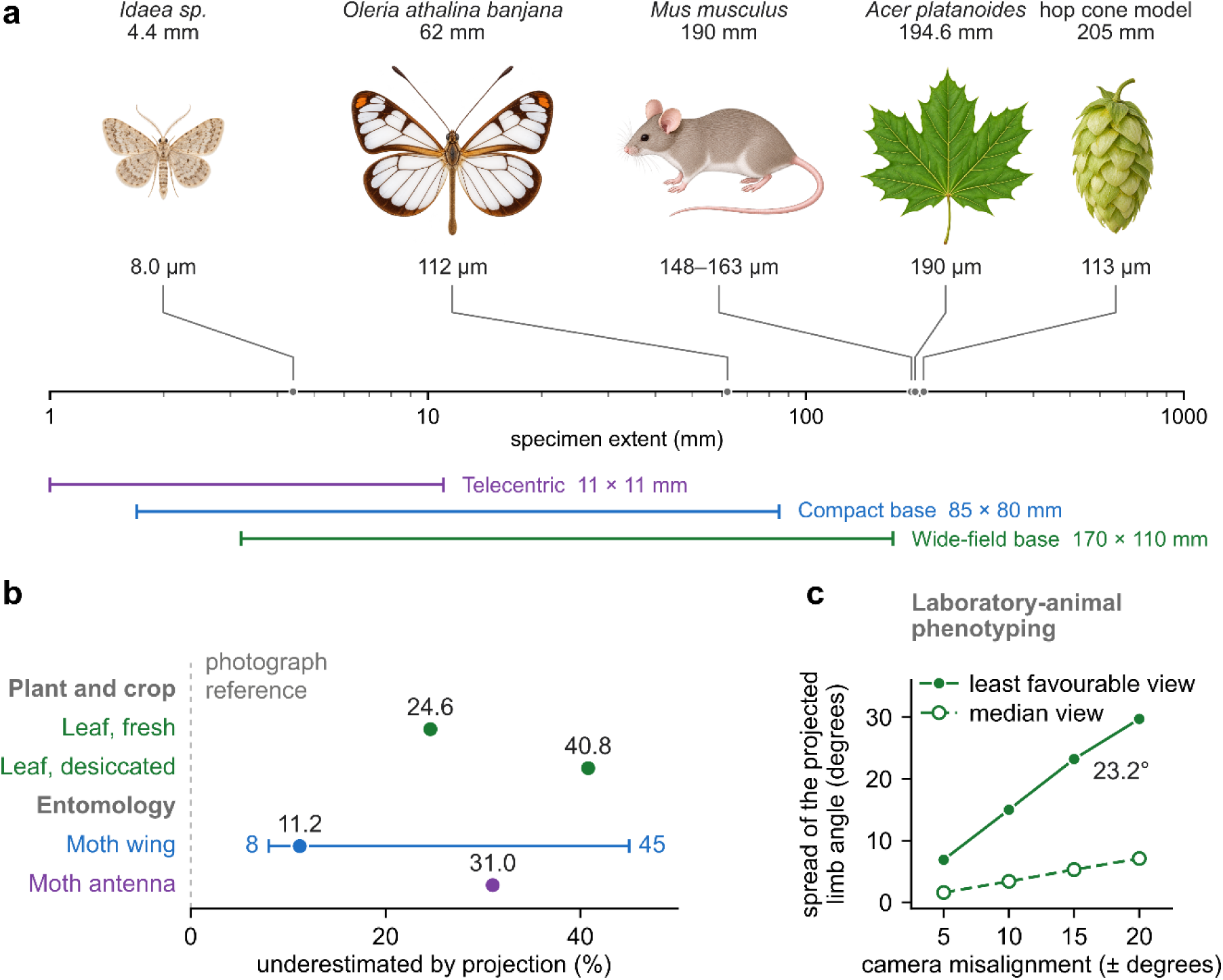
| Scope of the platform and the cost of projection. **a**, The five test objects referenced in this work, each placed on a logarithmic axis of specimen extent by a leader from its illustration, with the sampling achieved on it. Illustrations are not drawn to relative scale. Bars below give the field of view of the three optical configurations, all of which run on the same reconstruction pipeline. **b**, Underestimation of surface area, or of length for the moth antenna, when the same objects are measured from a single projection rather than from the reconstructed surface, grouped by the research domain each specimen stands for. The bar on the moth wing gives the range across regions within that one specimen. The dashed line at zero marks what a two-dimensional photograph reports. **c**, Spread of the projected inter-segment angle of the murine hind limb as a function of the camera misalignment permitted in azimuth and elevation, over the fourteen limb views, for the median view and the least favorable one. The three-dimensional angle does not change with viewing direction. Colors throughout denote the optical configuration used: purple telecentric, blue compact base, green wide-field base.

## Results

### Platform architecture and end-to-end workflow

The *G^3000^* combines two colour machine-vision cameras with variable-focal-length lenses, a digital light projector mounted between them, and a motorised rotary stage, assembled from 3D-printed parts and off-the-shelf components (**Fig. 2b**). The cameras view a common volume in a convergent arrangement, and stereo calibration of the configuration characterised here returns a convergence angle of 43.8 degrees between their optical axes. The measurement volume accommodates plant organs, arthropods, small rodents and isolated organs, and is rescaled by changing lenses and working distance without any change to the software.

**Fig. 2.**
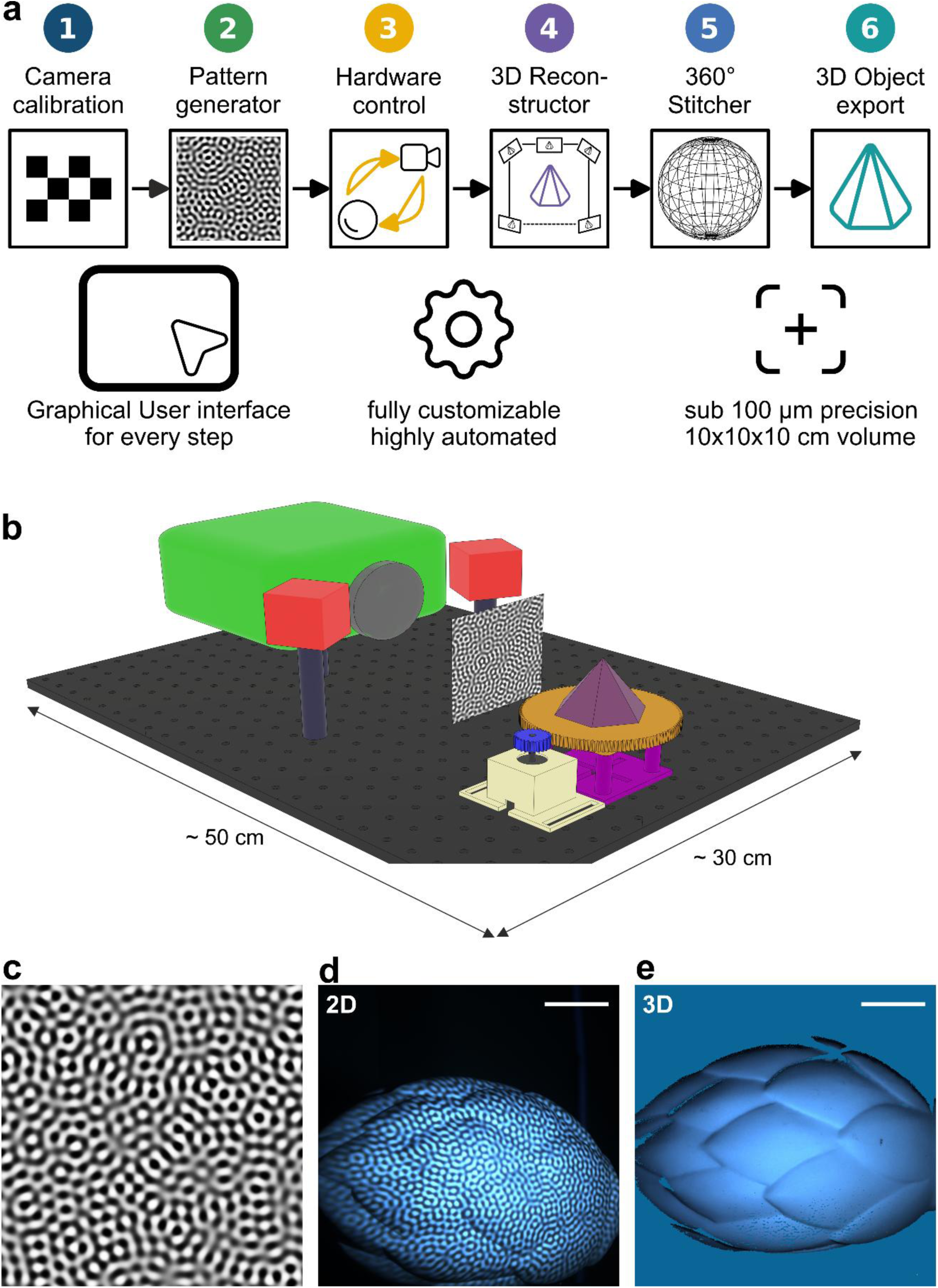
| The Gentschinator3000 platform and its workflow. **a**, The six software modules, in the order a user meets them: camera calibration, pattern generation, hardware control, single-view reconstruction, rotational stitching and export. Each is reachable both from the graphical interface and from source. **b**, The assembled system: two color machine-vision cameras (red) in a convergent arrangement, the digital light projector (green) between them, and the motorized rotary stage (orange and blue) carrying the specimen (purple), on an aluminum breadboard (dark grey) of approximately 50 by 30 cm. **c**, One realization of the band-limited random pattern used throughout, at 1000 by 1000 pixels and 50 pixels peak to peak. The working configuration projects 100 independent realizations. **d**, That pattern projected onto the hop cone model of Fig. 4, as recorded by the first camera. **e**, The single-view surface reconstructed from the correspondence between the two camera images of the pattern sequence, rendered as a shaded surface. Scale bars, 2 cm (**d, e**).

The motorised stage provides a minimum commanded rotation of 0.047 degrees, two orders of magnitude below the increments used for rotational scanning (**Methods**). Every non-standard part is supplied as a printable file dimensioned for common fused-filament printers and standard metric hardware. The material cost of a complete system was under 4000 Euro including tax at the time of purchase, of which cameras and lenses account for approximately 50 %. Modules can be exchanged independently, which enables the alternative configurations described below, and the frame can be dismantled, decontaminated and rebuilt, which matters for specimens that may not leave a containment laboratory or animal facility. Five implementations have been built to date, including systems assembled and operated by undergraduate students and one installed in the clinical research laboratory where the murine measurements reported below were acquired. The complete parts list, assembly sequence, wiring and firmware installation are given in **Supplementary Note 1** and **Supplementary Table 1**.

The custom software is written in Python and organised as six modules corresponding to the six steps of a measurement: camera calibration, pattern generation, hardware control, single-view reconstruction, rotational stitching and export (**Fig. 2a**). Each is reachable both from a graphical interface and from source, so a new user is never required to write, and an experienced user is never prevented from doing so, and a pre-recorded example project allows an installation to be verified before any hardware is connected. Every exported point retains the correlation coefficient at which its correspondence was found. That value travels through every subsequent stage of analysis as a per-point confidence channel, allowing the analyses below to state where a measurement is trustworthy.

Correspondences are encoded with two-dimensional band-limited random patterns (**Fig. 2c**) rather than with Gray-code or fringe sequences, building on band-limited random-pattern and statistical-speckle projection developed for stereophotogrammetric three-dimensional measurement (**Methods**)^36-38^. Confining the projected spatial frequencies to a narrow annulus makes every neighbourhood locally unique, so correspondence follows from correlation in two dimensions without a phase-unwrapping step, and a decoding failure remains local rather than propagating along a fringe order. That property matters for the discontinuous and self-occluding surfaces biological specimens present. Figure 1d and 1e show such a pattern on a specimen and the resulting single-view reconstruction.

### Reconstruction precision on a planar reference, and the settings that matter

We characterised reconstruction precision on a planar reference with a certified surface roughness below 1 µm, which separates instrument behaviour from specimen surface structure (**Fig. 3**). Three quantities are taken from the reconstruction after a fitted plane has been subtracted (**Methods**): the residual, a robust estimate of the spread of orthogonal distances to that plane; the local flatness, the root-mean-square residual at a 5 mm evaluation scale; and the completeness, the fraction of points assigned to the dominant planar surface rather than to a displaced population arising from false correspondence.

**Fig. 3.**
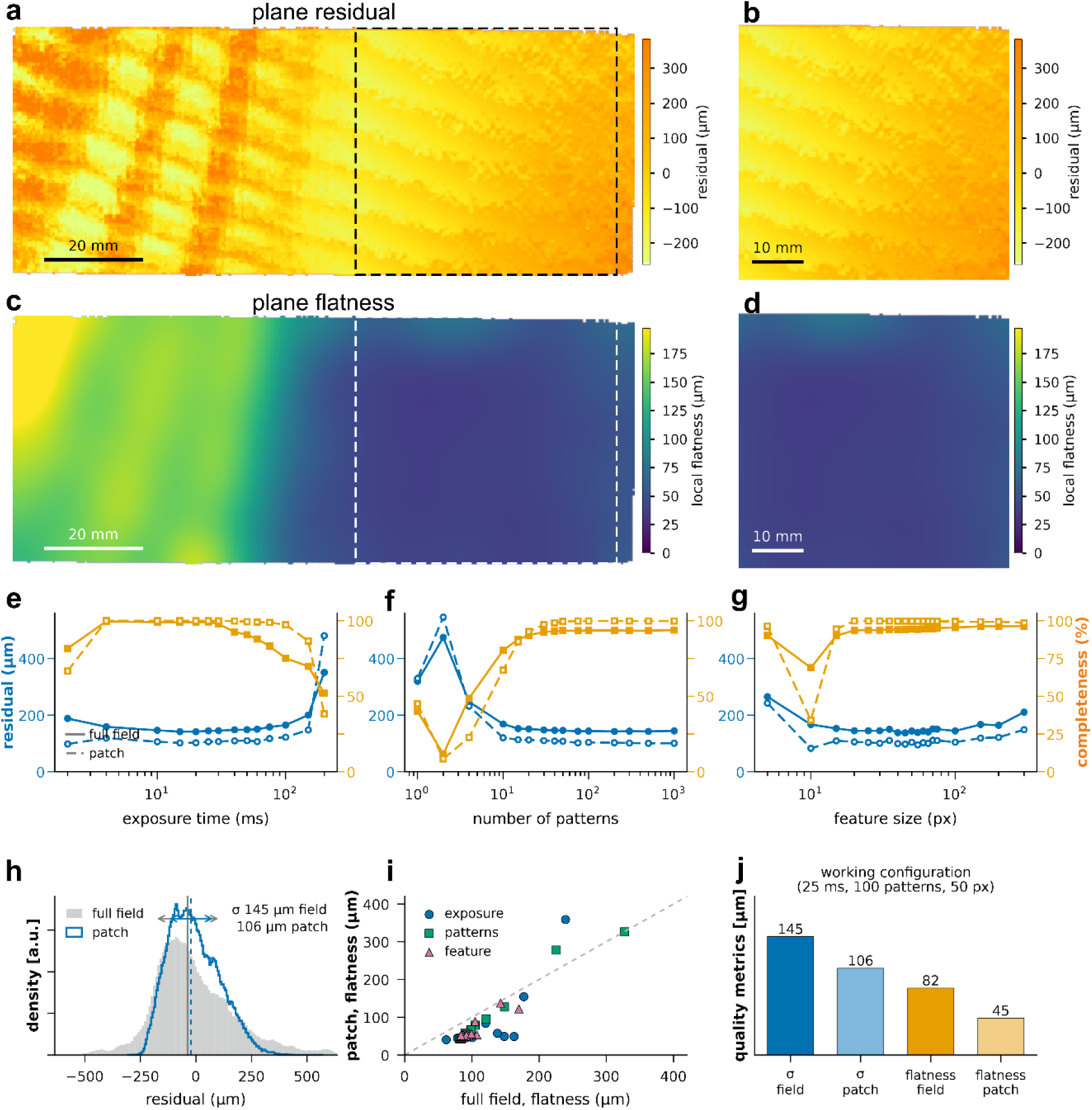
| Characterization against a planar reference with a certified surface roughness below 1 µm. **a**, Residual to the fitted plane over the full illuminated field, for an acquisition at the working configuration. The dashed box marks the 50 by 50 mm evaluation patch. **b**, The same map restricted to that patch. c, d, Local flatness of the same acquisition over the full field and over the patch, in the geometry of a and b. Color scales are shared within each row. The residual carries a weak periodic modulation aligned with the projection axis, strongest where the projected pattern is best focused (**Supplementary Note 5**). **e to g**, Robust residual (blue, left axis) and completeness (orange, right axis) against exposure time, number of projected patterns and pattern feature size, for the full field (solid, filled) and for the patch (dashed, open). In each sweep the remaining two parameters are held at the working configuration, and each point is a single acquisition. **h**, Distribution of the residual over the full field (filled) and over the patch (outline) at the working configuration, with the median and the robust interval marked for each. **i**, Local flatness of the patch against that of the full field for all 48 acquisitions, the dashed line is equality. **j**, The four quality metrics at the working configuration of 25 ms exposure and 100 patterns at 50 pixels peak to peak: residual 145 and 106 µm, local flatness 82 and 45 µm, completeness 99.2 and 100.0 %, full field and patch respectively. Scale bars, 20 mm (a, c) and 10 mm (b, d). n = 1 planar reference, 48 acquisitions.

All values are reported over two regions, the full illuminated field of 136 mm diagonal and a 50 by 50 mm evaluation patch defined once from the working-configuration acquisition and then held fixed for every sweep (**Methods**). Both are needed because the field is not entirely uniform. Local flatness varies fourfold across it, from about 160 µm at the near end to 40 µm across the far half (**Fig. 3a, c**), so where a specimen sits within the field is as consequential as how the acquisition is configured.

At the working configuration of 25 ms exposure and 100 patterns at 50 pixels peak to peak, the residual is 145 µm over the full field and 106 µm over the patch, the local flatness 82 µm and 45 µm, and the completeness 99.2 and 100.0 % (**Fig. 3j**). The full-field residual distribution is correspondingly broader and less regular than that of the patch (**Fig. 3h**). A weak periodic modulation aligned with the projection axis remains in the residual map, consistent with an intensity-gradient bias in sub-pixel correlation at sharp pattern edges rather than with a temporal artefact of the projector (**Supplementary Note 5**, **Supplementary Fig. 1**).

Three acquisition parameters can be adjusted without changing the optical configuration: exposure time, the number of patterns projected, and their characteristic feature size. Swept in turn, none of the three moves precision far, and each instead governs completeness (**Fig. 3e to 3g, Supplementary Table 3**).

*Exposure* is the least critical. Between 4 and 75 ms the full-field residual stays within 142 to 159 µm while completeness falls from 99.6 to 83.1 %, reaching 51.9 % at 200 ms. Once the brightest lobes of the projected pattern reach the top of the sensor range, the local contrast on which correlation depends is clipped, and correlation then returns spurious matches rather than none, so the surviving surface remains as precise as before while an increasing share of points is displaced away from it.

*Pattern number* produces the only genuine potential failure in the sweeps. Two patterns give 475 µm at 11.6 % completeness, because so few independent realisations leave neighbourhoods that are ambiguous rather than merely noisy, so the correspondence is unidentifiable rather than imprecise. Recovery is steep, reaching 169 µm at ten patterns and a plateau by thirty to forty, after which a twenty-five-fold further increase changes the residual by under 4 %. Independent realisations resolve ambiguity considerably faster than they average noise.

Feature sizes between 20 and 100 projector pixels peak to peak perform comparably, at completeness above 93.6 %. Outside that range the deterioration is not smooth. At 5 pixels the residual rises to 265 µm, consistent with attenuation of fine pattern structure by the combined modulation transfer of projector and imaging optics, and at 300 pixels too few independent features fall within a correlation window, and the correlation peak broadens, giving 210 µm.

The 10-pixel case shows why completeness cannot be omitted when assessing reconstruction quality. There the surviving points give a patch residual of 83 µm, the lowest value anywhere in the sweep and better than the working configuration achieves, while completeness over the patch collapses to 34.1 %. The third of the patch that survives is its best-conditioned part, so the apparent precision is a consequence of selective survival. On precision alone this acquisition would appear to be the best in the sweep; considered together with completeness, it is among the worst. This is why the correlation coefficient is retained throughout the pipeline as a per-point confidence value rather than discarded after reconstruction.

The three sweeps therefore define a broad operating region rather than an optimum, and the final configuration may depend on the specific specimen, while the working configuration used throughout the manuscript lies within all three plateaus. Precision is nearly invariant across that region while completeness is not, so an operating range defined from both quantities is a more useful criterion for routine measurement than a single best-case precision value.

### Registering a full rotation from the calibrated stage axis

Digitising an object from all sides requires views from several directions and merging them is the stage at which pipelines assembled from separate tools most often fail. Rather than register views on features of the object, which is unreliable for complex biological specimens, we recover the geometry of the stage once and use it thereafter. A planar chequerboard mounted off-axis is imaged at known angular increments through a full revolution, and the circular arcs traced by its reconstructed corners determine the direction of the rotation axis and a point on it in the camera frame, at a mean radial residual of 0.32 mm (**Fig. 4a**). The calibration holds until the stage is disturbed, and the transformation between consecutive views follows analytically from the known increment (**Methods**).

**Fig. 4.**
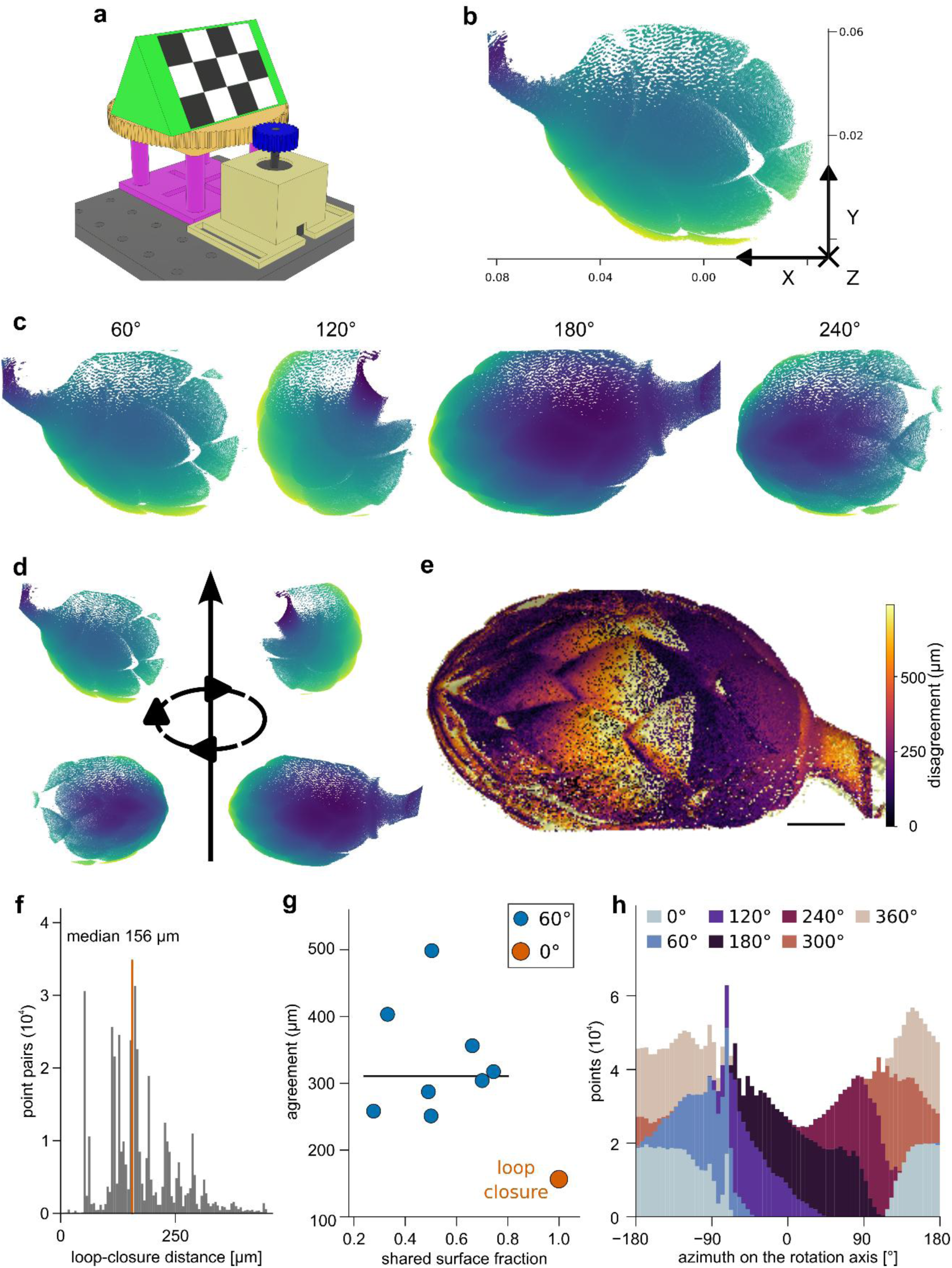
| Rotational scanning and registration from the calibrated stage axis. **a**, Axis calibration. A planar chequerboard mounted off-axis on the motorized stage is imaged at known angular increments, and the circular trajectories of its reconstructed corners give the direction and position of the rotation axis in the camera frame, at a mean radial residual of 0.32 mm. **b**, Single-view 3D point cloud reconstruction of a hop cone model, color-coded by depth. **c**, Four of the seven views, at the stage angles indicated. **d**, The views placed about the recovered rotation axis. **e**, The merged surface color-coded by inter-view disagreement, the distance from each point to the nearest point contributed by any other view, with a median of 230 µm. **f**, Loop-closure distance between the first and seventh views, which are acquired under the same viewing geometry after six 60 degree steps. The median of 156 µm over 546 000 point pairs is marked. **g**, Median agreement between view pairs against the fraction of surface they share. The point at unity is the loop closure, the eight remaining pairs are separated by 60 degrees, and their median is drawn as a horizontal line. **h**, Points contributed by each view as a function of azimuth about the rotation axis. Iterative refinement was disabled and no spatial filtering was applied. Scale bar, 20 mm (e). n = 1 specimen, 7 views.

We tested this on a hop cone model, whose overlapping bracts make it deeply self-occluding and leave it without globally distinctive features, the regime in which feature-based registration performs worst and a mechanical prior is worth most. Seven views were acquired at 60-degree steps so that the sequence closes on itself (**Fig. 4b to 4d**). Neighbouring views shared 28 to 74 % of their surface, and together they cover the observable surface continuously in azimuth, with every one-degree interval about the axis populated (**Fig. 4h**).

Six steps of 60 degrees return the seventh view to the direction of the first, so the two are acquired under the same viewing geometry. Their disagreement is an end-to-end measure of loop closure that combines repeated reconstruction uncertainty, stage repeatability and accumulated rotational error, and over 546,000-point pairs it is 156 µm in the median, 160 ± 116 µm in the mean, with a 95th percentile of 346 µm (**Fig. 4f**). Relative to the 205 mm bounding-box diagonal of the object this is approximately 1 part in 1300, obtained without any refinement of the analytical transformations and without filtering of 3D points.

Views separated by 60 degrees agree less closely, at a median of 311 µm and a range of 251 to 499 µm across the eight adjacent pairs (**Fig. 4g**). The zero-separation comparison gives the cleanest estimate of accumulated registration and repeat-acquisition error, because the two views sample the surface identically, whereas at 60 degrees the disagreement additionally reflects viewpoint-dependent sampling, since a region visible from two separated directions is seen at increasingly oblique incidence in at least one of them. Across the adjacent pairs the disagreement is unrelated to how much surface they share, so it follows the geometry of the specimen rather than the amount of common ground available to compare.

The same quantity mapped onto the merged surface is concentrated rather than distributed (**Fig. 4e**). Its median across the object is 230 µm, while the bract margins and the steeply inclined facets between them carry values several times higher, consistent with the poorer depth constraint expected at grazing incidence. No spatial filtering was applied at any stage, so the reconstruction reported here is the measurement rather than a cleaned version of it. Routine filtering options are described in **Supplementary Note 6**.

The merged cloud contains 2.94 million points at a median nearest-neighbour spacing of 113 µm (**Supplementary Table 3**). Screened Poisson reconstruction yields a mesh that is not watertight, because a rotational sequence cannot observe the supported underside of the object, and that exports directly into visualisation, mesh-processing and additive-manufacturing workflows (**Supplementary Fig. 2**).

Iterative refinement by generalised iterative closest point registration and pose-graph optimisation is available and disabled by default (**Methods**). On rigid, well-conditioned objects it improves loop closure only marginally, and on specimens that violate the rigid-surface assumption it can make the result worse, as we describe in the murine section.

With the precision of a single view and the reliability of a full rotation established, we turn to what the resulting surfaces support. The three applications that follow span different organismal groups, object geometries and analytical questions, and share one property: in each, the quantity of interest is conventionally taken from a photograph and can here be taken from the surface itself. They are demonstrations of measurement rather than biological studies, chosen also to show which quantifications are informative for a non-expert user and which can mislead (**Supplementary Notes 6, 9, 12**).

Registration from a mechanical prior rather than from surface features is what makes the platform usable on specimens that carry no distinctive texture, which describes most biological material.

### A leaf before and after desiccation

A leaf can be measured turgid and again after water loss, giving the same lamina before and after a large change in three-dimensional form, which makes it a direct test of projection bias. We scanned a single leaf of *Acer platanoides* fresh, and again after five days drying uncovered at ambient laboratory conditions, combining full rotational scans into two-sided reconstructions of both states (**Fig. 5a to 5f**). Because both faces of the lamina are represented, the fraction of the reconstructed surface visible from any single camera position can be calculated directly.

**Fig. 5.**
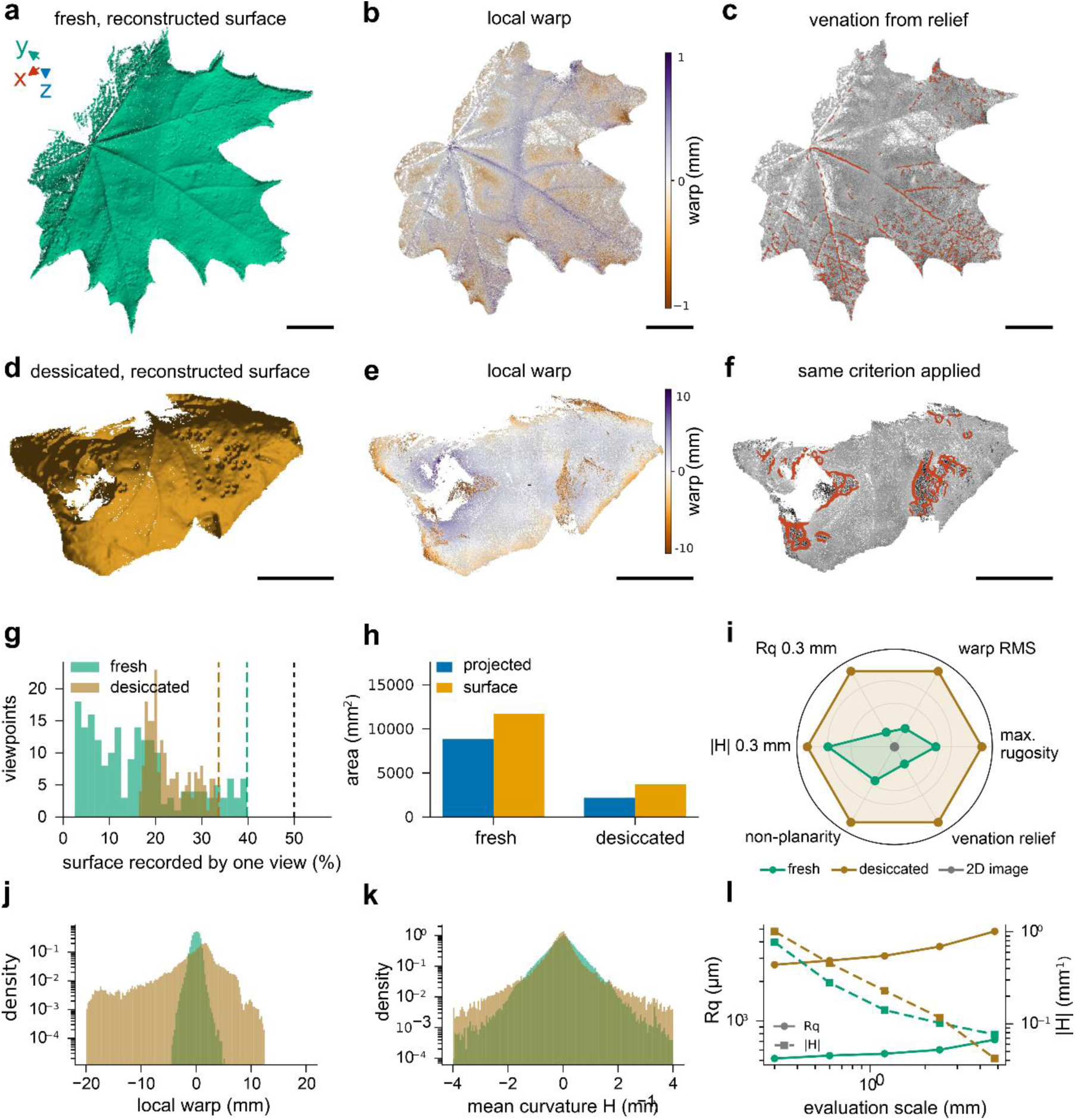
| A single leaf of *Acer platanoides* measured fresh (a to c) and after five days of drying (d to f). **a, d,** Reconstructed surface from the viewing direction that records the largest fraction of it. **b, e**, Local warp, the height remaining after removal of form at a 4.8 mm evaluation scale (note the different color ranges). **c, f**, Relief-based ridge detection, tracing venation on the fresh leaf and creases on the desiccated one under the identical criterion. **g**, Fraction of the reconstructed two-sided surface recorded by a single view, over 200 viewing directions on the sphere. Dashed lines mark the best direction for each state, and the dotted line the 50 % that one direction can expose at most for an opaque two-sided lamina. **h**, Projected against surface area for the surface visible from the best viewing direction, with the ratio R and the resulting projection error. **i**, Six descriptors for the two states and for a calibrated top-down image of the same leaf, each expressed as excess over a flat surface and normalized to the larger of the two states, so that the image lies at the origin on every axis. **j, k**, Distributions of local warp at a 4.8 mm evaluation scale and of mean curvature at 0.3 mm. **l**, Root-mean-square roughness (solid, left axis) and mean absolute curvature (dashed, right axis) against evaluation scale. Scale bars, 20 mm. n = 1 leaf measured in two states.

That fraction is the first cost of projection, which is not recoverable from photographic data alone. For an opaque two-sided lamina, a single direction can expose no more than one face-equivalent, corresponding to 50 % of the total surface. Swept over 200 viewing directions on the sphere, the best single view records 39.8 % of the fresh leaf and 33.6 % of the desiccated one, with medians of 14.6 % and 21.5 % and worst cases of 2.5 % and 16.3 % (**Fig. 5g**). Desiccation therefore does not simply make the leaf harder to photograph. It changes how strongly the measurement depends on where the camera stands: a flat leaf is excellent face-on and nearly invisible edge-on, while a curled one looks much the same from every direction, so the best-case falls while the median and the worst-case rise.

Even within the surface accessible from a given view, inclination compresses its apparent area. For the surface visible from the best viewing direction, the ratio of surface area to its own projected area is 1.326 fresh and 1.690 after desiccation, so projection underestimates the area of the visible surface by 24.6 % and 40.8 % respectively (**Fig. 5h**). The magnitude of the error is thus itself a function of the treatment under study, and a comparison of the two states drawn from projected images can be confounded.

Absolute areas set a limit on what any line-of-sight measurement can claim for a specimen of this kind. The area recovered in the full rotational reconstruction falls by 62.7 % between the two states and the projected footprint at the best viewing direction by 75.3 %, but neither figure represents loss of tissue: a curling leaf occludes much of its own lamina, so what is recovered after desiccation is the outward-facing part of a folded object even in a full rotation. Intensive descriptors of global form are therefore the ones to compare. The planarity index, the fraction of height variance accounted for by a fitted quadric surface, falls from 0.83 to 0.62, while the amplitude of the residual rises from 1.41 to 5.79 mm.

Roughness and curvature evaluated across scales separate two different and partly opposing aspects of the change (**Fig. 5l**). Root-mean-square roughness is five to seven times greater in the desiccated leaf throughout, rising from 518 to 720 µm fresh and from 2681 to 4813 µm desiccated between evaluation scales of 0.3 and 4.8 mm. Mean absolute curvature behaves differently: the desiccated leaf is more strongly curved at 0.3 mm, 1.01 against 0.77 mm^-1^, but less strongly curved at 4.8 mm, 0.042 against 0.077 mm^-1^, the two curves crossing near 2.4 mm. At fine scale the increased curvature of the desiccated lamina reflects the newly formed creases, whereas at coarse scale its broad folds carry less curvature than the lobes and raised midrib of the fresh leaf. A measurement at a single, intuitively chosen scale would have recorded one of these and concealed the other. The distributions behind the summary values carry information of their own (**Fig. 5j, k**). Local warp, the height remaining after removal of form at 4.8 mm, is sharply peaked in the fresh leaf, with a standard deviation of 1.41 mm and a kurtosis of 4.0. After desiccation it broadens to 5.79 mm and its kurtosis rises to 8.7, so the surface does not simply become rougher everywhere: most of it stays close to its local mean while a minority departs from it strongly, which is what folding into discrete lobes produces.

Venation is resolved on the fresh leaf as relief above the surrounding lamina, and a ridge criterion applied to that relief traces the midrib and the primary veins over 6.6 % of the lamina at a median height of 597 µm (**Fig. 5c**). Applied unchanged to the desiccated leaf, the same criterion returns creases rather than veins (**Fig. 5f**). After drying the folds reach several millimetres and no local relief criterion separates the two, so venation is reported for the fresh state only.

Taken together, these descriptors define a surface feature space that a two-dimensional projection cannot recover (**Fig. 5i, Supplementary Fig. 3**). When the same leaf is represented as a flat height field, warp, curvature, roughness, non-planarity and venation relief collapse to their flat-surface baselines and rugosity is unity by construction. Of note, 2D projection does not simply reduce the precision of these measurements, it removes the surface dimension on which they are defined.

Projected leaf area and canopy architecture are the routine currency of high-throughput plant phenotyping, where the treatment under study is frequently the thing that changes the geometry.

### Murine surface anatomy and the cost of measuring angles in projection

Mammalian specimens combine structures that differ strongly in scale, reflectance and scattering behaviour, from smooth skin and articulated limbs to dense fur and thin appendages, and they are the class of specimen on which an optical surface method is most easily overstated or misused. Large-scale external anatomy nonetheless carries phenotype, and it is here that the measurement itself can become the limiting factor. In beam-walking assays the foot base angle, the angle at which the sole of the hind paw meets the walking surface at the moment the contralateral leg is lifted, is a standard readout, and in models of hereditary spastic paraplegia it falls from roughly 75 to roughly 50 degrees^8,9,42^. Because the measurement is made on a projection, its value depends on where the camera stands relative to the animal as well as on the animal itself. The measurements below do not reproduce a beam-walk assay. The specimens are static and the hind paw is not resolved in most of them. What they are designed to do is to isolate the geometric problem underlying such image-based readouts, by asking how much a fixed three-dimensional limb angle changes when it is measured in projection.

We scanned four cadaveric mice, two homozygous for a knockout of an endoplasmic-reticulum-shaping protein and two wild-type littermates, all male and matched at 63 weeks of age (**Methods**). Each animal was shaved over the hindlimbs and rump and scanned from many directions on the rotary stage at a standoff of 1.1 m (**Fig. 6a**).

**Fig. 6.**
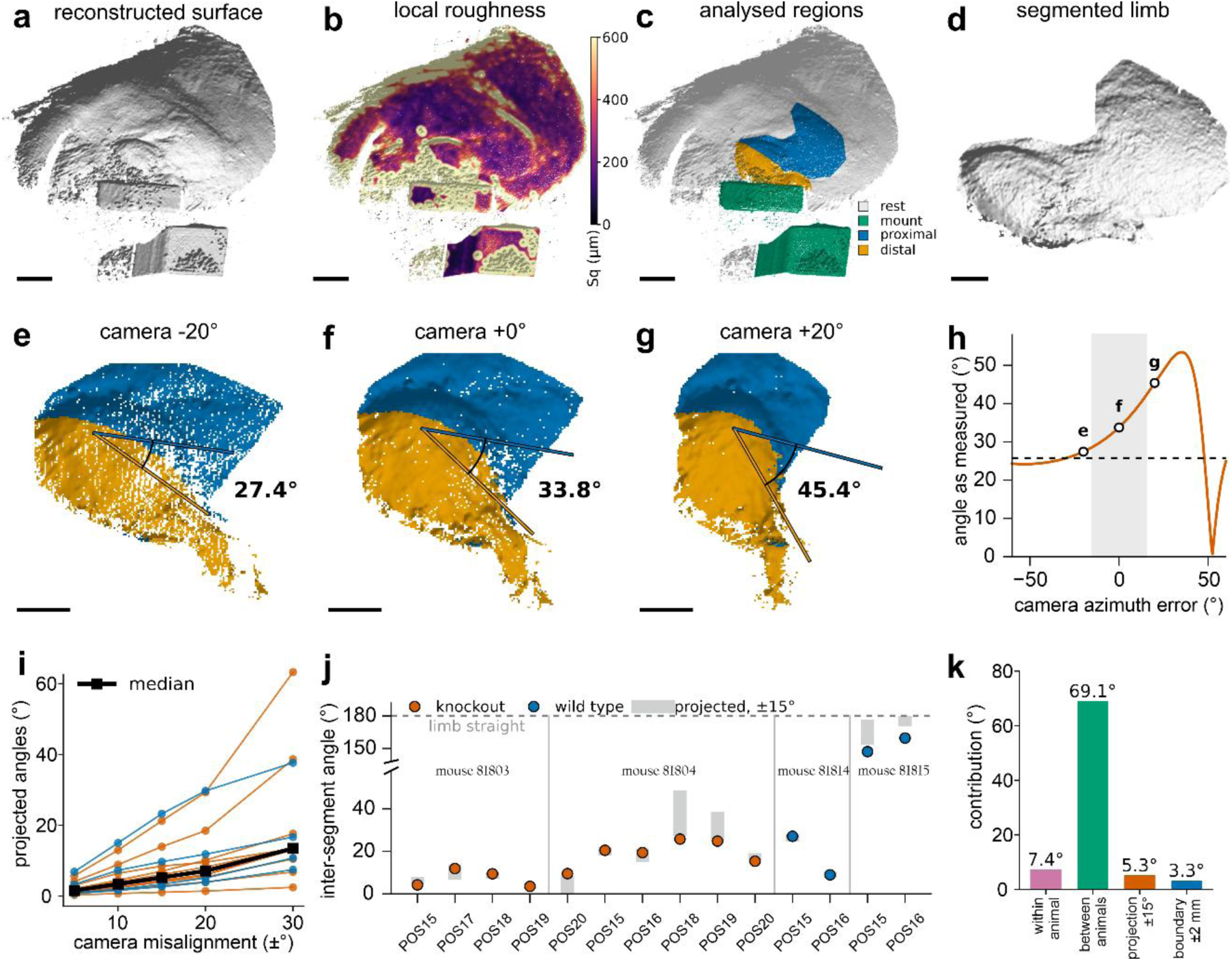
| Murine surface anatomy and the viewpoint dependence of projected limb angles. **a,** Reconstructed surface of a shaved animal. **b**, The same surface colored by local root-mean-square roughness, which contrasts exposed skin with fur, as fur returns a depth within a scattering volume rather than a surface. **c**, The analyzed regions in place within the animal: the mount, and the proximal and distal limb segments obtained by a manual cut at the visible joint bend. **d**, The segmented limb from a rotated viewpoint. **e to g**, One segmented limb rendered from three virtual camera positions 20 degrees apart, with the proximal segment in blue and the distal segment in orange. The two segment axes are drawn as they project into each image plane, and the angle an observer would measure between them is given. The actual, three-dimensional value is 25.8 degrees in all three. **h**, The projected angle for the same limb across a continuous sweep of camera azimuth, with the three positions of e to g marked and the three-dimensional value as a dashed line. Shading marks a virtual plus or minus 15 degree misalignment. **i**, Spread of the projected angle against the misalignment tolerance allowed, one line per view, median in black. **j**, Three-dimensional inter-segment angle per view, grouped by animal and colored by genotype, against the range of projected values within a plus or minus 15 degree misalignment (bars). 180 degrees would correspond to a straight/stretched out limb. Note the broken axis. **k**, Sources of angular variation compared: the mean within-animal standard deviation, the standard deviation of the four per-animal means, the median spread introduced by a plus or minus 15 degree change in viewing direction, and the median sensitivity to a plus or minus 2 mm displacement of the segment boundary. Scale bars, 20 mm (a to c) and 10 mm (d to g). n = 4 animals, 14 views.

Local surface roughness gives strong contrast between exposed skin and fur across the whole animal (**Fig. 6b**). Exposed skin returns 127 to 198 µm, close to the reconstruction noise floor at this standoff, whereas fur returns 307 to 704 µm and varies 1.8 times as strongly with viewing direction. The explanation is physical: fur is a scattering volume rather than a surface, so correlation returns a depth somewhere between the hair tips and the skin beneath, at a position that depends on the viewing geometry. An unsupervised classification separates the two regimes without training data or manual annotation (**Methods, Supplementary Note 10, Supplementary Figs. 4 and 5**).

In fourteen views a hind limb is fully visible. Each was divided into a distal and a proximal segment by a manual cut placed at the apex of the visible joint bend (**Fig. 6c, d**). The angle we report is that between the two segment axes, each directed away from their shared boundary, so 180 degrees corresponds to a straight limb and smaller values to a more strongly folded one.

That angle is a physical property of the limb and does not change with the viewing direction. Measured as a camera records it, the same angle does change. We rendered one limb from three virtual camera positions 20 degrees apart: the two segment axes project differently in each, and the angle an observer would draw between them reads 27.4, 33.8 and 45.4 degrees, while the three-dimensional value is 25.8 degrees throughout (**Fig. 6e to 6h**).

How much misalignment to allow the camera is a matter of experimental judgement, so we report the sensitivity as a function of it rather than at a single value (**Fig. 6i**). Across the fourteen views the median spread of the projected angle is 1.6 degrees for a misalignment of plus or minus 5 degrees in azimuth and elevation, 3.4 degrees at 10, 5.3 degrees at 15 and 7.1 degrees at 20, while the worst view reaches 6.9, 15.0, 23.2 and 29.7 degrees respectively. The spread is strongly view-dependent, increasing when the viewing geometry foreshortens the two segment axes unequally.

Because the three-dimensional segment axes are known, the projection error can be evaluated over a dense set of virtual viewing directions rather than only within a chosen tolerance band. Sampling over 600 directions on the sphere for each of the fourteen limbs gives a median error of 4.1 degrees and a 90th percentile of 17.5 degrees, with 55 % of directions below 5 degrees and 76 % below 10 (**Supplementary Fig. 7**). Nearly a quarter of camera placements would therefore mis-state the angle by more than 10 degrees, which is the size of the effect these assays are designed to detect^41,43^. The error is not distributed evenly but forms a structured band, smallest when the viewing direction is approximately normal to the plane spanned by the two segment axes and larger as one axis becomes increasingly foreshortened relative to the other.

The measured angles separate the four animals sharply (**Fig. 6j**). Three present a strongly folded limb, with per-animal means of 7.7, 18.0 and 21.2 degrees, while the fourth presents an almost straight one at 153.3 degrees. One knockout and one wild type sit at the folded end and one wild type at the extended end, so the spread follows specimen positioning rather than genotype. The cohort was neither intended nor powered for a genotype comparison, no genotype effect is tested or claimed, and with two animals per group a difference of the size reported for these assays would be indistinguishable from the positioning variation (**Supplementary Note 11**).

The relative magnitude of the terms involved can therefore be compared directly (**Fig. 6k**). Across the four animals the mean within-animal standard deviation of the measured angle is 7.4 degrees, combining repeated-view variation, postural relaxation during thawing and, in the one animal contributing both sides, left-right differences. The standard deviation of the four per-animal means is 69.1 degrees over a range of 146 degrees and is dominated by positioning. Against these, the median spread introduced by a plus or minus 15 degree change in viewing direction is 5.3 degrees and displacing the manually placed segment boundary by plus or minus 2 mm changes the angle by a median of 3.3 degrees. A purely geometric projection term is therefore of the same order as the variation within a single specimen, and larger than the median sensitivity to where the operator places the cut.

The same reconstructions support quantities a projection does not provide. Across the fourteen views the limb surface exceeds its own projection by a factor of 1.17 to 1.59, so an area taken from an image of a limb is short by a sixth to a third depending on how the limb happens to be turned. Surface texture is measurable at the same time and at a far finer scale: the caudal annuli, the ring-like folds of the tail skin, carry a relief of 60 to 140 µm peak to peak at a spacing of 1.39 ± 0.17 mm, and that spacing reads 17 % smaller in projection on average and 39 % smaller where the tail turns steeply away from the camera (**Supplementary Fig. 6**). A periodic texture is compressed in the same way an area is. What the murine data establish is what can be recovered from a mammalian surface, and how strongly an image-based geometric readout depends on the direction from which it is taken.

The same dependence applies wherever a posture, an angle or a body-surface property is scored from video, which is most of laboratory-animal phenotyping.

### A spread lepidopteran, and where the surface is lost

Lepidoptera and other winged insects are a case in which the cost of projection is easy to state and easy to underestimate. Wings are thin, inclined over much of their extent, and carry relief at several scales from venation down to the scale rows, while the standard morphometric measurements of wing shape and size are commonly taken from planar images of spread specimens^6^. We therefore reconstructed a spread lepidopteran (Nymphalidae: *Oleria athalina banjana*) and asked how much surface such a photograph would fail to record (**Fig. 7**).

**Fig. 7.**
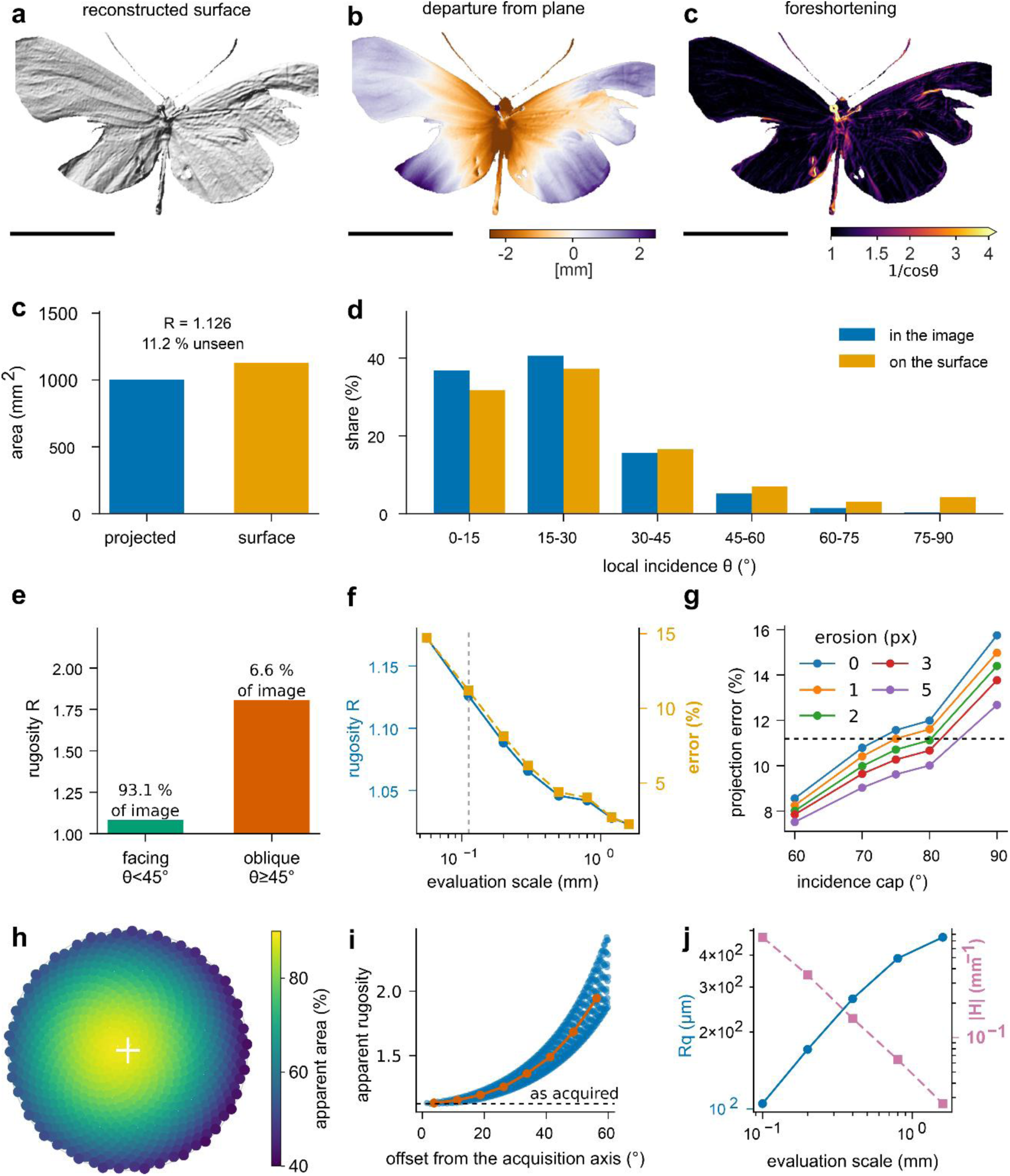
| Three-dimensional morphometry of a spread lepidopteran and the cost of projection. **a**, Reconstructed surface, rendered as a shaded surface from the acquisition direction. **b**, Departure from a plane fitted to the whole reconstruction. **c**, Local foreshortening factor, on a logarithmic scale. **d**, Projected against reconstructed surface area, with rugosity R and the resulting projection error. **e**, Share of the image and of the reconstructed surface contributed by each incidence band. **f**, Rugosity of the well-facing and oblique classes, split at 45 degrees of incidence, with the share of the image each occupies. **g**, Rugosity and projection error against the scale at which the surface gradient is evaluated. The dotted line marks the median point spacing, at which values are quoted, and the leftmost point lies below it. **h**, Sensitivity of the projection error to the two analysis conventions, across boundary erosion widths and incidence caps. The dashed line marks the value quoted in the text. **i**, Apparent area of the reconstructed surface as a fraction of its own area, for 750 virtual viewing directions within 60 degrees of the acquisition axis, in a Lambert azimuthal equal-area projection. The cross marks the acquisition direction. **j**, The same data as apparent rugosity against angular offset from the acquisition axis, with the binned median in orange and the value as acquired dashed. **k**, Root-mean-square roughness and mean absolute curvature against evaluation scale. Scale bars, 20 mm. n = 1 specimen, single view.

The reconstruction contains 1.50 million points over 62.0 by 35.5 mm at a median point spacing of 0.112 mm. Because the lateral coordinates are metric and depth runs along the optical axis, the cloud is a height field over a regular lattice, so every filled cell covers a fixed area in projection and the same area multiplied by the local foreshortening factor on the surface.

The specimen is not planar, although it was prepared to be, which is common in these specimens. Departure from a plane fitted to the whole reconstruction reaches 1.15 mm in the root mean square and 5.1 mm from the highest to the lowest point across a 62 mm span, at a planarity index of 0.72 (**Fig. 7b**). The wings rise towards their margins while the thoracic region sits below the fitted plane. Whatever its origin, this departure is present in the mounted specimen and therefore enters any measurement made from its projection.

Summed over the specimen, the reconstructed surface area is 1130 mm^2^ against a projected area of 1003 mm^2^, so the projection underestimates it by 11.2 %, at a median local incidence of 18.9 degrees and a median foreshortening of 1.06 (**Fig. 7d**).

Where that surface is lost matters as much as how much of it (**Fig. 7c, e**). Sorting the specimen by local incidence, surfaces inclined by less than 30 degrees carry 69 % of the reconstructed surface, while the steepest class beyond 75 degrees contributes 4.3 % of the surface from just 0.34 % of the image. Split at 45 degrees, the well-facing 93.1 % of the image has a rugosity of 1.082 while the remaining oblique fraction has a rugosity of 1.806 (**Fig. 7f**). Within the same specimen the area underestimate therefore ranges from about 8 % in well-facing regions to 45 % in oblique ones. Because a single view cannot contain occluded surface, and because the most steeply inclined facets are the least well sampled, the 11.2 % figure is a lower bound for the complete specimen (**Fig. 7h, Supplementary Table 7**).

Surface area derived from three-dimensional data is also not a single number. It depends on the scale below which structure is not resolved. At and above the median point spacing, rugosity falls from 1.126 at 0.112 mm to 1.089 at 0.2 mm and 1.023 at 1.6 mm, so the implied projection error falls from 11.2 to 2.2 % across that range (**Fig. 7g**). Any statement of surface area, or of the error a photograph makes in reporting it, is therefore incomplete without the scale at which it was evaluated. We quote values at the point spacing throughout, which is the finest scale the reconstruction supports.

The direction from which a specimen is recorded is a variable of the measurement in the same way. Projecting the reconstructed surface onto 750 virtual viewing directions within 60 degrees of the acquisition axis shows that the acquisition direction is close to the one that maximises apparent area, recovering 88.1 % against a maximum of 88.8 % (**Fig. 7i, j**). At 60 degrees from it the apparent area falls to 41.3 %, corresponding to an apparent rugosity of 2.42. For this specimen the apparent area is governed principally by angular offset from the acquisition axis, which accounts for 92 % of its variance, although the azimuthal dependence grows with offset and spans 41 to 68 % within the 45 to 60 degree band.

Again, the reconstruction supports measurements a calibrated image cannot give. Scale-resolved roughness rises from 105 µm at a 0.1 mm evaluation scale to 470 µm at 1.6 mm while mean absolute curvature falls from 0.75 to 0.026 mm^-1^ over the same interval, so amplitude and shape descriptors report different structural levels (**Fig. 7k, Supplementary Fig. 8**). Taken with the leaf, this specimen sets the lower end of the observed range: across the specimens measured here, projection underestimates reconstructed surface area by approximately 11 to 41 %.

Wing area and vein length taken from planar images of spread specimens underpin taxonomy, functional morphology and the digitization of natural history collections, all of which inherit this error.

### The same platform at other wavelengths and scales

Everything above applies to the base *G^3000^* configuration. Because illumination, imaging optics and reconstruction are separate modules connected only by calibrated geometry, the same architecture has been implemented in telecentric and deep-ultraviolet configurations without changing the reconstruction pipeline.

*Telecentric optics* in place of the endocentric lenses move the platform to the millimetre scale at isotropic micrometre resolution, reaching 8.0 µm laterally and 4.46 µm axially within an 11 by 11 by 6 mm^3^ volume. That is sufficient to reconstruct the body, wings and antennae of a moth of the genus *Idaea* under 4.4 mm in length, a specimen an order of magnitude smaller than the spread lepidopteran above, and it provides an independent measurement of projection error on a structure rather than a surface: the length of that antenna is underestimated by 31 % in projection^5^.

*Coherent speckle illumination at 266 nm* in place of the visible projector extends the platform to surfaces that return no usable texture in the visible^35^. Optically smooth glass produces no detectable pattern under visible illumination and fails outright, whereas deep-ultraviolet speckle generates sufficient backscatter for reliable correspondence on the same surfaces, and partially translucent biological material such as lepidopteran wing membrane reconstructs comparably well under both. Across the three configurations the platform spans two orders of magnitude in field of view and two decades in resolution on one software pipeline.

### The analysis is part of the instrument

A calibrated point cloud is not yet a result, and for the intended user it is not obviously closer to one than the specimen was. We therefore release, alongside the acquisition software, the analysis notebooks used for every application reported here (**Supplementary Note 9**). Each runs from an export to finished figures and tables with no intervention beyond selecting inputs and, optionally, a region of interest, and each reports the quality metrics on which its own output depends. These are not offered as a universal analysis, which does not exist, but as a working and readable starting point in the language the user will eventually have to speak. Metric coordinates with per-point confidence also admit spectral and graph descriptors, correspondence across specimens, and neural architectures that consume unordered point sets directly^18,19^.

## Discussion

The *G^3000^* brings structured-light metrology within reach of laboratories that have neither an optics workshop nor a computational group, while keeping the quantitative rigour that makes the measurement worth making. It measures a planar reference to a local flatness of 45 µm over a 50 by 50 mm evaluation patch and 82 µm over the full 136 mm field, registers a full rotation with a loop closure of 156 µm from the calibrated stage axis alone, and costs about an order of magnitude less than industrial optical scanners of comparable specification, with the whole chain from bill of materials to analysis notebook open. Structured-light scanning has been mature for a generation, its limited penetration into biology has never been a matter of physics, and the barrier that persists longest is not the instrument but the absence of a path from a point cloud to a number a biologist can use.

Our most consequential result is also the simplest. On a fresh leaf, on the same leaf after desiccation, on a spread lepidopteran and, from the telecentric configuration, on the antenna of a millimetre-scale moth, projection underestimates surface quantities by 24.6, 40.8, 11.2 and 31 % respectively. Because the magnitude of this bias varies with the condition under study, it can confound a comparison rather than merely add measurement noise, and within one specimen it varies from about 8 % on well-facing wing membranes to 45 % on oblique flanks. The descriptors that collapse to their flat-surface baselines in the two-dimensional leaf control make the point from the other side: warp, curvature, roughness and venation relief are not measured imprecisely by imaging, they are not measured at all. What changes with a metric surface is therefore not the accuracy of an existing number but which questions can be asked quantitatively.

At least two properties of these quantities have implications beyond the specimens measured here. The first is that a surface area is defined only once the scale below which structure is not resolved has been stated, so a value quoted without its evaluation scale is incomplete and two studies can be compared only at a common scale. The second is that the direction of view is itself a variable of the measurement, the acquisition direction for the lepidopteran specimen happening to be close to the one maximising its projected area while a 60 degree offset would have halved it. Neither property is peculiar to that specimen. Both follow from measuring a rough, curved object by projecting it, and both are invisible unless the surface is known.

The murine limb shows the same problem in its angular form, and points to where it will matter most in our opinion. There the quantity of interest is an angle rather than an area, and a change of viewing direction of 15 degrees shifts the projected value by a median of 5 degrees and by up to 23 degrees in the least favourable view, while the three-dimensional value does not change. Over the sampled sphere of viewing directions, nearly a quarter mis-state the angle by more than 10 degrees, comparable to the changes reported for established beam-walk readouts^9^. Misalignment of that size is not hypothetical: in a mouse model of hereditary spastic paraplegia the hind paws become markedly externally rotated during locomotion^42^, and two-dimensional human gait analysis shows mean absolute errors of 4.0 to 7.4 degrees across hip, knee and ankle angles against simultaneous three-dimensional motion capture^43^, under conditions far more controlled than a mouse crossing a beam. In a fixed specimen a misalignment is a setup error that could in principle be controlled. In a freely moving animal it is not an error at all but a variable the experiment cannot fix, because the animal turns as it crosses the beam and every frame therefore has a different viewing geometry. Nor is projection the only geometric error such measurements carry^41^, since skin moves relative to the skeleton: during rat locomotion, skin-derived joint angles have been reported to differ from X-ray-derived skeletal angles by up to 39 degrees at the knee around paw contact^44^. Separating these sources is necessary for interpreting image-based kinematics at all.

The three applications are demonstrations rather than studies, but each stands for a field in which the same argument applies: plant phenotyping, where projected proxies for area and architecture are routine and the bias is a function of the treatment, so that a drought or disease time course measures its own confound alongside its effect; laboratory-animal phenotyping, wherever a posture, an angle or a body-surface property is read from video; and entomology, where wing area and vein length underpin taxonomy and functional morphology. The telecentric configuration carries the same measurement to specimens of *Drosophila* scale and the deep-ultraviolet configuration to surfaces that return no usable texture in the visible.

We report the reliability of the platform deliberately, because in inexperienced hands an instrument that cannot report where it failed is more dangerous than one that is simply less precise. The parameter sweeps are presented as an operating range rather than an optimisation, since a user who is told which settings are safe does not need to know why they are safe, and the correlation coefficient travels as a per-point confidence channel through every stage, so unreliable regions are labelled rather than silently interpolated over. The murine data are the clearest case, returning a surface over the shaved regions, a documented refusal over the fur, and an unsupervised classification that separates the two from surface geometry alone. That classification may be the most immediately extensible part of this work, since adding a spectral channel, which the modular illumination already permits, would extend it towards separating materials that are all genuinely surfaces, and so towards automated segmentation of body regions and the monitoring of skin and coat condition in health and disease.

The main limitations follow from the physics. Strongly specular and highly transparent surfaces defeat triangulation in a single spectral band, and although the ultraviolet configuration widens the accessible class considerably, no single wavelength addresses the general case. Volumetric scatterers such as fur, feathers, hydrogels and optically cleared tissue return a depth that is not a surface. Being a line-of-sight technique, the method cannot see into deep cavities or undercuts, and a rotational sequence leaves the supported underside unmeasured, which is what makes absolute areas on the desiccated leaf properties of the acquisition geometry as much as of the object, and the lepidopteran projection error a lower bound. The current implementation assumes a static or quasi-static specimen, which is why the murine demonstration is made on cadavers rather than during locomotion, and carrying it into a freely behaving cohort requires faster projection together with motion-aware reconstruction, which we are pursuing. What else a designed phenotyping study would require is set out in **Supplementary Note 11**. The instrument is characterised against a planar reference but is not a certified coordinate measuring machine, and applications requiring formal traceability will need validation appropriate to that context. Several extensions follow directly from the architecture: a rigid reference artefact carried on the specimen mount would give a per-scan noise floor measured simultaneously with the specimen, additional cameras would reduce occlusion and convert the lepidopteran lower bound into a closed measurement, and multi-spectral projection would give material-specific contrast on heterogeneous tissue.

Our aim is less to improve the accuracy of any particular measurement than to change which laboratories can make it, and what questions they can ask once they have. We offer the platform as a documented, extensible and reproducible starting point, especially for those who are not experts in optics or programming.

## Methods

### Hardware

The platform comprises two color machine-vision cameras (AVT 1800 U-811c color C-Mount, 1458×1088 pixels) with variable-focal-length lenses (C-25-F1.8-10MP-T2-3), a digital light projector (TX-127) mounted between them, and a motorized rotary stage. Cameras are held in printed mounts on standard 25 mm posts and post holders and clamped to an optical table or aluminum breadboard of approximately 50 by 30 cm. The stage consists of a printed base carrying a lazy-Susan bearing and a printed 120-tooth gear driven through a printed 25-tooth pinion by a NEMA 17 stepper motor, giving a gear ratio of 4.8; a two-phase microstep driver set to 1600 pulses per motor revolution is commanded from a Raspberry Pi Pico running the Telemetrix firmware, so that the smallest commanded rotation is 0.047 degrees. The complete bill of materials, the assembly sequence, the wiring diagram and the firmware installation are given in **Supplementary Note 1** and **Supplementary Table 1**. For the configuration characterized here the stereo baseline is 705 mm at a working distance of 920 mm, giving a field of view of 170 mm by 110 mm and a convergence angle of 43.8 degrees between the optical axes.

### Camera calibration

Intrinsic and extrinsic parameters are estimated from images of a planar chequerboard held in varied positions and orientations across the measurement volume, using a pinhole model with radial and tangential distortion terms^45,46^. Corners are detected to sub-pixel accuracy and the calibration refined by minimizing reprojection error. Stereo calibration then estimates the relative pose of the two cameras. The chequerboard square size sets the metric scale of every subsequent reconstruction and must be entered correctly; the calibration walkthrough and its diagnostics are given in **Supplementary Note 3**. A calibration remains valid until a camera, lens, focus setting or aperture is changed.

### Pattern generation

Band-limited random patterns are generated in the Fourier domain. A field of complex Gaussian noise is multiplied by an annular band-pass filter whose radius sets the characteristic feature size and whose width sets the spectral bandwidth, and the inverse transform is taken. The resulting real field is mapped to the projector’s intensity range by a probability integral transform, which preserves the spatial statistics while using the full dynamic range. Each realization is generated from an independent noise seed, so the sequence carries no systematic structure. Feature size is quoted throughout as the mean peak-to-peak distance of the projected intensity field in projector pixels, which follows from the annulus radius. The working configuration uses 100 realizations of 1000 by 1000 pixels at 50 pixels peak to peak, preceded by a uniform white frame used for computing per-point intensity.

### Acquisition, correspondence and triangulation

The pattern sequence is projected and imaged synchronously by both cameras at fixed exposure. For each pixel of the first camera, the temporal intensity sequence forms a signature that is matched along the corresponding epipolar line in the second camera by normalized cross-correlation, refined to sub-pixel accuracy by parabolic interpolation of the correlation peak over three interpolation steps by default. The fundamental matrix from stereo calibration is estimated with the normalized eight-point algorithm under least-median-of-squares outlier rejection^47,48^, and images are rectified so that corresponding points share image rows^49^. Accepted correspondences are triangulated by linear least squares and refined by minimizing reprojection error. Every reconstructed point retains the correlation coefficient at which its correspondence was found. A threshold of 0.01 is applied at reconstruction so that the full correlation distribution is preserved for later gating, and each analysis reported here applies its own threshold, stated per application.

### Rotational registration

The rotation axis is calibrated once per hardware configuration. A planar chequerboard is mounted off axis on the stage and imaged at known angular increments over a full revolution; the reconstructed corner positions describe circular arcs about the axis, and coplanar circles are fitted to all corner trajectories simultaneously to recover the axis direction and a point on it in the camera frame. The mean radial residual of that fit is reported as the axis calibration error. Given the axis and the commanded angular increment, the rigid transformation between consecutive views follows analytically and is applied directly.

For the hop cone reconstruction, seven views were acquired at 60 degree steps and gated at a correlation coefficient of 0.50. Because six steps of 60 degrees return the seventh view to the direction of the first, those two views are acquired under the same viewing geometry, and the distribution of nearest-neighbor distances between them is reported as the loop closure. Agreement between other view pairs is computed on the subset of points whose nearest neighbor in the other view lies within 1 mm and is reported together with the fraction of points meeting that criterion, since pairs at large angular separation share almost no surface and any agreement figure computed on them reflects the gate rather than the measurement. Inter-view disagreement across the merged cloud is the distance from each sampled point to the nearest point contributed by any other view. Azimuthal coverage is the fraction of one degree bins about the rotation axis that contain points.

Iterative refinement is available and disabled by default. When enabled, each analytical transformation initializes multi-scale generalized iterative closest point registration^39^, which models the neighborhood of every point as a Gaussian, and the refined transformations form the edges of a pose graph closed by a loop edge and optimized globally^40^, implemented on Open3D^50^. Surfaces are reconstructed from merged clouds by screened Poisson reconstruction^51^ at depth 10 and exported as PLY, OBJ, STL or XYZ.

### Planar reference characterization

A planar reference with a certified surface roughness of better than 1 µm was scanned across a systematic sweep of three acquisition parameters: exposure time at 14 settings from 2 to 200 ms, number of projected patterns at 15 settings from 1 to 1000, and pattern feature size at 19 settings from 5 to 300 projector pixels peak to peak, each with the remaining two parameters held at the working configuration. Every point is a single acquisition.

Each reconstruction was reduced in three steps. First the dominant coplanar population was found by random sample consensus at a 0.5 mm inlier tolerance and refined by total least squares. Consensus rather than iterative trimming is necessary here: a trimmed least-squares fit is drawn onto a detached surface whenever that surface carries an appreciable share of the points, which on these acquisitions produces a plane that is wrong and a residual that is meaningless. Second, the points were assigned to three populations relative to that plane. Points more than 1 mm from it form the detached population, which appears as coherent sheets displaced from the surface and arises from false correspondences rather than from noise. Of the remainder, points within five robust scale estimates of the median form the bulk, which is the surface itself, and the rest form a near-plane spread concentrated at one end of the field. Completeness is the fraction of points in the bulk. Third, precision was quantified on the bulk alone, as a robust scale estimate given by 1.4826 times the median absolute deviation of the residual. A conventional standard deviation is not usable on these data because a small, detached population inflates it by an order of magnitude while leaving the surface unchanged.

These exports do not carry the per-point correlation coefficient, so the classification above is purely geometric. Where the coefficient is available, as in every other application reported here, it provides the same separation directly and at lower cost.

Local flatness was computed by rasterizing the bulk residual onto a 0.5 mm grid, removing form by normalized convolution at a 5 mm Gaussian width and taking the root-mean-square of what remains, again by normalized convolution so that unfilled cells do not bias the estimate. Two regions of interest are reported. The full illuminated field is the extent of the bulk population, 136 mm across its diagonal. The evaluation patch is the largest square region whose median local flatness lies within 20 % of the lowest value attainable anywhere in the field, determined once by exhaustive search over window size and position on the working-configuration acquisition, then converted to a fixed box in world coordinates and applied unchanged to every acquisition, the reference having remained in place throughout. That procedure returns a 50 by 50 mm region.

### Normalized convolution

Several analyses require a smoothed version of a field that is defined only where data exists. Throughout, this is computed as normalized convolution: the Gaussian smoothing of the field with unfilled positions set to zero is divided by the Gaussian smoothing of the binary mask of filled positions, and the quotient is retained only where the smoothed mask exceeds 0.2. This avoids the bias that interpolation across unfilled regions introduces, which is substantial when the filled fraction is well below unity, and it is used for form removal, for local roughness, and for the smoothed height fields from which gradients and curvatures are taken.

### Leaf specimens and analysis

A single leaf of *Acer platanoides* was scanned in a full rotational sequence, then left uncovered on the optical table for five days at ambient laboratory temperature and humidity and scanned again in the same way. Both faces of the lamina are represented in each merged reconstruction. Point clouds were reduced to a 0.25 mm voxel grid, the largest connected component at a 1.0 mm linkage distance was retained, and the specimen holder was removed by an excess-green index computed from the per-point color, retaining points above a threshold of 0.10. This left 478,121 points in the fresh state and 177,505 after desiccation, at a median point spacing of 0.19 mm in both.

The fraction of the reconstructed surface recorded by a single view was computed over 200 directions distributed on the sphere by a spherical Fibonacci lattice. An area is attributed to each point as a disc of diameter equal to the median distance to its six nearest neighbors, and the reconstructed surface area is the sum of those areas. For each direction, points whose normal lies within 75 degrees of the camera are projected onto a 0.6 mm lattice perpendicular to it; a point is recorded when its depth lies within 1 mm of the nearest such point in its cell, so that surface hidden behind other surface contributes nothing. The recorded fraction is the summed area of recorded points divided by the reconstructed surface area. For an opaque two-sided lamina, a single direction can expose at most one face-equivalent, so 50 % is a geometric reference rather than an empirical ceiling.

Surface descriptors were computed from the viewing direction that records the largest fraction of the surface. Points whose normal lies within 75 degrees of that direction were rasterized as a height field at 0.15 mm. Rugosity is the ratio of the summed foreshortened area of that face to its projected area. The planarity index is the fraction of height variance accounted for by a fitted quadric surface. Local warp is the residual after removing form at a 4.8 mm evaluation scale, and mean curvature is computed from the first and second fundamental forms of the height field smoothed at 0.3 mm; both scales are stated wherever those quantities are reported, because the sign of the difference between states reverses with scale for curvature. Roughness parameters follow ISO 25178 ^52^ and were evaluated at Gaussian widths from 0.3 to 4.8 mm. Local shape was classified as bowl, saddle or dome from the signs of the principal curvatures.

Venation was segmented by a ridge criterion rather than by a relief threshold, which does not separate the network from the surrounding lamina. Relief was taken above a 2 mm detrend, smoothed at 0.45 mm, and the more negative principal curvature of that relief field used as a ridge response. Pixels above the 88th percentile of ridge response and the 55th percentile of relief were retained, morphologically closed, and components smaller than 0.6 mm^2^ discarded. The identical criterion was applied to both states.

### Murine specimens and segmentation

Cadaveric material from four male mice, two homozygous for a knockout of an endoplasmic-reticulum-shaping protein and two wild-type littermates, all 63 weeks of age, was obtained from a colony maintained by the Hübner laboratory (Institute of Human Genetics, Jena University Hospital) in the course of its hereditary spastic paraplegia research programme. No procedures were performed on living animals for this study, no animals were bred or killed for it, and all measurements were made post mortem on carcasses surplus to that programme. No separate animal experimentation approval was therefore required. Each animal was shaved over the hindlimbs and rump and scanned from many directions on the rotary stage at a ground sampling distance of 0.148 to 0.163 mm and a standoff of 1.13 to 1.18 m. No live animals were exposed to structured illumination in this study. Specimens were thawing over the course of each session, so posture was not identical between views, and the animals were not positioned identically between sessions; both are treated explicitly in the analysis rather than corrected.

Point clouds were exported at a correlation threshold of 0.25. Hind limbs, the tail and a flat surface fixed in the laboratory frame were segmented manually in CloudCompare, and each limb was subdivided into a distal and a proximal segment by a freehand cut placed at the apex of the visible joint bend. No anatomical landmarks were identified, and the hind paw could not be resolved in these acquisitions. Segments are therefore referred to as distal and proximal rather than by anatomical name, and all reported limb quantities are geometric. Views in which the limb was not fully visible, or in which the segmentation could not be placed with confidence, were excluded; fourteen views across the four animals were retained.

### Range-image analysis and surface classification of murine scans

Each view was rasterised onto the regular lattice defined by the first-camera pixel coordinates as a structured range image, with the correlation coefficient carried as a per-pixel confidence channel. Correspondences below a correlation coefficient of 0.50 were discarded, a depth gate was applied as the 0.5 to 99.5 percentile interval of the retained depth distribution widened by 15 % of its width, and the dominant connected component of the resulting mask after morphological closing was taken as the object. Ground sampling distance was obtained from the median world-space distance between adjacent lattice positions.

Large-scale form was removed by normalized convolution and roughness quantified as the local root-mean-square of the residual computed the same way. Principal curvatures, curvedness and the surface normal were computed from the first and second fundamental forms of the smoothed range image rather than from a Laplacian, since the lattice is a camera raster on which slopes of order one are common. Areal parameters follow ISO 25178 ^52^ and were computed separately within each surface class. Surface regimes were defined from the joint distribution of locally smoothed correlation and local roughness, spatial coherence of each class was assessed by connected-component analysis after morphological opening, and separability was quantified as the area under the receiver operating characteristic, reported only for quantities that do not enter the class definition. No training data, manual annotation or prior knowledge of the scene was used. Full results are given in **Supplementary Note 10**.

### Limb segment axes and angles

Each segment axis is the first principal direction of its point set, signed to point away from the junction between the two segments, which is located as the centroid of the distal points nearest the proximal set. The reported angle is the angle between these two axes, so that 180 degrees corresponds to a straight limb and smaller values to a folded one; it is invariant to viewing direction and does not use the reference plane. The projected angle for a given viewing direction is obtained by removing the component of each axis along that direction and measuring the angle between the remainders, which is the operation an observer performs when drawing two lines on an image; the projection is undefined where either axis lies within 15 degrees of the viewing direction and those directions are excluded. Projected angles were evaluated over azimuths from −90 to +90 degrees at the nominal elevation, over misalignment boxes of plus or minus 5, 10, 15, 20 and 30 degrees in azimuth and elevation, and over 600 directions distributed on the sphere by a spherical Fibonacci lattice, the last reported as the absolute deviation from the three-dimensional value.

The reference plane was taken from a flat surface fixed in the laboratory frame and visible in every view. Because it presents several faces, planar faces were extracted sequentially by random sample consensus at a 0.5 mm inlier tolerance, refined by total least squares, and the face closest to a direction common to all views retained. Its normal is reproducible to between 0.02 and 0.64 degrees across the views of a session, at a plane residual of 0.14 to 0.21 mm. It differs by approximately 7 degrees between the two sessions, which were four days apart and involved rebuilding the arrangement, so it is a within-session frame and no angle is compared across sessions through it.

Sensitivity to the placement of the segment boundary was quantified by defining a planar cut through the junction whose normal points from one segment centroid to the other, displacing it from −4 to +4 mm in 1 mm steps, reassigning points accordingly and recomputing both axes. Axis signs were held to the unshifted solution so that a shortened segment cannot flip. The reported sensitivity is the range of the angle over displacements within the stated tolerance, a median of 3.3 degrees within 2 mm and 6.0 degrees within 4 mm.

### Limb surface and projected area

The two segments were combined and rasterized onto a 0.35 mm grid perpendicular to the nominal viewing direction, keeping the point nearest the camera in each cell. Projected area is the filled cell count multiplied by the cell area. Surface area is the same sum weighted by the local foreshortening factor, the reciprocal cosine of the angle between the surface normal and the viewing direction, computed from the smoothed depth field, with cells beyond 75 degrees of incidence excluded because the factor diverges as a surface turns edge on. Rugosity is the ratio of the two.

### Caudal annuli analysis

Tail segments were aligned to their own principal axes and rasterized at 0.08 mm onto the plane of the two dominant axes, with depth as the third. The densely sampled central strip was retained, form was removed by normalized convolution at a 0.5 mm Gaussian width, and the residual relief transformed by a two-dimensional fast Fourier transform under a Hann window. The reported annulus spacing is the reciprocal of the radial frequency of the strongest peak between 0.25 and 1.6 mm, and its prominence the excess of that peak over the median spectral power in the same band. Views whose peak prominence fell below 14 dB were rejected, since below that threshold the returned spacing is not merely noisy but wrong: peak prominence and reported spacing correlate at +0.78 across the twelve tail segments, and all four rejected views return spacings outside the range recovered by the accepted ones. Eight of twelve views were accepted.

For display, the annuli were also profiled directly. A profile taken along the tail axis does not recover them, because the crests run obliquely and the tail curves, so that averaging along the axis cancels a 60 µm modulation against several hundred micrometers of tail irregularity. The local wave vector was therefore obtained from the two-dimensional spectrum of a dense window and the points binned by their projection onto it, which gives a profile perpendicular to the crests. The spectral spacing is the quoted measurement; the counted crests corroborate it and run systematically slightly larger, because the smoothing required to display them merges shallow adjacent crests. The apparent spacing in projection was computed per view as the surface spacing multiplied by the cosine of the local incidence angle over all retained cells below 75 degrees and is reported as its median and 10th to 90th percentile range.

### Lepidopteran analysis

The lepidopteran specimen (Nymphalidae: *Oleria athalina banjana*) was collected in 1999 in Ecuador, dried, and months later remoistened, spread on a spreading board and dried again^53^. Its wings are slightly damaged, probably due to a bird attack (G.B., personal observation). It was scanned in the standard visible configuration at a working distance of 256 mm, with correspondences gated at a correlation coefficient of 0.70 on export. The specimen holder lies behind the specimen along the optical axis and was removed by a depth threshold at 264 mm, retaining 1,496,204 specimen points against 112,832 belonging to the holder.

Because lateral coordinates are metric and depth runs along the optical axis, the export is a height field over a regular lattice, and points were rasterized at 0.112 mm, the median point spacing. Projected area is the filled cell count multiplied by the cell area; surface area is the same sum weighted by the local foreshortening factor computed from the smoothed height field. One lattice cell was removed at the object outline before any derivative was taken, because cells at the silhouette report the step from surface to background rather than a real slope, and cells beyond 75 degrees of incidence were excluded because the foreshortening factor diverges as a surface turns edge on. The sensitivity of the resulting projection error to both conventions was mapped across erosion widths of zero to five cells and incidence caps between 60 and 90 degrees.

The gradient from which foreshortening is computed depends on the width at which the height field is smoothed, and with it the surface area. Rugosity was therefore evaluated at Gaussian widths from 0.056 to 1.6 mm and is reported as a function of that width, with values quoted at the median point spacing of 0.112 mm, the finest scale the reconstruction supports. The value at 0.056 mm lies below the nominal sampling scale and is shown only to illustrate the sensitivity to undersmoothing.

Departure from planarity was taken as the residual after fitting a plane to the whole reconstruction by least squares, and the planarity index as the fraction of height variance that fit accounts for. The dependence of apparent area on viewing direction was computed by treating each retained lattice cell as a facet with area equal to its foreshortened area and normal taken from the smoothed height field and summing the facet areas weighted by the absolute cosine between facet normal and viewing direction, over 750 directions distributed within 60 degrees of the acquisition axis by a spherical Fibonacci lattice. This quantifies how the apparent area of the reconstructed surface changes with viewing direction; it is not a simulation of a new acquisition, since surface occluded in the original view is absent from the reconstruction and reconstruction performance itself changes with incidence. The sweep is capped at 60 degrees for that reason.

Scale-resolved roughness and curvature were evaluated at Gaussian widths from 0.1 to 1.6 mm. Texture descriptors were computed on band-passed relief rather than on the raw height field, so that the wing tilt does not dominate: areal roughness, the dispersion of the surface normal, mean curvature, curvedness and the shape index. A radially averaged power spectrum of the band-passed relief was computed on the largest hole-free patch of the reconstruction under a Hann window.

### Ultraviolet and telecentric configurations

Both alternative configurations replace hardware modules while leaving the reconstruction pipeline unchanged, and both are reported in full elsewhere. The telecentric configuration substitutes bilateral telecentric optics for the endocentric lenses, giving a measurement volume of 11 by 11 by 6 mm³ at 8.0 µm lateral and 4.46 µm axial resolution^5^. The deep-ultraviolet configuration replaces the projector with coherent speckle illumination at 266 nm and the visible optics with ultraviolet-transmitting equivalents^30^.

### Surface rendering

Surfaces are displayed as rendered surfaces rather than scatter plots. Points are projected orthographically from a chosen viewpoint onto a regular grid, the point nearest the camera is retained in each cell so that occlusion is handled correctly, surface normals are computed from the resulting depth field, and the surface is shaded by a single directional light with an ambient term. Scalar quantities are draped over that shading. Where a fine relief is displayed, points are averaged per cell rather than depth-buffered, because the cell size is comparable to the point spacing, and the relief is taken as the difference between two normalized-convolution smoothing that bracket the structure of interest. Specimens are reconstructed with the camera looking along the optical axis, which places them inverted relative to their true orientation; every rendering therefore flips the vertical image axis. Views rotated more than 30 degrees from the acquisition direction expose surfaces that a single view does not measure and are not used. Distributions of viewing directions on the sphere are drawn in a Lambert azimuthal equal-area projection, with radius proportional to the square root of one minus the absolute cosine of the polar angle, so that equal solid angles occupy equal areas on the page.

### Statistics and reproducibility

No statistical hypothesis tests are reported. Sample sizes are stated in every figure legend and are not the result of a power calculation: the planar reference is one artefact across 48 single acquisitions, the rotational reconstruction one specimen in seven views, the leaf one organ measured in two states, the lepidopteran one specimen in a single view, and the murine cohort four animals in two genotypes with two to five usable views each. No randomization or blinding was applied, and none would have been meaningful at these sample sizes. Where a range is quoted it is the range observed across the stated views or states, and where a mean and standard deviation are quoted the number of contributing measurements is given. Robust statistics are used wherever a distribution is contaminated by a minority population of gross errors and are identified as such at each use.

## Data availability

The reconstructed surfaces supporting this study are available at Zenodo under https://doi.org/10.5281/zenodo.22167250.54

Source data for all graph panels are provided with this paper; for panels showing rendered surfaces, the underlying reconstructions are in the same record.

## Code availability

The analysis notebooks, environment specifications, derived data and per-panel source data are available at Zenodo under https://doi.org/10.5281/zenodo.22167598.55

The reconstruction software, build documentation and minimal working examples are available at https://doi.org/10.5281/zenodo.22167471.56

The software and analysis notebooks are released under the MIT licence and the hardware design files under CERN-OHL-P v2. The visible-light platform described here is released without patent restriction; the deep-ultraviolet configuration is the subject of German patent application DE 10 2021 001 366 A1, held by Friedrich Schiller University Jena, and is described separately in ref. 35.

## Acknowledgements

C.F. thanks all Franke Lab members for support. The authors would like to thank Silvia Ruthardt in particular for organizational, administrative and moral support.

This work was supported by the German Federal Ministry of Research, Technology and Space (BMFTR) within the Research Program Quantum Systems, programme 3DVens, project number 13N16890, by the Thüringer Aufbaubank (TAB, FGR 0060 KI-supER), and by the Deutsche Forschungsgemeinschaft (DFG) via the Collaborative Research Centre PolyTarget (CRC 1278, project number 316213987, project B07) and HU 800/14-2.

G.J.G., A.W.S. and C.F. thank the K1 Start-Up Service of the Friedrich-Schiller University for supporting the Gentschinator3000 project.

We thank the students and early users who assembled, operated and broke the first *G3000* prototypes, whose questions and failures shaped the documentation and the software more than any of our own testing did.

## Author contributions

G.J.G., A.W.S. and C.F. conceived the project. C.F. supervised the project. C.F. and C.A.H. acquired funding. A.W.S. established the structured-light instrument design on which the platform is based and built the telecentric configuration reported here. G.J.G. designed and built the *G^3000^*, produced the hardware documentation and assembly guide, and performed the majority of the data acquisition, with contributions from M.G., A.P. and A.W.S.. G.J.G. prepared and positioned the animals for scanning. M.G. developed and wrote the reconstruction software. A.P. wrote the pattern generator. C.F. and G.J.G. developed the analysis pipelines and wrote the analysis notebooks. G.B. provided the lepidopteran specimen and contributed to data analysis and interpretation. C.A.H. and J.C.H. provided the murine specimens under their institutional animal license and contributed to data analysis and interpretation. G.J.G. and C.F. prepared the figures and wrote the manuscript, with contributions from all authors. All authors read and approved the final version.

