## Supplementary Information for "Democratizing three-dimensional surface phenotyping: an open structured-light platform reveals and removes the projection bias in biological imaging"

#### Contents

**Supplementary Note 1** Build guide: parts, assembly, wiring and firmware

**Supplementary Note 2** Software installation and verification

**Supplementary Note 3** Calibration walkthrough and diagnostics

**Supplementary Note 4** Your first measurement and reconstruction

**Supplementary Note 5** Origin of the periodic modulation in the plane residual

**Supplementary Note 6** Known limitations and when the measurement should not be trusted

**Supplementary Note 7** Troubleshooting

**Supplementary Note 8** Frequently asked questions

**Supplementary Note 9** Analysis notebooks and figure-generation scripts

**Supplementary Note 10** Surface classification on the murine specimens

**Supplementary Note 11** The nature of the murine demonstration, its limitations and what a designed study would require

**Supplementary Note 12** The scale at which a surface area is defined

**Supplementary Tables 1 to 7**

**Supplementary Figures 1 to 9**

**Supplementary Data 1 to 4**

### Supplementary Notes 1 to 4

#### Supplementary Note 1: Build guide

This note is the build manual. It assumes access to a fused-filament printer, a metric hex key set and a soldering iron, and no prior experience of assembling an optical instrument. Printed parts are supplied as STL and STEP files and are dimensioned for standard metric fasteners and bearings, so no drilling, tapping or reaming is required. The complete parts list is **Supplementary Table 1**. Assembly photographs are shown in line in chronological order as virtual panels of **Supplementary Fig. 9**.

The build described here is not limited to the original development setup. At the time of writing, five physical implementations exist: three base $G^{3000}$ systems, one deep-ultraviolet configuration and one telecentric configuration. During development, base systems were assembled and operated by undergraduate students and by one high-school student with only limited supervision, and one base system is permanently installed in the laboratory of our clinical collaborators and was used for the murine measurements reported in the main text.

#### 1.1 Camera assembly

1. Mount a post in its post holder and fasten the holder to the optical table with an M6 screw and washer, either directly or through a clamp.

2. Place an M4 nut in the printed camera mount and fix the mount to the post (**Supplementary Fig. 9a**).

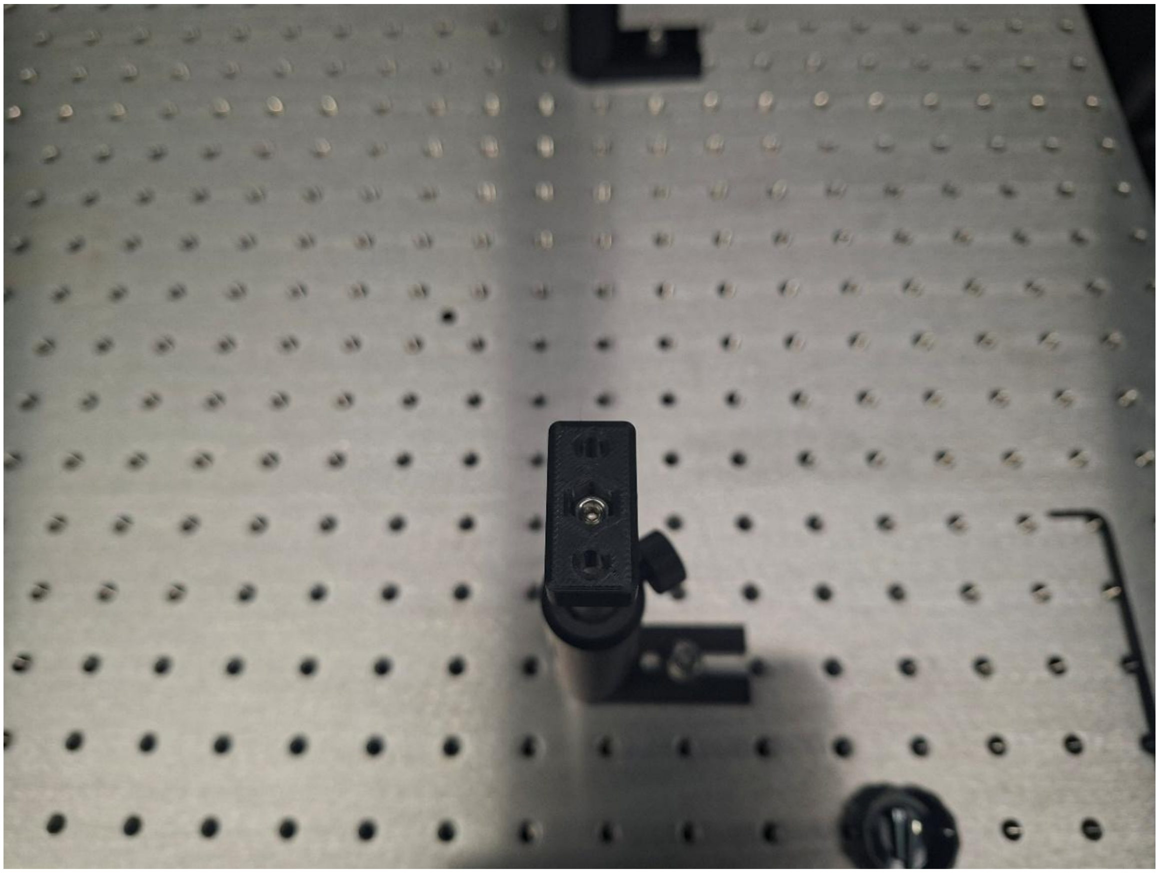

3. Secure the camera to the mount with two M3 screws (**Supplementary Fig. 9b**).

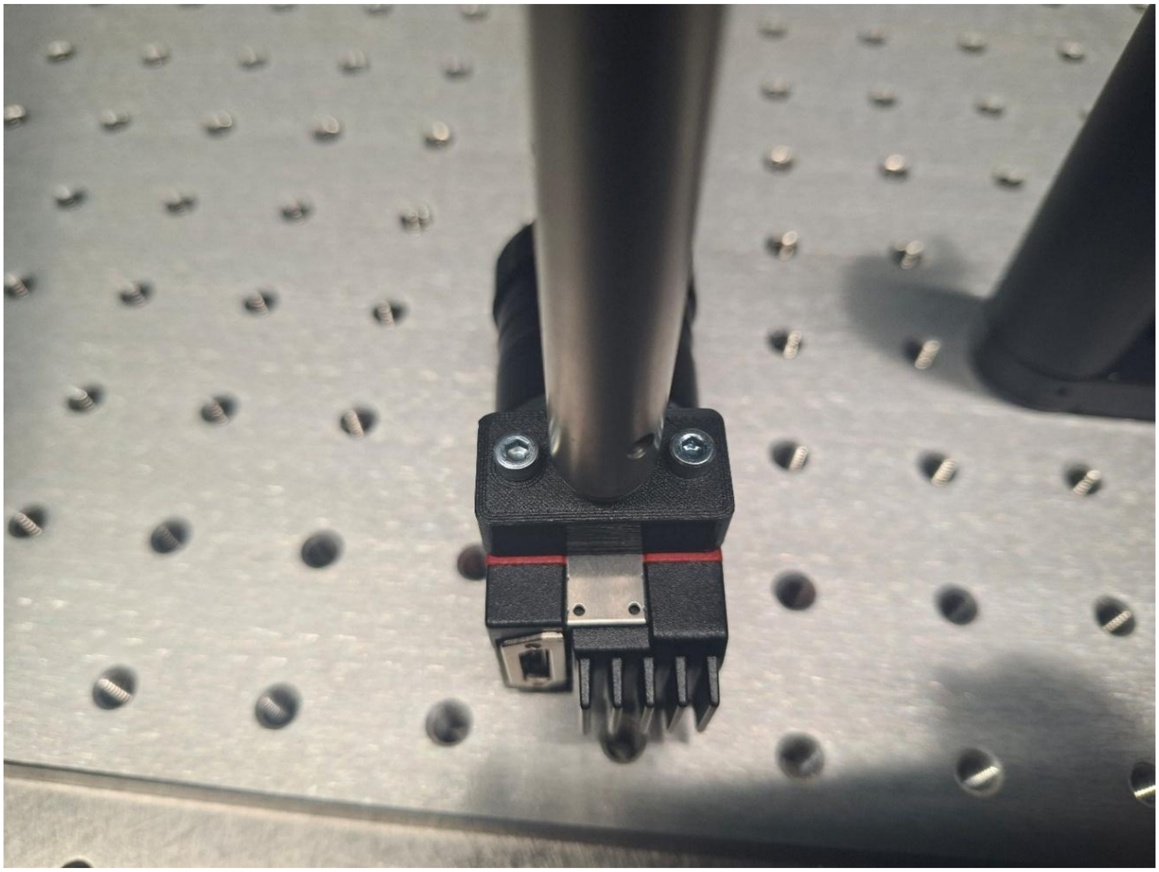

4. Connect the camera to the computer with a USB Micro-B cable rated USB 3.0 or higher (**Supplementary Fig. 9c**).

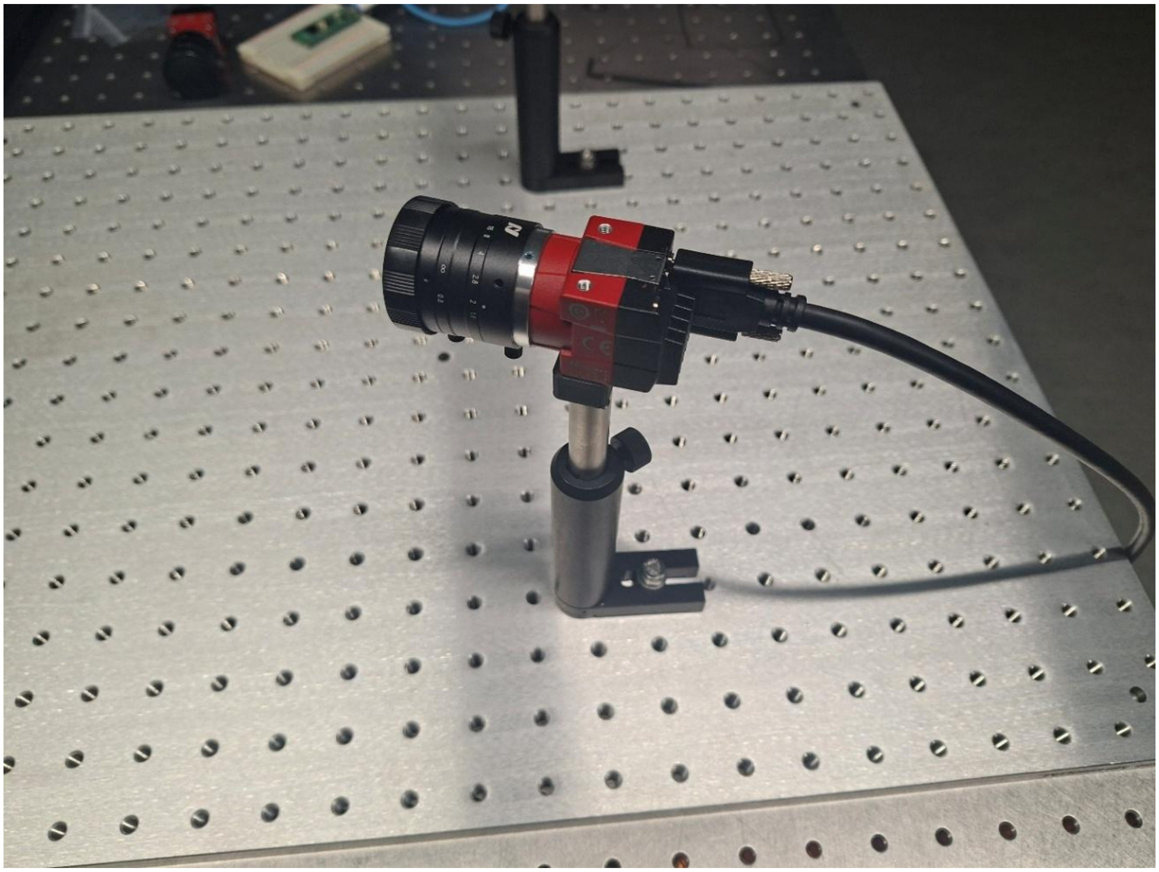

5. Remove the lens cap. This is optional for the build but necessary for calibration and measurement.

6. Repeat for the second camera.

##### 1.2 Turntable assembly

1. Fasten the printed base to the table with two M6 by 12 mm screws and washers.

2. Place the lazy-Susan ball bearing on the base, set four M4 by 6 mm screws in the corners and press down on them to fix the bearing. Driving each screw into the printed material with a hex key or screwdriver gives a more secure connection (**Supplementary Fig. 9d**).

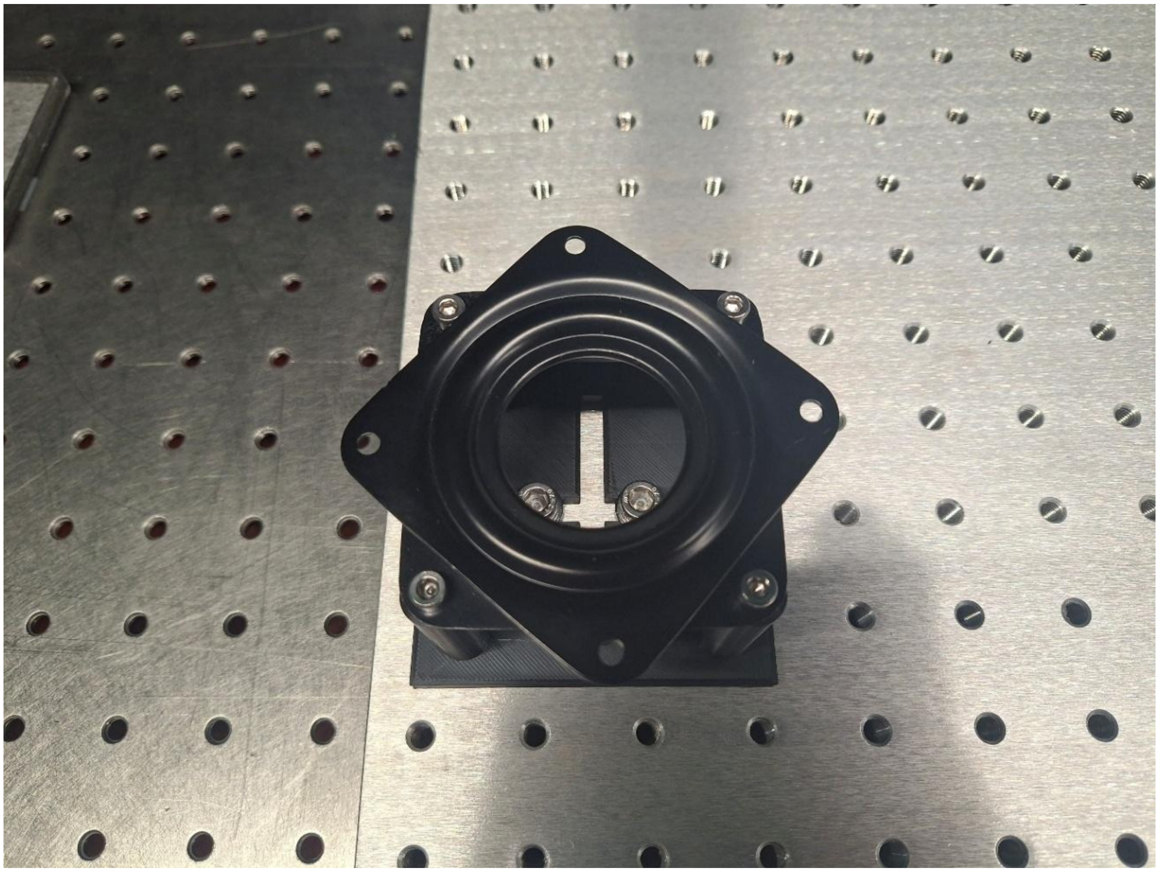

3. Stack the large gear on the ball bearing. It should slide into the inset on the gear.

4. Fit the motor cover to the stepper motor, orient it so that the distance between the motor shaft and the large gear is minimal, and fix it to the table with M6 screws.

5. Push the small gear down the motor shaft until it lines up with the large gear, checking that the teeth of the two mesh rather than collide.

6. Confirm that the gears are in contact and that the motor can drive the large gear (**Supplementary Fig. 9e**).

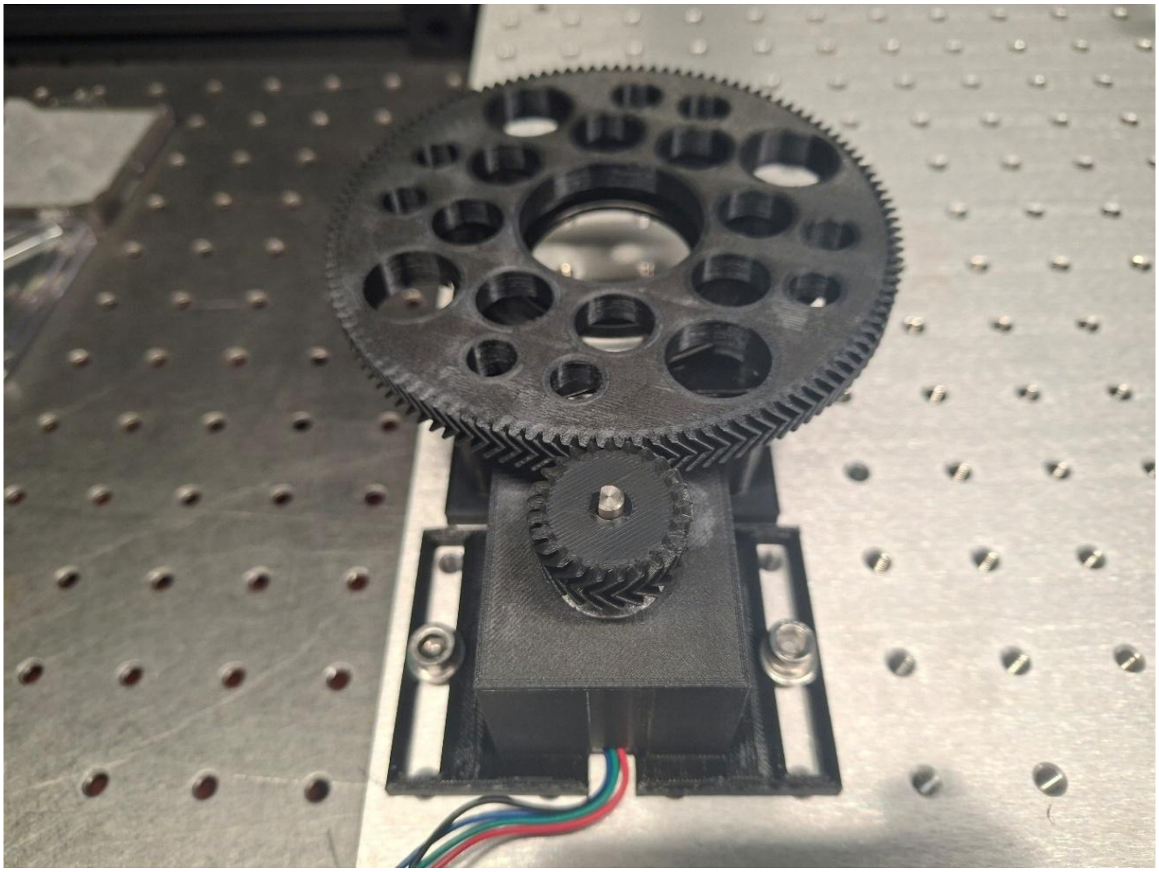

7. Place the Raspberry Pi Pico on the breadboard and connect its USB cable.

8. Connect jumper cables to pins 3 and 5. Cable colors are arbitrary and carry no meaning (**Supplementary Fig. 9f, g**).

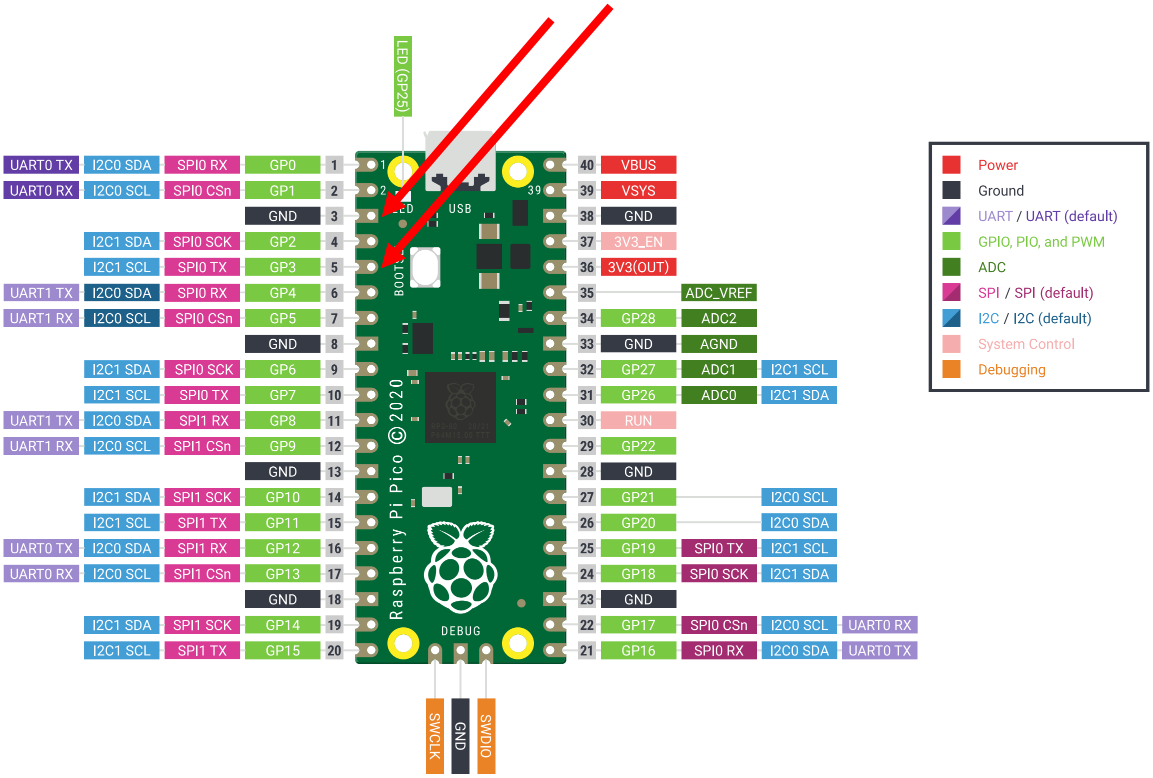

LED (GP25)
USB
BOOTSEL
Raspberry Pi Pico © 2020
DEBUG
UART0 TX | I2C0 SDA | SPI0 RX | GP0 | 1
UART0 RX | I2C0 SCL | SPI0 CSn | GP1 | 2
GND | 3
I2C1 SDA | SPI0 SCK | GP2 | 4
I2C1 SCL | SPI0 TX | GP3 | 5
UART1 TX | I2C0 SDA | SPI0 RX | GP4 | 6
UART1 RX | I2C0 SCL | SPI0 CSn | GP5 | 7
GND | 8
I2C1 SDA | SPI0 SCK | GP6 | 9
I2C1 SCL | SPI0 TX | GP7 | 10
UART1 TX | I2C0 SDA | SPI1 RX | GP8 | 11
UART1 RX | I2C0 SCL | SPI1 CSn | GP9 | 12
GND | 13
I2C1 SDA | SPI1 SCK | GP10 | 14
I2C1 SCL | SPI1 TX | GP11 | 15
UART0 TX | I2C0 SDA | SPI1 RX | GP12 | 16
UART0 RX | I2C0 SCL | SPI1 CSn | GP13 | 17
GND | 18
I2C1 SDA | SPI1 SCK | GP14 | 19
I2C1 SCL | SPI1 TX | GP15 | 20
40 | VBUS
39 | VSYS
38 | GND
37 | 3V3_EN
36 | 3V3(OUT)
35 | ADC_VREF
34 | GP28 | ADC2
33 | GND | AGND
32 | GP27 | ADC1 | I2C1 SCL
31 | GP26 | ADC0 | I2C1 SDA
30 | RUN
29 | GP22
28 | GND
27 | GP21 | I2C0 SCL
26 | GP20 | I2C0 SDA
25 | GP19 | SPI0 TX | I2C1 SCL
24 | GP18 | SPI0 SCK | I2C1 SDA
23 | GND
22 | GP17 | SPI0 CSn | I2C0 SCL | UART0 RX
21 | GP16 | SPI0 RX | I2C0 SDA | UART0 TX
SWCLK
GND
SWDIO
Power
Ground
UART / UART (default)
GPIO, PIO, and PWM
ADC
SPI / SPI (default)
I2C / I2C (default)
System Control
Debugging


97

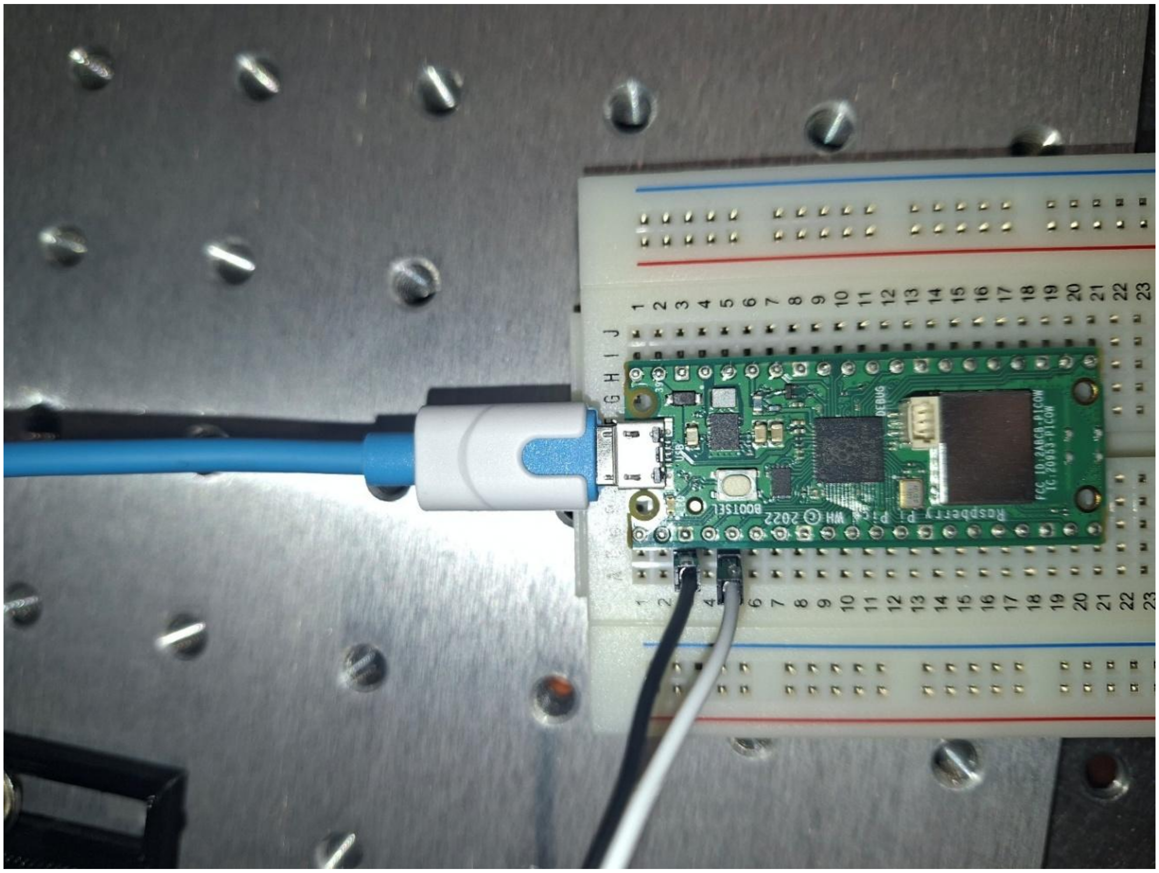

98

9. Connect pin 5 of the Pico to the PUL+ input of the driver and pin 3 to the PUL− input and fasten both (**Supplementary Fig. 9h**).

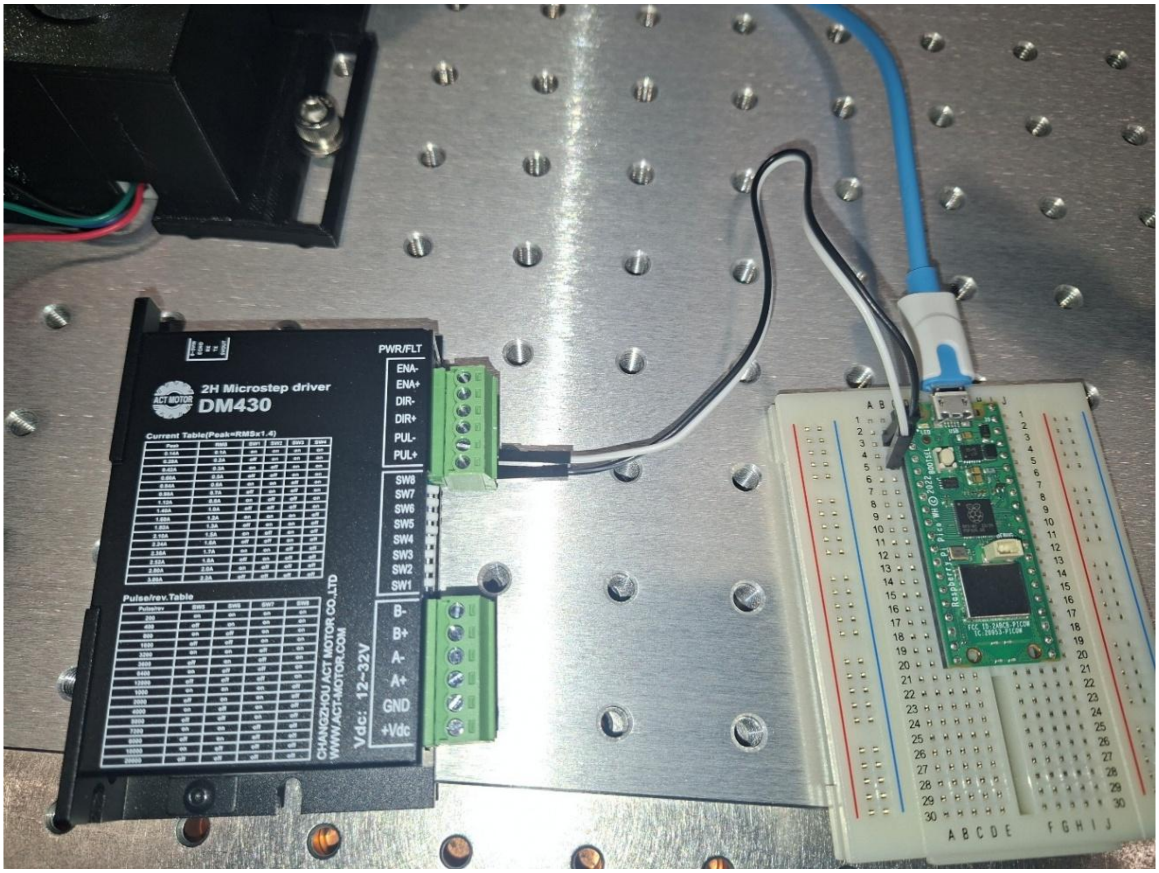


10. Connect the motor windings to the corresponding driver inputs. Consult the documentation for the motor; for the motor used here the mapping is black to A+, green to A−, red to B+ and blue to B−. Motor documentation is occasionally wrong, in which case the correct configuration has to be found by trial, for which `turntable_test.py` is used at step 15 (**Supplementary Fig. 9i**).

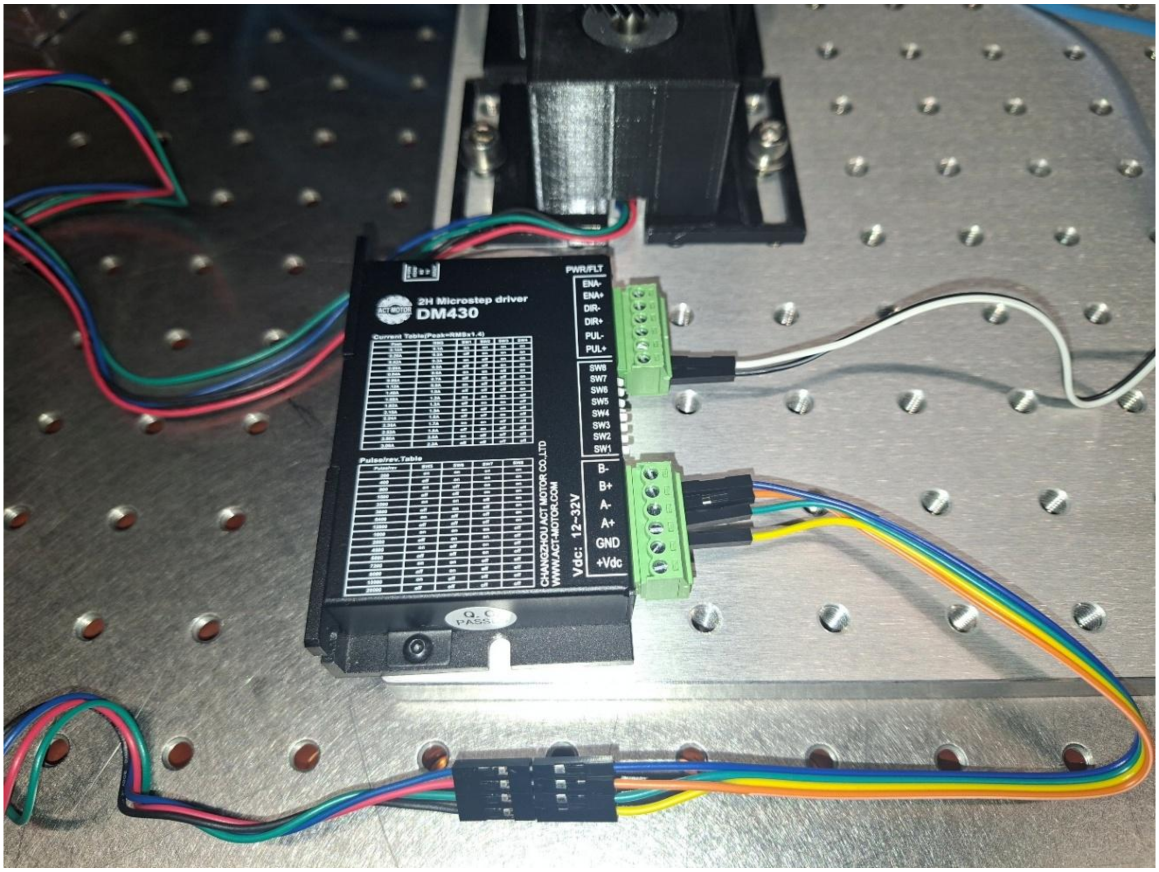


11. Connect the power supply adapter to the driver with jumper cables, positive to +Vdc and negative to GND.

12. Set the switches on the driver so that 1600 pulses correspond to one motor revolution, and so that the motor receives the appropriate voltage and current. Refer to the motor documentation and the printing on the driver (**Supplementary Fig. 9j, k**).

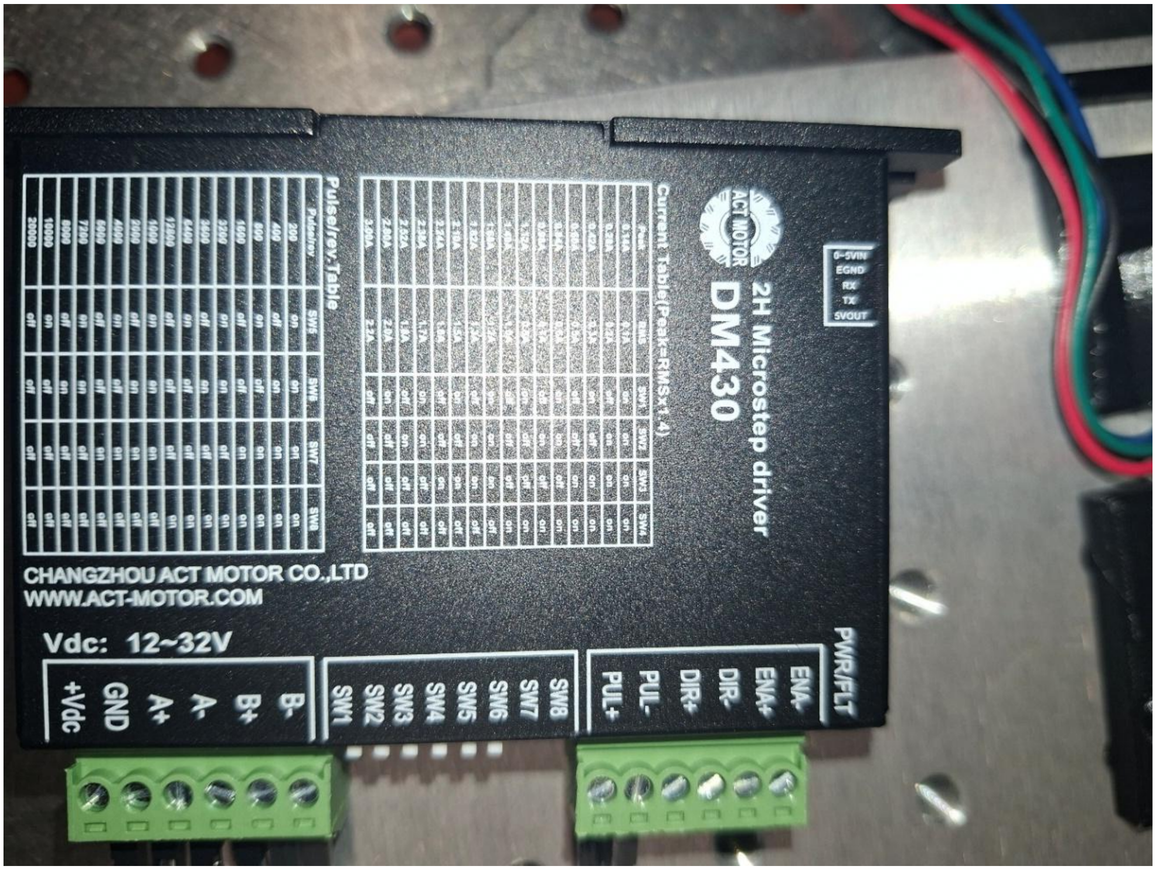
2H Microstep driver
DM430
ACT MOTOR
Current Table(Peak=RMSx1.4)
Pulse/rev Table
CHANGZHOU ACT MOTOR CO.,LTD
WWW.ACT-MOTOR.COM
Vdc: 12~32V
+Vdc
GND
A+
A-
B+
B-
SW1
SW2
SW3
SW4
SW5
SW6
SW7
SW8
PUL+
PUL-
DIR+
DIR-
ENA+
ENA-
PWR/FLT

113

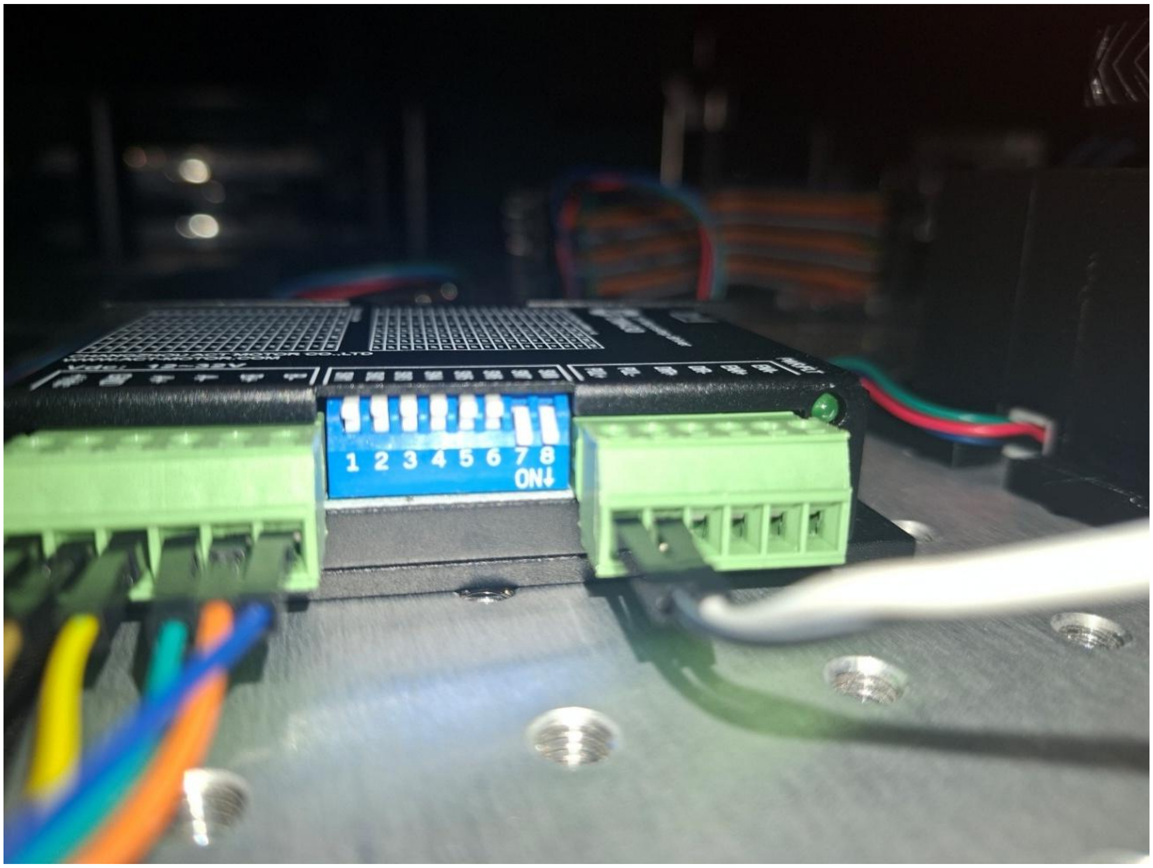


13. Hold the BOOTSEL button on the Pico while connecting it to the computer. A new device should be detected, and a window should open containing two files (**Supplementary Fig. 9l**).

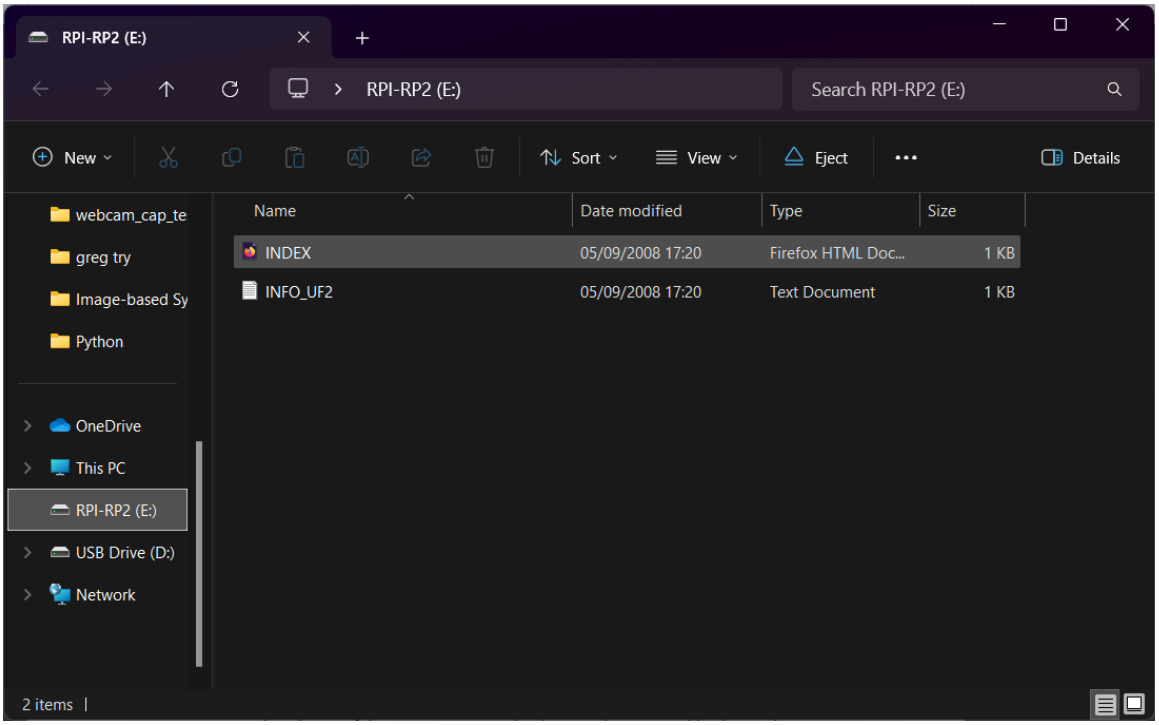

14. Download the Telemetrix firmware for the Pico and place the `.uf2` file in the RPI-RP2 folder. The folder and the device should then disappear. 15. Connect the power supply to the driver and run `turntable_test.py`. The turntable should rotate if everything is configured correctly (**Supplementary Fig. 9m**).

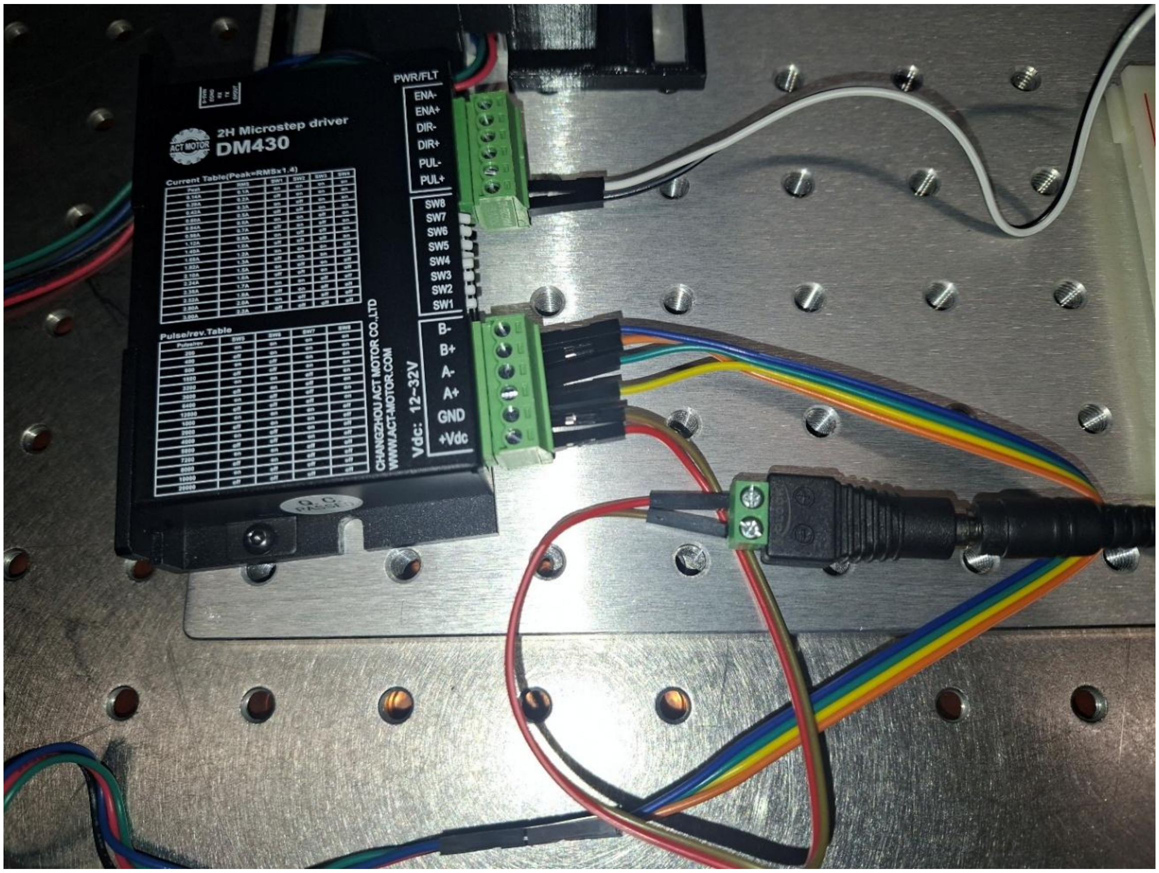


#### 1.3 Motion electronics and firmware

The stepper motor is driven through a two-phase microstep driver set to 1600 pulses per motor revolution, commanded from the Raspberry Pi Pico running the Telemetrix firmware. The printed 120-tooth gear is driven through a printed 25-tooth pinion, giving a gear ratio of 4.8, so the smallest commanded rotation is 0.047 degrees. That is two orders of magnitude below the increments used for rotational scanning, so the stage resolution is never the limiting term.

#### 1.4 Optical layout

The two cameras view a common volume in a convergent arrangement, with the projector between them. The layout is not fixed: baseline and working distance are chosen together to set the field of view, and the software requires no change when they do. Two arrangements have been used in practice and either can be rebuilt from the parts list.

The wide-field arrangement places the cameras on a 705 mm baseline at a working distance of 920 mm, giving a field of view of approximately 170 by 110 mm and a convergence angle of 43.8 degrees between the optical axes. This is the configuration characterised in the main text and used for the rotational and murine measurements.

The compact arrangement shortens the baseline to 365 mm at a working distance of 315 mm, giving a field of view of approximately 85 by 80 mm. It suits specimens that fit within that field and needs considerably less bench space, at the cost of a shallower measurement volume.

Position the cameras at roughly equal distances from the centre of the turntable, leaving open space between them for the projected pattern to pass through. Measuring and recording the distance from the turntable centre to the camera lens makes the arrangement easier to rebuild after it has been disturbed. If space is limited, a lens in front of the projector shortens its throw; a 500 mm lens was used for one such arrangement.

##### 1.5 First-light checklist

Before the first calibration, confirm that both cameras are detected by the capture program, that the projected pattern is sharp in the plane through the center of the turntable, that both camera views overlap and are centered on that plane, that the turntable rotates under `turntable_test.py`, and that the focus dials on both objectives are fixed. A calibration is valid only for the exact optical configuration in which it was acquired.

#### Supplementary Note 2: Software installation and verification

The toolbox targets Python 3.12.13 and is installed from the supplied environment specification. Using the command “conda create --file environment.yml“ and the provided environment all required packages apart from the AVT camera SDK can be installed. Tested on Windows 11 25H2.

A small, pre-recorded example project is included with the distribution. Running the complete pipeline on it reproduces a reference point cloud, which allows an installation to be verified before any hardware is connected.

Two programs are used during acquisition and one during reconstruction. `live_capture_alvium_gui.py` acquires calibration images, `Gui_turntable_preview.py` runs a measurement, and `reconstructor.exe` performs the calibration and the reconstruction. Each is described where it is first needed, in **Supplementary Notes 3 and 4**.

#### Supplementary Note 3: Calibration walkthrough and diagnostics

##### 3.1 Calibration setup

Place the turntable and cameras in their intended positions and fix them to the optical table. Place the projector and check that its projection is sharp in the plane through the center of the turntable. Then place a chequerboard of known square size in the field of view of both cameras, using `live_capture_alvium_gui.py` to see what they see.

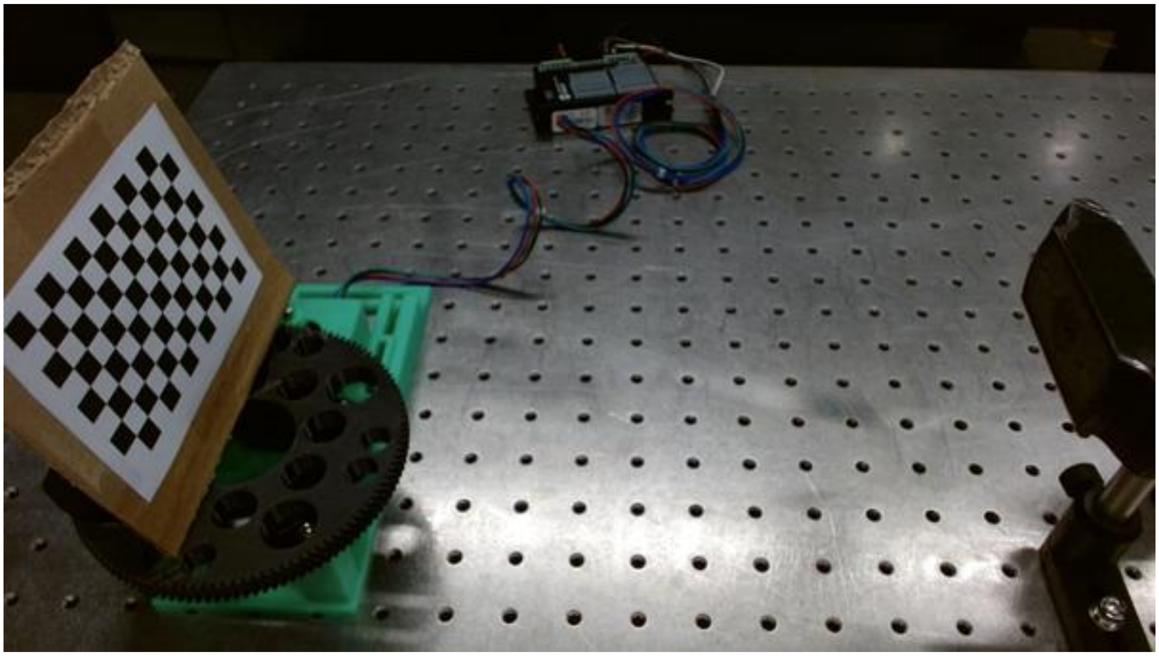

##### 3.2 The capture interface

`live_capture_alvium_gui.py` presents four controls.

| Control | Function |
|---|---|
| Output folder | Location of saved images. Press the three dots to define it. |
| Exposure time | Exposure of both cameras, set either in the field or with the slider below it. |
| Start capture | Opens the camera preview. |
| Preview window | Press the space bar to save an image pair and Q to stop. The frame counter starts at zero when Start capture is pressed. |

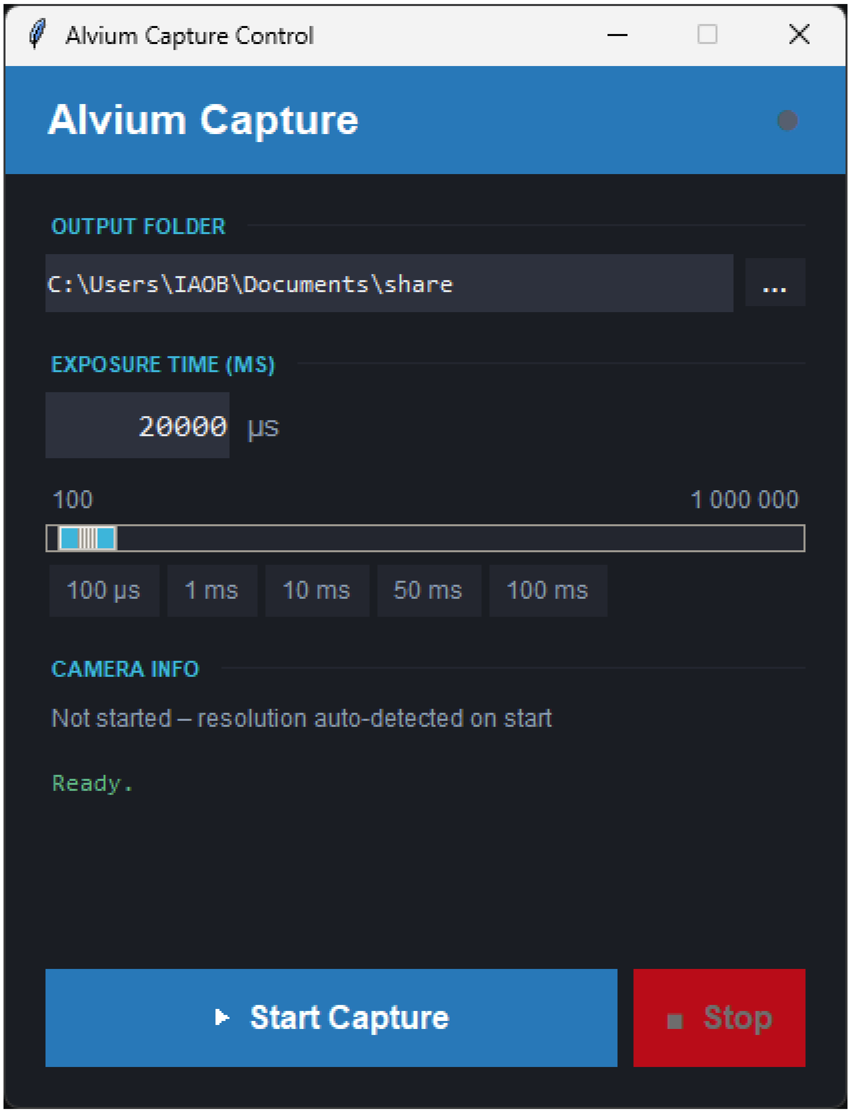

#### 3.3 Acquiring calibration images

1. Open the program and define the output folder.

2. Press Start capture to open the preview.

3. Place the chequerboard in the field of view and adjust the focus of both cameras, usually with the dial on the objective. Adjust the posts so that the two views overlap and are centered. The center hole of the large gear is a convenient reference for both placing the chequerboard and aligning the cameras. Check that the edges of the chequerboard are sharp and that the image is bright enough for the pattern to be made out.

4. When the focus and alignment are satisfactory, fix the dials on both objectives. The calibration that follows is valid only for this exact configuration.

5. Press the space bar once as a test. Confirm in the file browser that two images were saved, that the frame counter advanced, and that the saved images match the preview in sharpness and brightness.

6. Move the chequerboard through the field of view, covering it as fully as possible and paying particular attention to the edges. The chequerboard must be fully visible in both cameras in every image and should be tilted at least slightly with respect to them. More images generally improve the calibration; at least 30 are advised.

7. Repeat until the field of view has been covered.

#### 3.4 Image quality, what to avoid and what to aim for

Two failures account for most poor calibrations: an image too dark for the chequerboard corners to be detected, and an image in which the chequerboard is out of focus, so that corner positions are recovered imprecisely or not at all. Both are visible in the preview before any image is saved.

**INCORRECT!** Unsharp Image!

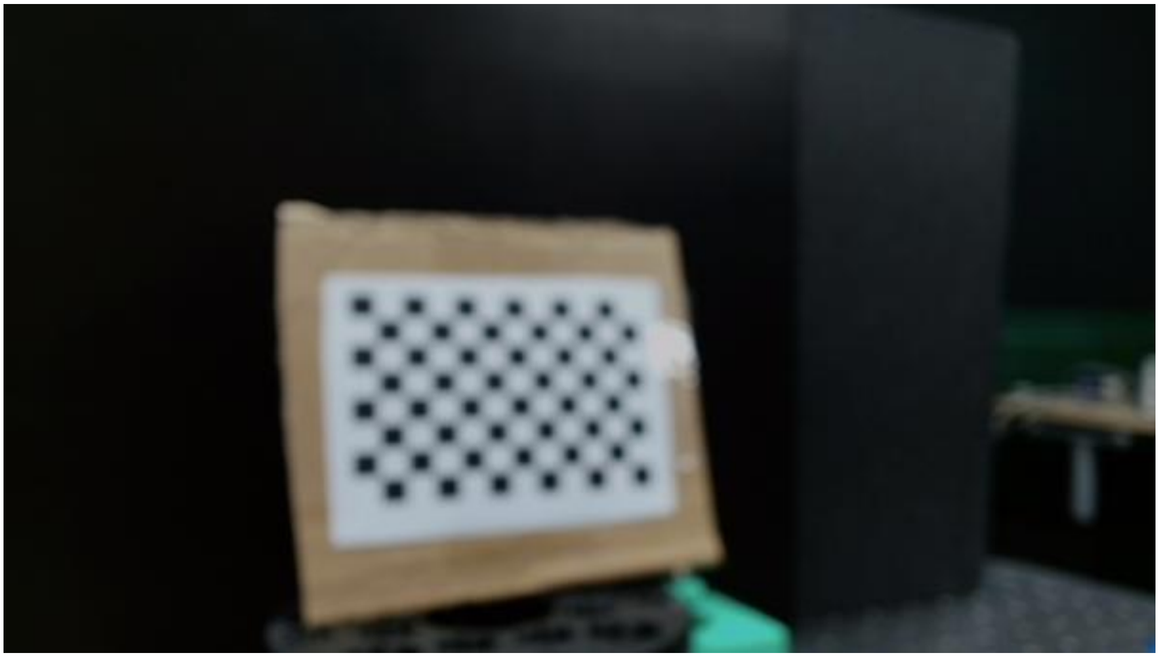

**INCORRECT!** Image too dark!

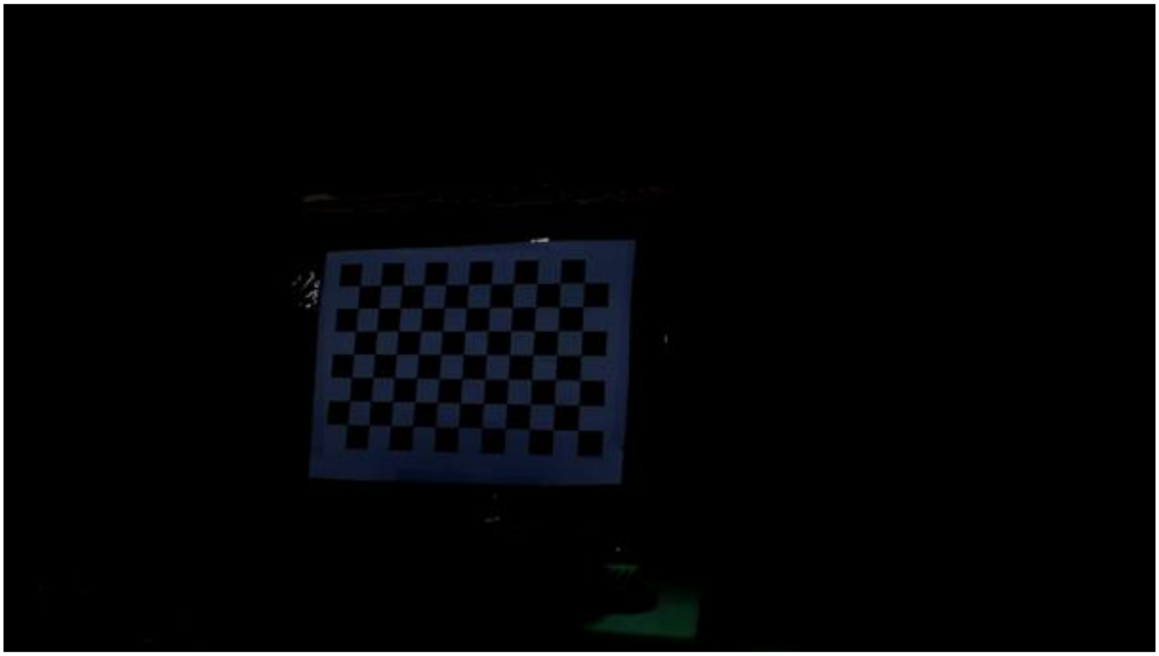

**CORRECT! Correct focus and brightness results in a *sharp* and *bright* image!**

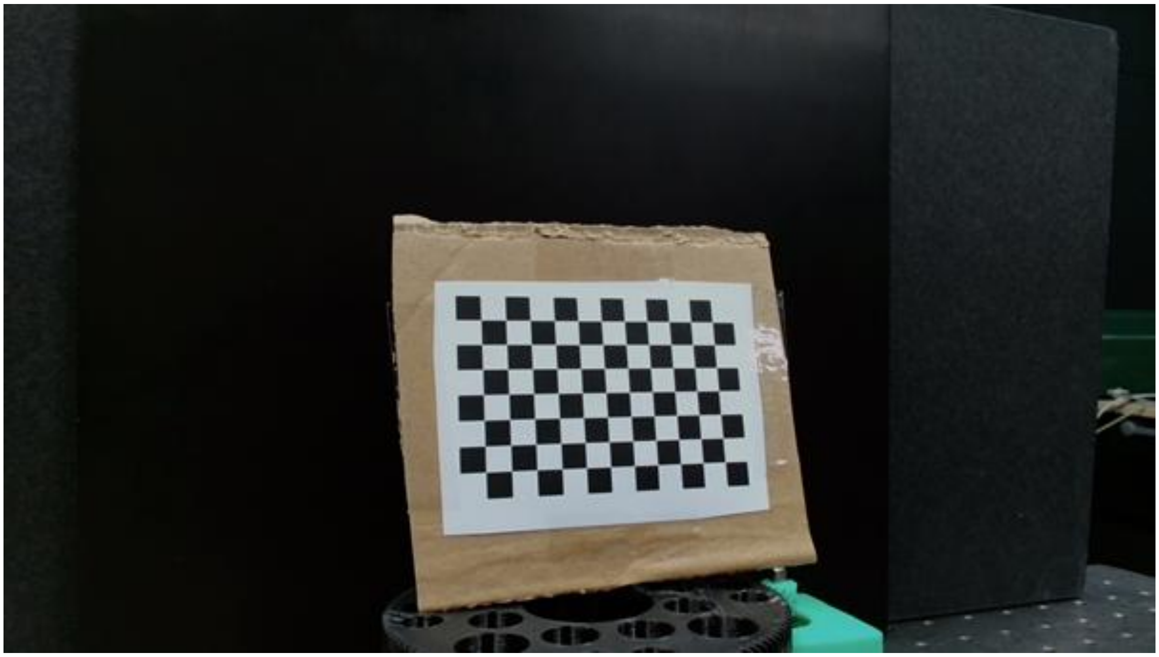

##### 3.5 Running the calibration

Open `reconstructor.exe`. The program may take some time to load. Press the camera calibration button, which opens a small dialog.

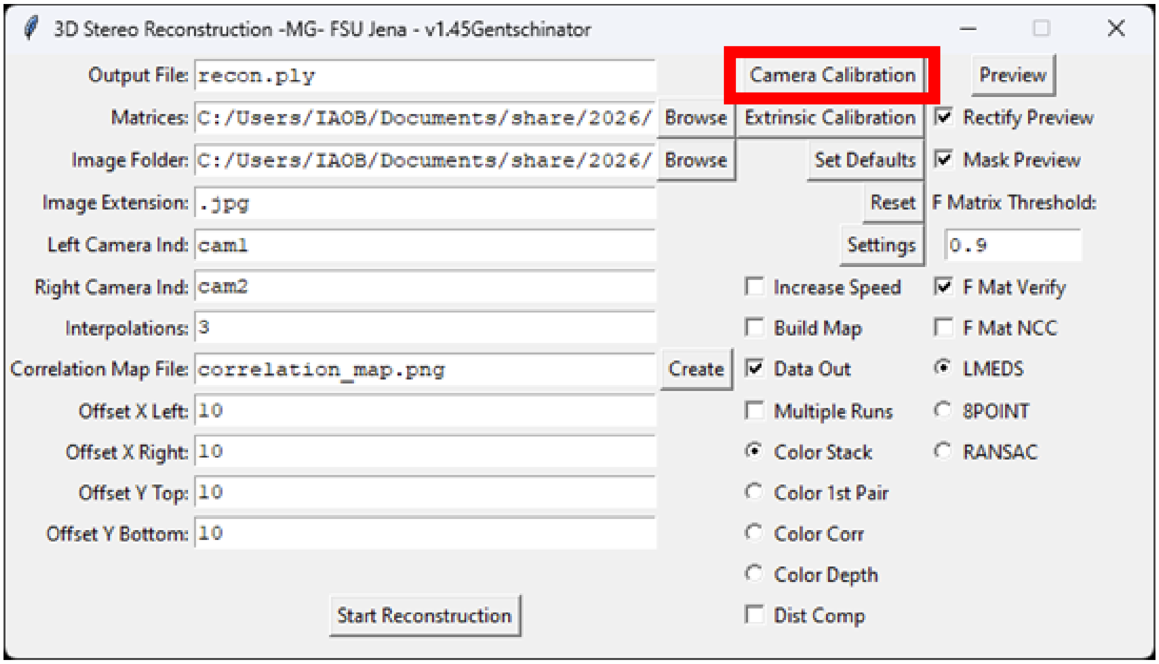

Browse to the folder of calibration images and select it; the folder may appear empty in the file browser, which is expected. The left and right indicators identify the images from each camera and can be left at their defaults. The result folder defines where the calibration matrices are written, conveniently a subfolder of the calibration images.

Rows, columns and square size describe the chequerboard used. Rows and columns are the numbers of interior corners, that is the crossings between squares, in each direction, and not the numbers of squares. The size is the edge length of one square in meters.

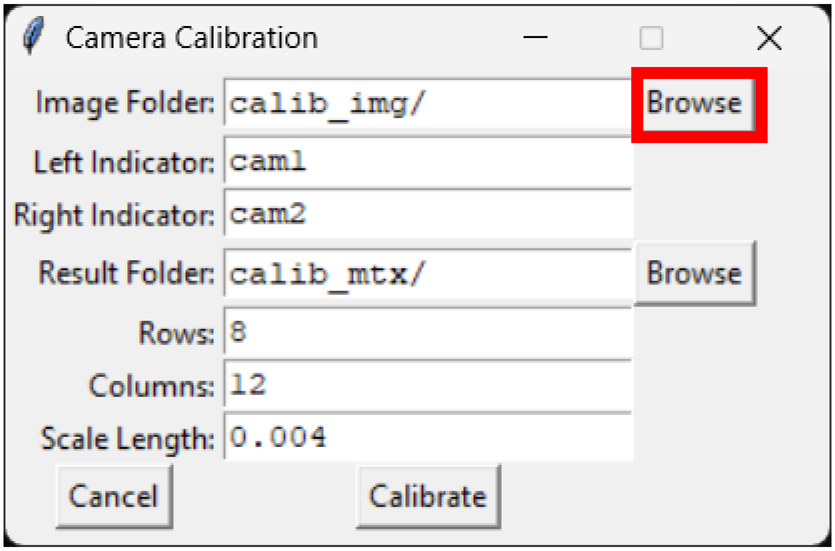


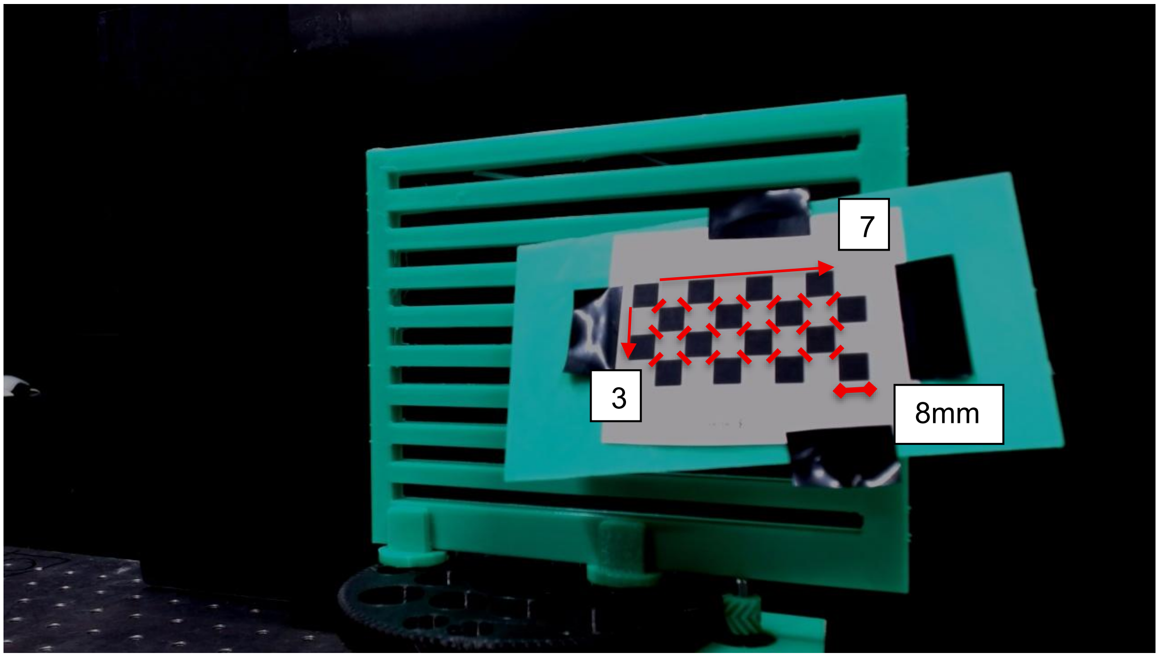


With the parameters set, press Calibrate and watch the second window that opened with the program. Progress bars should fill and the message "Calibration complete" should appear. Confirm that the matrices have been written to the result folder.

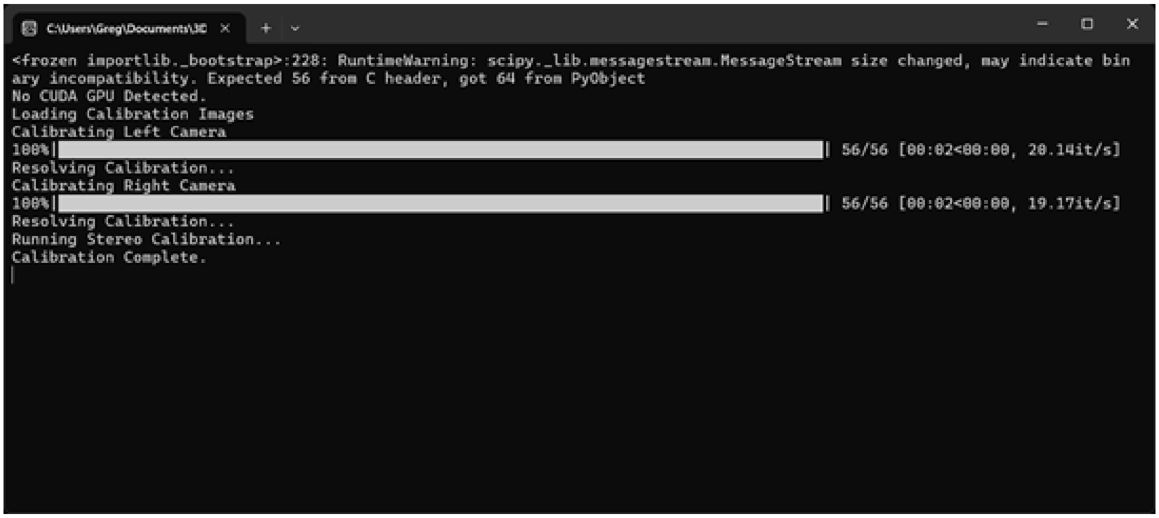


##### 3.6 When to recalibrate

A calibration remains valid until a camera, lens, focus setting or aperture is changed, or until either camera is moved relative to the other. Recalibrate after any of these, and after the instrument has been dismantled and rebuilt. The axis calibration used for rotational scanning is separate and remains valid until the stage is disturbed.

#### Supplementary Note 4: Your first measurement and reconstruction

##### 4.1 Pattern generation

Patterns for illumination are generated before the first measurement, using `pattern_generator.exe`.

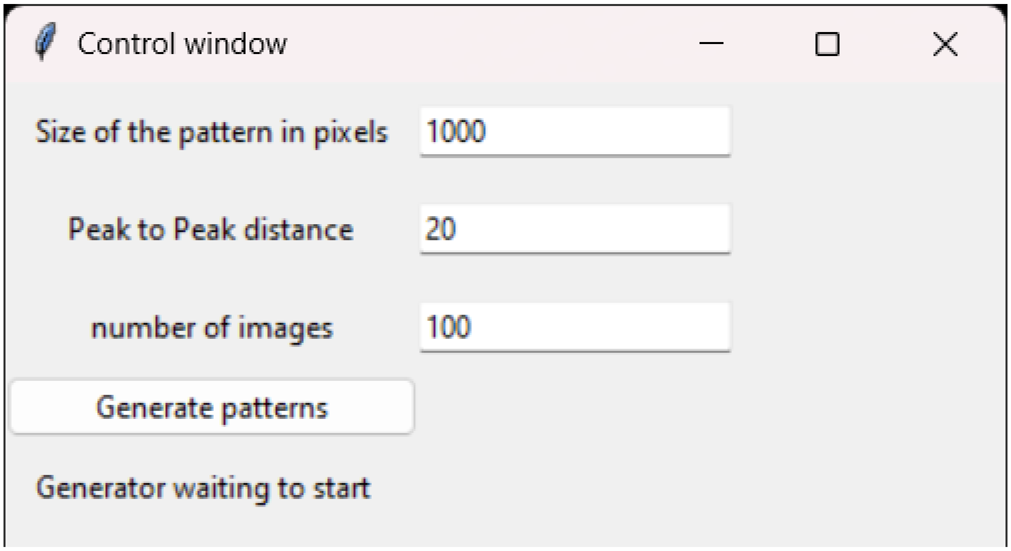

| | |
|---|---|
| Size of the pattern in pixels | How large a pattern you want to generate, since the pattern is square the maximum height resolution of your projector will limit it<br>(1000 is usually a good value if you are not sure) |
| Peak to Peak distance | How small or large the pattern will be, refer to the paper what influence this parameter has on your precision<br>(20 is a good value to start) |
| Number of images | How many images you want to generate in this go, refer to the paper what influence this parameter has on your precision<br>(for a first test 50 should be fine) |
| Generate patterns button | Starts the generator by prompting you to select a folder to put your pattern images into, we recommend making a new folder and writing down its directory path |
| Generator waiting to start | Label indicating if the generator is busy (“Generator busy”) or done (“Generator done”) |

After generation, add a uniform white image named `a_white` to the folder. The reconstructor uses it to compute per-point color.

#### 4.2 First test measurement

Use `Gui_turntable_preview.py`. For a first measurement only one view is needed, so the Pico must be connected to the computer, but the motor driver can be left unpowered.

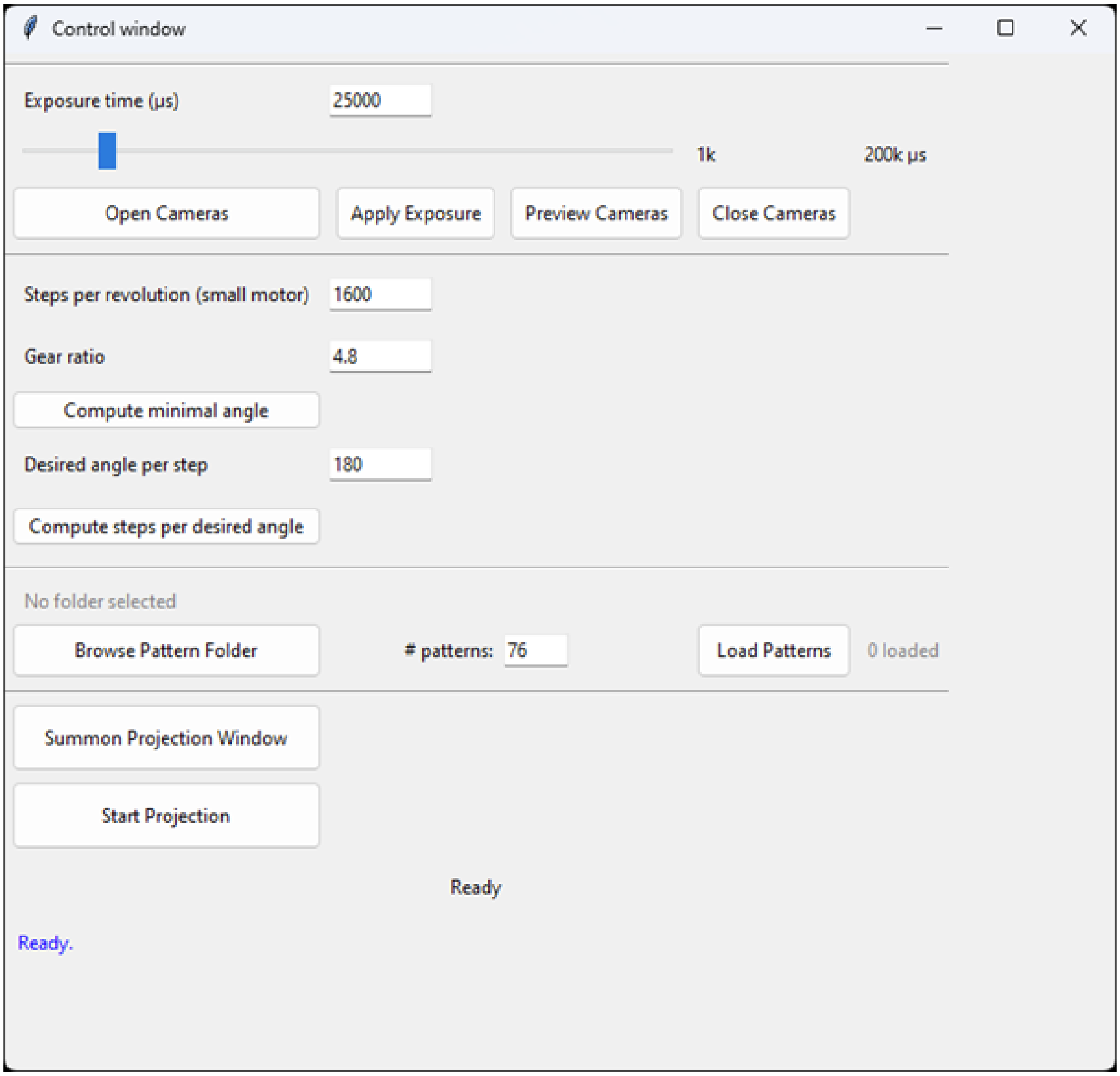


For this initial measurement we are only interested in one view of the object, you will still need to connect the raspberry pi pico to your computer but can leave the power source unconnected to the motor driver.

| | |
|---|---|
| Camera Control Panel | Directly define exposure time directly or use the slider (input values outside of the slider window directly)<br>First press “Open Cameras” to activate them for a preview or measurement, then “Apply Exposure” to use the direct input or use the slider.<br>“Preview Cameras” opens a semi-live preview window<br>“Close Cameras” severs connection to cameras in case of a required restart |
| Turntable Control Panel | Input the required steps for a full revolution (earlier in the setup we recommended 1600) and the gear ratio which is defined by:<br>For 120 and 25 teeth respectively, this should be 4.8<br>“Compute minimal angle” button is required to calculate the smallest angle the motor can turn with one step<br>Input the desired angle between single views<br>“Compute steps per desired angle” button is required to calculate the total number of steps required to turn the desired angle |
| Pattern loading Panel | “Browse Pattern Folder” lets you choose the location of your generated patterns from before, this is also how you can repeat measurements with different pattern sizes<br>Number of patterns is the full number of patterns in the folder by default – adapt this if you want to use less<br>“Load Patterns” button loads patterns and prepares them for projection |
| Projection Panel | “Summon Projection Window” button creates the window you move over to your projector to illuminate object to be measured (required)<br>“Start Projection” button lets you choose a folder to save your images, make a new folder so that only the images from your measurement are saved there, later the program will generate subfolders required for the reconstruction |
| Status Messages | Provide feedback to actions taken by the user and is used for monitoring |

##### 4.3 First test reconstruction

Open `reconstructor.exe` again. Only a subset of its controls is needed for a first reconstruction; the remainder are advanced options that can be left at their defaults.

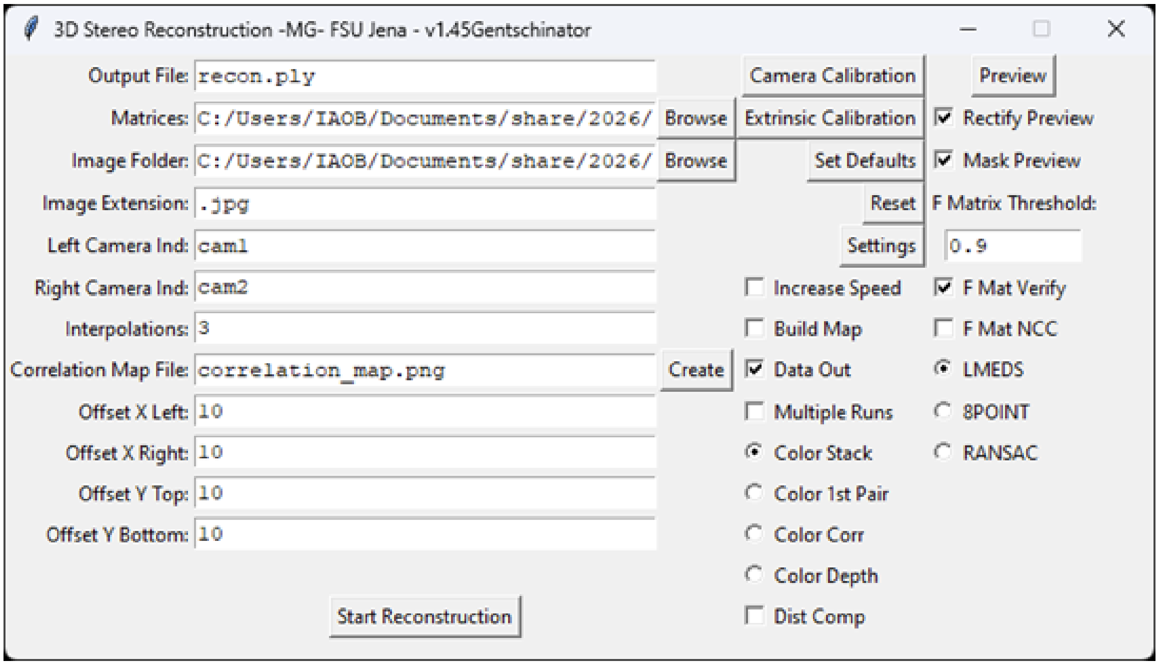


| | |
|---|---|
| Output file input | Input the desired name for your file, the ".ply" ending is important and must stay that way, make sure you don't put spaces behind it or press enter. |
| Matrices input | Put in the directory of the matrices you got from the calibration manually or use the browse button next to the input to select the folder where they are located |
| Image Folder | Location of your measurement, to check a single view navigate to one of the subfolders, for full reconstruction use the measurement folder |
| Image Extension, Left images and right images Ind | Can be ignored as they are defined by the Gui turntable program |
| Interpolations | Controls the number of subpixel interpolations made between each pixel - can be left at default value |

| | |
|---|---|
| Correlation Map File | Controls name and location of output correlation map, which is an image produced using the dimensions of the first left camera image, and filled in based on which points were successfully matched with corresponding points in the right camera images - can be left at default value |
| Offset X/Y left/right/top/bottom | Define the area in both camera images that is used for the correlation search – minimum value is 1 |
| Checkboxes<br><br><br><br><br>Checkboxes:<br>“F Mat Verify” , “F Mat NCC“ , “LMEDS” , “8POINT” and “RANSAC” are advanced options and can be left at default values | “Increase Speed” - use Speed Interval from Settings menu to do a quick reconstruction skipping the defined number of image columns for reconstruction<br>“Build Map” - additionally output a correlation map image at the same time as the point cloud .ply format output<br>“Data Out” - output a text file containing more information than the .ply file (useful for filtering)<br>“Multiple Runs” - will run the reconstruction program on a folder of image folders instead of a single image folder, with a separate reconstruction for each folder (used for 360° measurements)<br>“Color Stack” - computes the color for the 3D points as an average of the stack<br>“Color 1$^{st}$ Pair” - computes the color for the 3D points from the first image pair<br>“Color Corr” - use blue-green-yellow-red scale of correlation score (Jet Colormap)<br>“Color Depth” - use blue-green-yellow-red scale of distance from camera pair (Jet colormap)<br>“Dist Comp” - load distortion compensation arrays and modify the images accordingly to compensate (recommended) |

Before starting, tick Data out and Multiple runs, then open Settings and change the correlation threshold from 0.9 to 0.01. This deliberately permissive reconstruction threshold preserves almost the full correlation distribution for later gating rather than discarding low-correlation points at reconstruction. Application-specific thresholds are then applied during export or analysis, as listed in Supplementary Table 2 and in the corresponding Methods sections.

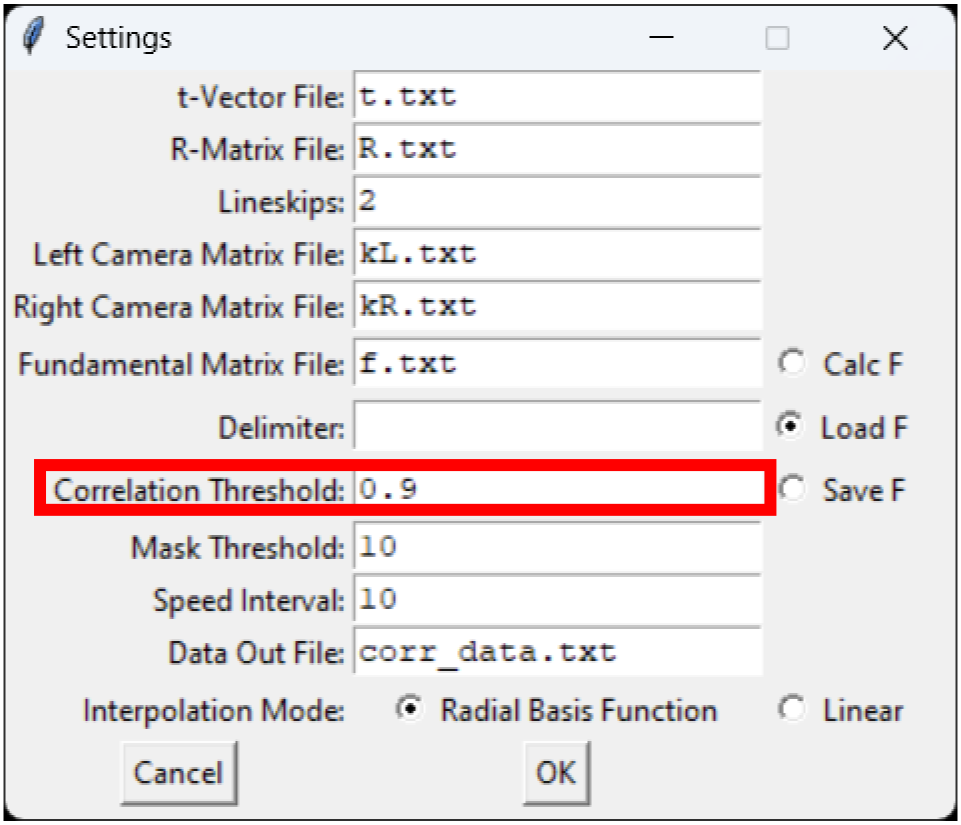


Press Preview. A window opens showing the reconstruction, which takes a few moments for a single view and considerably longer for a full measurement folder. The object should be readily recognizable in both images. Stretching, or an object that cannot be recognized, indicates a problem with the calibration or with the parameters entered during acquisition.

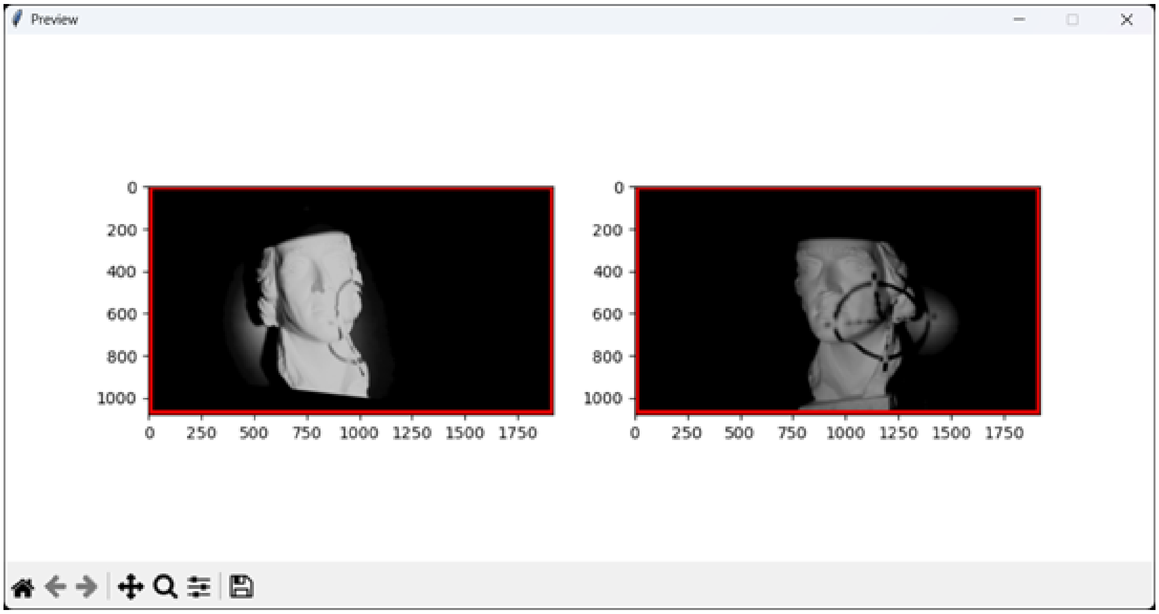


Close the preview and press Start reconstruction. A progress bar appears in the command window that opened alongside the program. When the message "Reconstruction complete" appears, the point cloud will be found under the name given, in the folder containing `reconstructor.exe`, together with a file named `corr_data.txt`.

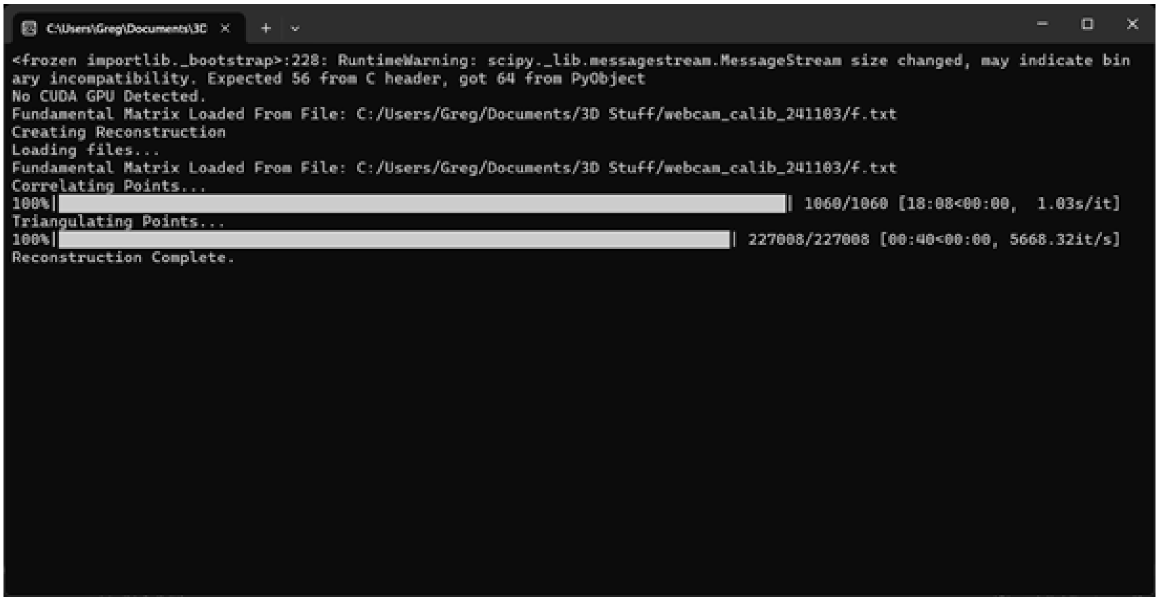


Both files can be opened in any point-cloud viewer. CloudCompare is free, open source and sufficient for inspection and for the manual segmentation used in this work.

#### Supplementary Note 5: Origin of the periodic modulation in the plane residual

The residual map of the certified plane retains a weak periodic modulation whose orientation follows the projection axis (**Fig. 3a,b**). We investigated this systematically, because periodic structure in a flatness measurement is easily mistaken for a defect of the artefact or for a temporal artefact of the projector.

The modulation is not caused by temporal aliasing between the digital micromirror device and the camera integration window. Varying the exposure time across the full range reported in **Fig. 3a** changes the amplitude of the modulation only in proportion to the overall residual and does not change its spatial period or phase, which would be the signature of a temporal beat. The modulation is instead strongest where the projected pattern is best focused and becomes smoother where the pattern is slightly defocused, which is the opposite of what improved contrast alone would predict.

The mechanism is an intensity-gradient bias in sub-pixel correlation. Where the pattern is sharply focused, the intensity gradient across a correlation window is large and asymmetric, and parabolic refinement of the correlation peak is biased towards the steeper flank. The bias reverses sign with the sign of the gradient, so a quasi-periodic intensity pattern produces a quasi-periodic depth error. Slight defocus lowers the gradient and symmetrizes the local intensity profile, suppressing the bias at the cost of a modest loss of contrast.

Two mitigations are available and both are implemented as options. Slightly defocusing the projection reduces the modulation without a measurable penalty in residual standard deviation over the range tested. Increasing the number of independent pattern realizations averages the bias over different local gradient configurations, which is one reason the residual in **Fig. 3b** continues to improve slightly beyond the point at which the valid fraction has saturated.

#### Supplementary Note 6: Known limitations and when the measurement should not be trusted

##### 6.1 Physical limits

**Specular and transparent surfaces.** Triangulation requires backscattered texture. Polished metal, glass and water surfaces return specular highlights or nothing at all in a single spectral band. The deep-ultraviolet configuration extends the accessible class considerably but does not solve the general case.

**Volumetric scatterers.** Fur, feathers, hydrogels and optically cleared tissue return a depth somewhere within the medium rather than a surface, at a position that depends on viewing geometry. The measurement fails in a way that registration detects but cannot be repaired. The murine data in the main text quantify this case.

**Line of sight.** Cavities, deep undercuts and the supported underside of an object on a turntable are not observed. Reported surface areas are therefore lower bounds whenever a specimen occludes part of itself, as the desiccated leaf and the spread lepidopteran both do.

**Motion.** The instrument assumes a static or quasi-static specimen, because acquiring a pattern sequence takes longer than a moving animal will hold still.

**Traceability.** The system is characterized by a certified plane but is not a certified coordinate measuring machine. Applications requiring formal traceability need validation appropriate to that context.

**Brightness.** Depending on the reflectivity and color of the specimen or object enough light must be collected by the camera to enable the cross-correlation algorithm, too low signal will worsen the accuracy and completeness.

##### 6.2 Analysis limits

**Grazing incidence.** The foreshortening factor diverges as a surface turns edge on, so a small number of near-silhouette cells can contribute a large and unreliable share of a computed surface area. Any area or rugosity figure should be reported together with the incidence cap applied to it.

**Silhouettes on thin specimens.** At the outline of a specimen the numerical gradient reports the step from surface to background rather than a surface slope. Removing one lattice cell at the outline is usually sufficient; removing more erases genuinely thin structures such as antennae and legs.

**Fixed-percentile thresholds.** A descriptor defined by a percentile of its own distribution cannot differ between conditions. Absolute, physically stated thresholds should be used for any quantity intended to be compared between states.

**Fixed cell counts.** Rasterizing two specimens of different extents onto grids of the same cell count evaluates every scale-dependent descriptor at a different physical scale. A common physical grid spacing should be used instead.

**Principal-axis sign.** Principal axes fix the plane of a flat specimen, but not which side is up, and the sign returned by a singular value decomposition is arbitrary. Orientation should be fixed by a physical criterion, here by requiring that relief point upward.

##### 6.3 Removing outliers, when it is appropriate

No post hoc spatial filtering was applied to the reconstructions reported in the main text, if not stated otherwise, so that the reported measurement is the measurement. On densely sampled surfaces, isolated outlying points can be removed by statistical outlier removal, radius-based neighborhood filtering, confidence thresholding on the retained correlation coefficient, or exclusion of points acquired beyond a stated incidence angle. Any of these changes the reported surface statistics and should be stated when used. Under optimal conditions a correlation coefficient threshold of 0.9, or higher, can still result in a dense and complete point cloud. However, in most “real-world”, e.g. biological, measurement scenarios this can differ drastically and, in our experience, there is no “one solution for all”. We recommend making test measurements and “playing around” with different correlation coefficient thresholds and keeping one constant for more complex tasks, multiview measurements.

#### Supplementary Note 7: Troubleshooting

**Calibration does not converge, or the reprojection error is high.** The chequerboard backing is not flat, the board is not fully visible in both cameras, or too few images cover the field edges. Reprint the board on a rigid backing and acquire more images with the board near the corners and tilted about both axes.

**Reconstruction shows large holes on one side of the specimen only.** The projected pattern does not reach that side, or that side is occluded from one of the two cameras. Rotate the specimen or reposition the projector so that the illuminated cone covers the region seen by both cameras.

**The valid fraction is low everywhere.** The exposure is clipping, the pattern feature size is outside the usable range, or too few patterns were projected. Check the histogram for saturation first, then confirm that the feature size lies between 20 and 100 projector pixels and that at least 40 patterns were used.

**The point cloud shows a stepped or terraced surface.** Sub-pixel interpolation is disabled, or the correlation window is too small for the projected feature size. Restore the default of three interpolation steps and increase the feature size.

**A visible seam appears between the first and last views of a rotational sequence.** The axis calibration is stale, or the stage was disturbed between calibration and acquisition. Recalibrate the axis; enabling iterative refinement masks the symptom without fixing the cause.

**The merged cloud expands or contracts radially.** The commanded angular increment does not match the actual rotation, usually because the driver microstep setting differs from the value entered in the interface. Verify that the driver is set to 1600 pulses per revolution and that the gear ratio entered matches the fitted gears.

**Depth is correct in the center of the field and wrong at the edges.** The distortion model is under-constrained at the field edges. Acquire additional calibration images with the board near the corners and confirm that distortion compensation is enabled at reconstruction time.

**Specular highlights produce spikes or outliers.** Reduce exposure, cross-polarise if possible, or coat/matte the specimen where this is acceptable. Confidence thresholding removes most such points at the cost of coverage.

**Dark specimens are noisy even at long exposure.** Increase the number of patterns rather than the exposure. Independent realizations improve correspondence in a regime where longer integration only adds saturation risk.

**The turntable position drifts over a long acquisition.** The driver current is set too low for the load, or the gears are slipping. Check the gear mesh and the driver current setting.

**The interface does not detect one or both cameras.** Confirm the USB 3.0 connection and that no other application holds the camera. Close and reopen the camera connection from the interface before restarting the software.

#### Supplementary Note 8: Frequently asked questions

**How much does the system cost, and what dominates the cost?** Approximately 4000 Euro, including (german) taxes at the time of purchase, of which the two cameras and their lenses account for roughly 50 %. Everything else is printed parts, standard opto-mechanics and commodity electronics. To reduce the cost dramatically, albeit with a tradeoff in measurement quality, one can replace the machine vision cameras with consumer cameras or webcams.

**How long does one measurement take?** A single view is the projection of 100 patterns at the chosen exposure, so of the order of 10 s at the working settings. A seven-view rotational sequence is that multiplied by the number of views plus the stage moves.

**Do I need an optical table?** No. The breadboard is convenient and makes rebuilds reproducible, but any rigid flat surface works provided nothing moves between calibration and measurement.

**Can I use cheaper cameras or webcams?** In principle yes (see above), and the modular mounts permit it. The cost of doing so is accuracy, and we have not characterized that trade-off quantitatively.

**How large a specimen can be measured?** The measurement volume follows from the lenses and the working distance and is rescaled by changing them. The configuration characterized here accommodates plant organs, arthropods, small rodents and isolated organs, but even scaling to human size is possible, at the cost of resolution.

**Which specimens will not work?** Anything specular, transparent or volumetrically scattering. See **Supplementary Note 6**.

**Does the specimen need preparation?** Usually not. Matte, dry and static is ideal. Powdering or matting sprays improve difficult surfaces but alter the surface being measured and should be reported.

**How do I know whether to trust a given region of a scan?** Use the per-point correlation coefficient, which is retained in every export, and the fill fraction. Both are used as confidence channels throughout the analysis notebooks, and the murine section of the main text shows how they separate a measured surface from a scattering volume.

**Do I have to write code?** No. All six modules are reachable from the graphical interface. The analysis notebooks require running cells but not writing code, and each is configured from a single block at the top.

**How often must I recalibrate?** Whenever a camera, lens, focus, aperture or mount is changed. The axis calibration additionally requires the turntable to be undisturbed. We recommend calibrating regularly after the first initialization, as screws and parts might settle in and less, but regularly, in common usage.

**Can I compare measurements taken on two different builds?** Yes, provided each is calibrated and the analysis uses a common physical grid spacing and stated evaluation scales. Descriptors evaluated at a fixed number of pixels are not comparable between builds.

**Which file format should I export?** PLY preserves the per-point confidence value and is the format the analysis notebooks expect. OBJ, STL and XYZ are provided for interchange.

#### Supplementary Note 9: Analysis notebooks

Seven analysis workflows accompany the toolbox. Each runs from an export to finished figures and tables without intervention beyond selecting inputs and, where required, a region of interest or manually segmented structure. Each workflow writes its numerical outputs to file so that an analysis can be reproduced and audited after the run.

The reconstructed surfaces supporting this study are available at Zenodo under https://doi.org/10.5281/zenodo.22167250.[54] The analysis notebooks, environment specifications, derived data and per-panel source data are available at Zenodo under https://doi.org/10.5281/zenodo.22167598.[55]

**Leaf surface descriptors.** Input is the reconstructed leaf surface for each state. Output comprises the fraction of reconstructed surface visible from sampled viewing directions, rugosity and projection error for the best view, planarity and warp, scale-resolved roughness and curvature, venation relief and the distributions underlying the summary values.

**Murine limb geometry.** Input is the segmented distal and proximal limb parts. Output is the three-dimensional inter-segment angle, the corresponding projected angle over virtual viewing directions and five misalignment tolerances, sensitivity to segment-boundary placement, and limb surface area relative to its projection. The fixed laboratory reference surface is characterized separately and does not enter the three-dimensional inter-segment angle.

**Caudal annuli.** Input is a segmented tail cloud. Output is the annulus spacing from a two-dimensional spectrum with peak prominence, relief amplitude, a profile taken perpendicular to the crests, and the apparent spacing after projection into the image plane.

**Murine range-image quantification.** Input is one or more native correspondence exports. The workflow builds structured range images with the correlation coefficient retained as a confidence channel and produces per-view panels, surface-class readouts, ISO 25178 parameters, scale dependence of roughness, descriptor separability, and correlation and incidence envelopes.

**Lepidopteran projection and texture analysis.** Input is a single-view export with metric lateral coordinates. Output includes projected and reconstructed surface area, the implied projection error, sensitivity to boundary erosion and incidence cap, incidence-resolved area accounting, virtual-view apparent area, scale-resolved roughness and curvature, band-passed relief, normal dispersion, curvature-based shape classification and the radially averaged power spectrum.

**Rotational merging and export.** Input is a directory of single-view clouds together with a calibrated stage axis. Output is the merged cloud, loop-closure and inter-view diagnostics, the reconstructed mesh and exports in PLY, OBJ, STL and XYZ. ***Surface rendering.*** Input is any segmented point cloud. Output is a shaded surface rendering from a chosen viewpoint, with optional scalar quantities draped over the shading. This workflow is used for the three-dimensional surface displays in the manuscript.

#### Supplementary Note 10: Surface classification on the murine specimens

The murine section of the main text reports that local roughness provides strong contrast between exposed skin and fur without training data. This note gives the underlying quantities, which characterize the measurement rather than the animal.

Across the shaved views included in the surface-class analysis, exposed skin returns a local root-mean-square roughness of 127 to 198 μm and fur returns 307 to 704 μm, a contrast of 2.0 to 4.2 within a view. Skin values vary by 14.3 % across viewing directions and fur values by 25.4 %, so the fur estimate depends 1.8 times more strongly on camera position. The explanation is physical: fur is a scattering volume rather than a surface, so correlation returns a depth somewhere between the hair tips and the skin beneath, at a position that depends on viewing geometry. Rigid multi-view registration of such regions is ill-posed irrespective of specimen movement, which is why these surface-class measurements are quantified per view rather than merged.

Areal parameters computed to ISO 25178 sharpen the distinction (Supplementary Table 5). In the shaved views the smooth class has skewness near zero and kurtosis between 3.3 and 4.1, consistent with a compact residual distribution from a measured surface. The rough class has kurtosis between 73 and 126 and skewness from −6.7 to +1.9, consistent with a heavy-tailed decorrelation distribution rather than a single physical surface. Spatial statistics agree: the smooth class forms a few large, connected components holding most of its area, while the rough class fragments into hundreds of smaller components (**Supplementary Table 5**).

Ranked by area under the receiver operating characteristic, the strongest discriminator is the fill fraction, with an AUC of 0.988 to 0.996 across views. Normal dispersion follows at 0.906 to 0.966 and curvedness at 0.882 to 0.916, while curvedness evaluated at a coarser scale falls to 0.73 to 0.85. The cheapest quantity in the pipeline is therefore the most informative, with the shape descriptors corroborating rather than carrying the separation.

The unshaved views complete the picture. There the skin-like class returns 186 to 202 μm, at the upper end of the shaved range, and occupies a field fraction falling from 0.318 to 0.021 across the three directions, compared with 0.258 to 0.548 in the shaved measurements. Median correlation falls from 0.837 to 0.678 and the retained point fraction from 0.335 to 0.085. Shaving therefore does not improve the measurement of an already measurable surface so much as expose surface that was previously inaccessible behind fur.

Two internal controls support this interpretation. The rigid specimen mount is classified into the trustworthy regime wherever it is visible, with contiguous and correctly delineated boundaries, so the separation responds to surface physics rather than merely to the choice of threshold. The measurement envelopes also behave as expected: pooled across views, roughness falls as correlation increases and rises strongly with incidence angle. Full per-view values are provided in Supplementary Data 3.

#### Supplementary Note 11: Nature of the murine demonstration, limitations and requirements for a designed study

The murine section uses specimens from a genotyped cohort, but it is a measurement demonstration rather than a phenotype study. The distinction should be explicit because the dataset is small, the specimens are cadavers, and the largest differences in the measured limb angle arise from specimen positioning rather than genotype.

##### 11.1 What was measured

Four cadaveric male mice, two knockout and two wild-type littermates, all 63 weeks of age, were scanned from multiple directions. Fourteen views across the four animals show a hind limb sufficiently well to divide it into distal and proximal geometric segments. For each retained view we report the three-dimensional inter-segment angle, the same axes after projection into virtual camera planes, sensitivity to manual placement of the segment boundary, and limb surface area relative to its projection. The fixed laboratory reference surface is characterized separately for frame reproducibility but does not enter the three-dimensional inter-segment angle. Twelve tail segments provide the caudal-annulus analysis, of which eight pass the spectral-prominence criterion.

##### 11.2 What the measurement resolves

The per-animal mean inter-segment angles are 7.7, 21.2, 18.0 and 153.3 degrees. The standard deviation of these four means is 69.1 degrees, and their range is 145.6 degrees, dominated by how the cadavers were positioned. The meaning within-animal standard deviation is 7.4 degrees; this combines repeated-view variation, postural relaxation during thawing and, for the one animal contributing both sides, left-right differences. Displacing the manually placed segment boundary by ±2 mm changes the angle by a median of 3.3 degrees. A virtual ±15-degree change in viewing direction produces a median projected-angle spread of 5.3 degrees. Projection therefore contributes variation of the same order as the within-animal variation and larger than the median sensitivity to boundary placement. The reference plane is reproducible within a session to 0.02–0.64 degrees, but this is a property of the laboratory frame and is not an uncertainty term of the three-dimensional inter-segment angle.

##### 11.3 Why no phenotype is claimed

**Positioning dominates the dataset.** The largest difference, between an almost straight limb and strongly folded limbs, does not follow genotype. It follows how each cadaver was laid on the stage.

**The specimens are cadavers.** Limb posture in a thawing cadaver is not the load-bearing posture of a gait assay. The specimens relaxed during the sessions, and this contributes to the within-animal spread.

**The cohort is too small for inference.** There are two animals per genotype, no standardized positioning protocol, and no blinding or randomization of preparation. No statistical genotype comparison is appropriate, and none is presented.

**The assay is not the beam-walk assay.** The published foot-base angle is defined at a specific phase of locomotion. The quantity reported here is a geometric inter-segment angle in a fixed specimen. The purpose is to isolate projection geometry, not to reproduce the physiological endpoint.

##### 11.4 Descriptive checks

The descriptive checks do not reveal a genotype-consistent pattern. One knockout and one wild type occupy the strongly folded part of the postural range, while the almost straight limb belongs to the second wild-type animal. Accepted caudal-annulus spacings overlap across all four animals and give an overall mean of 1.39 ± 0.17 mm. These observations are reported to show what was inspected, not as evidence for equivalence between genotypes.

##### 11.5 What designed study would (at least) require

**1. A positioning protocol and positioning control.** A defined posture, a jig that reproduces it, and deliberate rescanning of at least one specimen in recorded alternative postures would turn the dominant preparation variable into a calibrated factor.

**2. Adequate group sizes and blinding.** Genotype should be concealed from the operator positioning the specimen and from whoever places segmentation boundaries, because both decisions affect the resulting geometry.

**3. Live acquisition for the gait endpoint.** Reproducing a beam-walk measurement requires reconstruction at a defined instant of unconstrained movement, which in turn requires faster projection/capture and motion-aware reconstruction.

**4. Landmark-based segmentation.** Shorter working distance and acquisitions designed for the paw would allow anatomical landmarks to replace the operator-defined segment cut and remove the boundary-placement term.

**5. A measured distribution of animal heading.** The projection term is currently evaluated across stated virtual misalignments. A designed locomotion study should record the actual distribution of body and limb orientation relative to the cameras.

#### Supplementary Note 12: The scale at which a surface area is defined

The area assigned to a reconstructed surface depends on the scale below which structure is not resolved. For a height field that scale enters through the width at which the field is smoothed before its gradient is taken, because the gradient sets the local foreshortening factor and therefore the area attributed to each lattice cell.

The dependence is large enough to matter. On the lepidopteran specimen, evaluating the gradient at Gaussian widths from the median point spacing of 0.112 mm to 1.6 mm gives rugosities of 1.126, 1.089, 1.066, 1.046, 1.042, 1.028 and 1.023, so the implied projection error falls from 11.2 % to 2.2 % across that range. On the leaf, roughness and curvature likewise change strongly with evaluation scale, and the direction of the curvature difference between fresh and desiccated states reverses between 0.3 and 4.8 mm.

Three consequences follow. First, a surface area, rugosity or projection error quoted without its evaluation scale is incomplete. Second, two reconstructions should be compared at a common physical scale supported by both rather than each at its own sampling limit. Third, evaluating below the sampling scale does not provide a finer measurement of the same surface; it increasingly measures numerical noise and undersmoothing. In the lepidopteran specimen, evaluating at 0.056 mm, half the median point spacing, raises rugosity to 1.173, and that value is shown only to demonstrate this sensitivity.

Surface-area and rugosity values in this work are therefore quoted at an explicitly stated supported scale, while roughness and curvature are shown across multiple physical evaluation scales where their scale dependence itself carries biological or methodological information.

#### Supplementary Tables

**Supplementary Table 1 | Bill of materials.** Parts required for one complete instrument. Prices are omitted pending a dated quotation.

At the time of purchase, approximately one half of the cost was located with the cameras and objectives at around 2000 € before tax, with the projector, assorted optical, electrical and mechanical parts making up the rest of the ~4000 Euro budget.

| Category | Item | Qty | Supplier or source | Note |
|---|---|---|---|---|
| Cameras | Alvium camera 1800 U-811c color | 2 | Allied Vision | RBG, USB 3.0 |
| Cameras | C-25-F1.8-10MP-T2-3 objective | 2 | Allied Vision | |
| Cameras | TR75V/M post | 2 | Thorlabs | |
| Cameras | PH50E/M post holder | 2 | Thorlabs | |
| Cameras | CF125C/M clamp | 2 | Thorlabs | Fixing to optical table |
| Cameras | M3 screw, 12 mm | 4 | Thorlabs | Camera to mount |
| Cameras | M6 screw, 16 mm, and washer | 2 | Thorlabs | |
| Cameras | M4 nut | 1 | Thorlabs | Mount to post |
| Cameras | 3D printed camera mount | 2 | Printed | |
| Cameras | USB Micro-B cable | 2 | Retailer | USB 3.0 or higher |
| Projector | TX-127 projector | 1 | Retailer | |
| Projector | HDMI cable | 1 | Retailer | |
| Turntable | Raspberry Pi Pico W | 1 | Retailer | Control unit |
| Turntable | USB cable, micro-USB to USB | 1 | Retailer | |
| Turntable | Jumper cable set, male to male | 1 | Retailer | |
| Turntable | NEMA 17 stepper motor | 1 | Retailer | |
| Turntable | Two-phase microstep driver | 1 | Retailer | DM430 or equivalent |
| Turntable | Electronic breadboard | 1 | Retailer | |
| Turntable | Lazy-Susan ball bearing | 1 | Retailer | |
| Turntable | Power supply, 12 V 1 A, with adapter | 1 | Retailer | |
| Turntable | 3D printed base | 1 | Printed | |
| Turntable | 3D printed gear, large, 120 teeth | 1 | Printed | |
| Turntable | 3D printed gear, small, 25 teeth | 1 | Printed | |
| Turntable | 3D printed motor cover | 1 | Printed | |
| Turntable | M4 screw, 6 mm | 4 | Thorlabs | Bearing to base |
| Turntable | M6 screw, 16 mm, and washer | 4 | Thorlabs | Base and motor to table |
| Optional | Aluminum breadboard | 1 | Retailer | Mobility and reproducible rebuilds |

| | | | | |
|---|---|---|---|---|
| Optional | Lens, 500 mm focal length | 1 | Retailer | Shortens projector throw where space is limited |
| Computing | Control computer | 1 | | |

**Supplementary Table 2 | Acquisition and reconstruction parameters.**

| Parameter | Plane | Rotational object | Leaf | Murine | Lepidopteran |
|---|---|---|---|---|---|
| Pattern size (px) | 1000 × 1000 | 1000 × 1000 | 1000 × 1000 | 1000 × 1000 | 1000 × 1000 |
| Feature size (px peak-to-peak) | swept 5–300; working 50 | 50 | 50 | 50 | 50 |
| Patterns per view | swept 1–1000; working 100 | 75 | 100 | 100 | 100 |
| Exposure (ms) | swept 2–200; working 25 | 25 | 25 | 25 | 25 |
| Correlation threshold at reconstruction | 0.01 | 0.01 | 0.01 | 0.01 | 0.01 |
| Primary analysis/export gate | geometric bulk classification; no correlation used | 0.50 | none; geometric cleaning and color-based holder removal | 0.25 export; 0.50 for range-image analysis | 0.70 |
| Search offsets L/R/T/B (px) | 450/450/10/10 | 450/450/10/10 | 450/450/10/10 | 450/450/10/10 | 450/450/10/10 |
| Sub-pixel interpolations | 3 | 3 | 3 | 3 | 3 |
| Distortion compensation | on | on | on | on | on |
| Outlier rejection | LMedS | LMedS | LMedS | LMedS | LMedS |
| Views / angular sampling | 1 acquisition per sweep setting | 7 views at 60° | full rotational scan in each of 2 states | multiple views; 14 limb views retained | 1 view |
| Iterative refinement | not applicable | disabled | not reported / not required for descriptor analysis | not applied to the single-view analyses reported | not applicable |

**Supplementary Table 3 | Instrument performance summary.** Each entry is a single acquisition.

| Section | Metric | Current value |
|---|---|---|
| Planar reference | Working configuration | 25 ms, 100 patterns, 50 px peak-to-peak |
| Planar reference | Robust residual, full field / patch | 145 / 106 µm |
| Planar reference | Local flatness, full field / patch | 82 / 45 µm |
| Planar reference | Completeness, full field / patch | 99.2 / 100.0 % |
| Exposure sweep | Stable residual range, 4–75 ms | 142–159 µm full field; 102–119 µm patch |
| Exposure sweep | Patch local flatness, 4–75 ms | 40–49 µm |
| Exposure sweep | Completeness | 99.6 % at 4 ms; 83.1 % at 75 ms; full field 69.8 % and patch 86.6 % at 150 ms; full field 51.9 % at 200 ms |
| Pattern-number sweep | 2 patterns | 475 µm residual at 11.6 % completeness |
| Pattern-number sweep | Recovery / plateau | 169 µm at 10 patterns; 148 µm at 30 patterns; 93.1 % completeness at 40 patterns; plateau at ~30–40 patterns |
| Feature-size sweep | Stable range, 20–100 px | 137–151 µm full field; 95–111 µm patch; completeness >93.6 % |
| Feature-size sweep | 5 px | 265 µm residual |
| Feature-size sweep | 10 px selective-survival case | 83 µm patch residual at 34.1 % patch completeness |
| Feature-size sweep | 300 px | 210 µm residual |
| Stage-axis calibration | Mean radial residual | 0.32 mm |
| Rotational registration | Loop closure | median 156 µm; mean 160 ± 116 µm; 95th percentile 346 µm; maximum 2.99 mm; 546,000 point pairs |
| Rotational registration | Relative loop closure | ~1 part in 1300 of a 205 mm object diagonal |
| Merged rotational cloud | Sampling / roughness | 2.94 million points; median nearest-neighbor spacing 113 µm; median local roughness 43 µm |

**Supplementary Table 4 | Summary of murine surface-class measurements**

| Subset | Readout | Value |
|---|---|---|
| Shaved views | Exposed-skin local RMS roughness | 127–198 μm |
| Shaved views | Fur local RMS roughness | 307–704 μm |
| Shaved views | Within-view roughness contrast | 2.0–4.2-fold |
| Shaved views | Variation across viewing directions | skin 14.3 %; fur 25.4 %; fur varies 1.8-fold more strongly |
| Unshaved views | Skin-like class roughness | 186–202 μm |
| Unshaved views | Skin-like field fraction across three directions | 0.318 to 0.021 |
| Shaved comparison | Skin-like field fraction | 0.258 to 0.548 |
| Unshaved views | Median stereo correlation | 0.837 to 0.678 |
| Unshaved views | Retained point fraction | 0.335 to 0.085 |

**Supplementary Table 5 | ISO 25178 areal parameters by class, shaved murine views.** Ranges over the six views.

| Parameter | Smooth class | Rough class |
|---|---|---|
| Sa (μm) | 111.9 to 149.2 | 458.8 to 981.1 |
| Sq (μm) | 146.9 to 190.6 | 851.4 to 2831 |
| Ssk | −0.185 to −0.008 | −6.72 to +1.87 |
| Sku | 3.32 to 4.08 | 73.3 to 125.6 |
| Sdq | 0.676 to 1.607 | 0.918 to 3.640 |
| Curvedness ($m^{-1}$) | 51.9 to 70.7 | 198.5 to 274.8 |
| Normal dispersion (°) | 3.26 to 4.26 | 9.54 to 17.34 |
| Fill fraction | 1.000 in all views | 0.694 to 0.752 |
| Connected components | 29 to 51, largest five hold 92 to 99 % | 202 to 847, largest five hold 18 to 40 % |

**Supplementary Table 6 | Leaf surface descriptors, fresh and desiccated.** Both states resampled onto a common 0.15 mm grid.

| Descriptor | Unit | Fresh | Desiccated | Change / comment |
|---|---|---|---|---|
| Rugosity R | – | 1.326 | 1.690 | +27.5 % |
| Area underestimated by projection at best viewing direction | % | 24.6 | 40.8 | +16.2 percentage points |
| Best single-view recorded fraction | % of reconstructed two-sided surface | 39.8 | 33.6 | −6.2 percentage points |
| Median single-view recorded fraction | % | 14.6 | 21.5 | +6.9 percentage points |
| Worst single-view recorded fraction | % | 2.5 | 16.3 | +13.8 percentage points |
| Area recovered in full rotational reconstruction | relative to fresh | 100 % | 37.3 % | −62.7 % |
| Projected footprint at best viewing direction | relative to fresh | 100 % | 24.7 % | −75.3 % |
| Planarity index | 0–1 | 0.832 | 0.624 | lower after drying |
| Local warp RMS at 4.8 mm | mm | 1.41 | 5.79 | ×4.11 |
| Local warp kurtosis | – | 4.0 | 8.7 | higher after drying |
| Rq at 0.3 mm | mm | 0.518 | 2.681 | ×5.18 |
| Rq at 4.8 mm | mm | 0.720 | 4.813 | ×6.68 |
| Mean absolute curvature at 0.3 mm | $mm^{-1}$ | 0.770 | 1.011 | ×1.31 |
| Mean absolute curvature at 4.8 mm | $mm^{-1}$ | 0.077 | 0.042 | ×0.55 |
| Venation coverage | % of lamina | 6.6 | not separable | – |
| Median venation relief height | µm | 597 | not separable | – |

*Absolute area changes are reported as changes in recovered area, not as tissue loss. Folding and self-occlusion make the desiccated reconstruction incomplete even after a full rotational scan.*

**Supplementary Table 7 | Sensitivity of the lepidopteran projection error to the two analysis conventions.** Rugosity R and the implied projection error, for boundary erosion widths in lattice cells and incidence caps in degrees.

| Quantity | Range tested | Quoted / key value | Interpretation |
|---|---|---|---|
| Boundary erosion width | 0–5 lattice cells | 1 cell | tested jointly with incidence cap |
| Incidence cap | 60–90° | 75° | cells above the cap excluded |
| Projection error at quoted convention | – | 11.2 % | rugosity R = 1.126 at 0.112 mm evaluation scale |
| Projection-error range across tested convention grid | – | 7.5–15.8 % | magnitude remains stable, exact value convention-dependent |
| Evaluation scale used for quoted area | 0.056–1.6 mm explored | 0.112 mm | median point spacing; finest supported scale |
| Below-sampling sensitivity point | 0.056 mm | R = 1.173 | shown only to illustrate undersmoothing |
| Coarser-scale examples | 0.2 / 1.6 mm | R = 1.089 / 1.023 | projection error decreases with smoothing scale |

#### Supplementary Figures

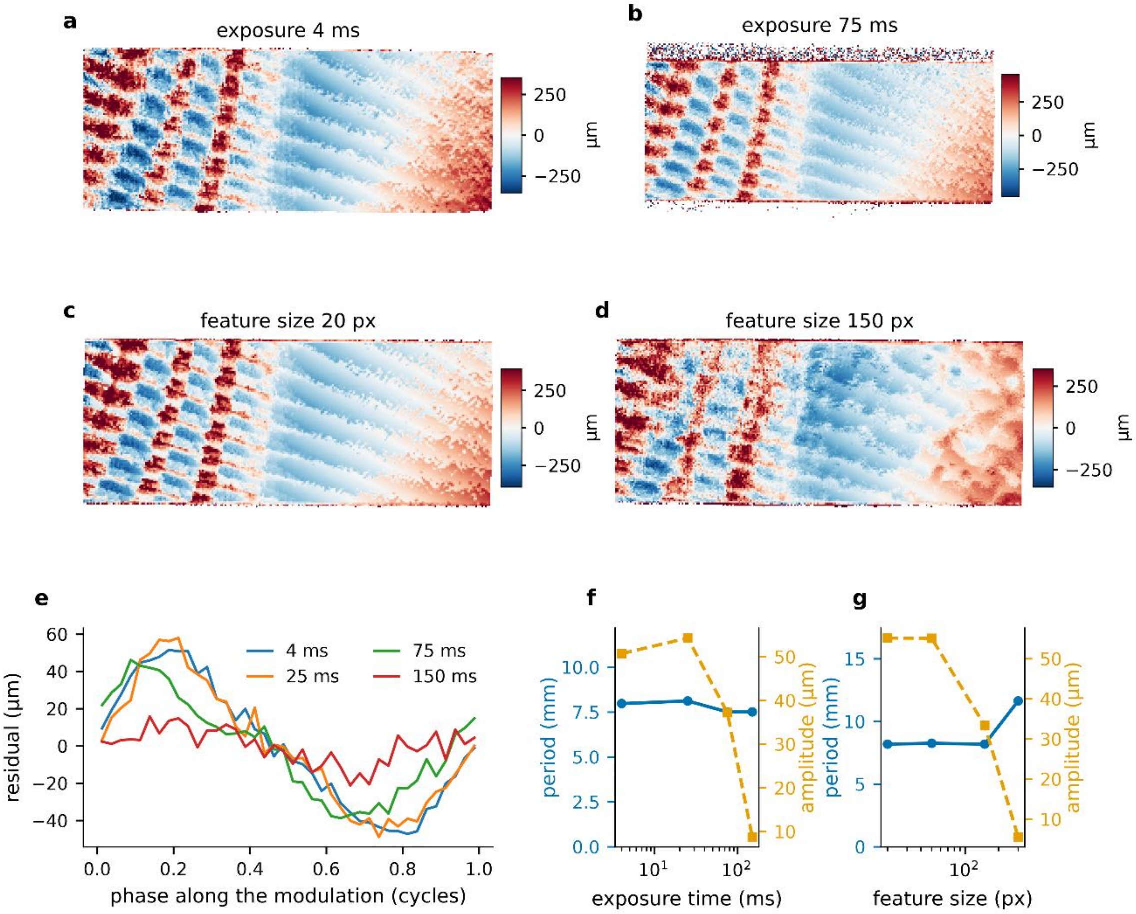


**Supplementary Fig. 1 | The periodic modulation in the plane residual.** Residual to the fitted plane for the planar reference, at two exposure times (**a**, **b**) and two pattern feature sizes (**c**, **d**), on a common color scale, accentuated for visibility. **e**, The residual averaged by phase along the modulation, for four exposure times. **f**, **g**, Period of the modulation (blue, left axis) and its amplitude (orange, right axis) against exposure time and against feature size. The period stays between 7.5 and 8.3 mm and the orientation within two degrees across both sweeps, while the amplitude falls from 51 to 9 µm as the exposure rises from 4 to 150 ms and from 55 to 33 µm as the feature size rises from 20 to 150 px. At 300 px no modulation is detectable and the values plotted are those of the noise floor. n = 1 planar reference, one acquisition per setting.

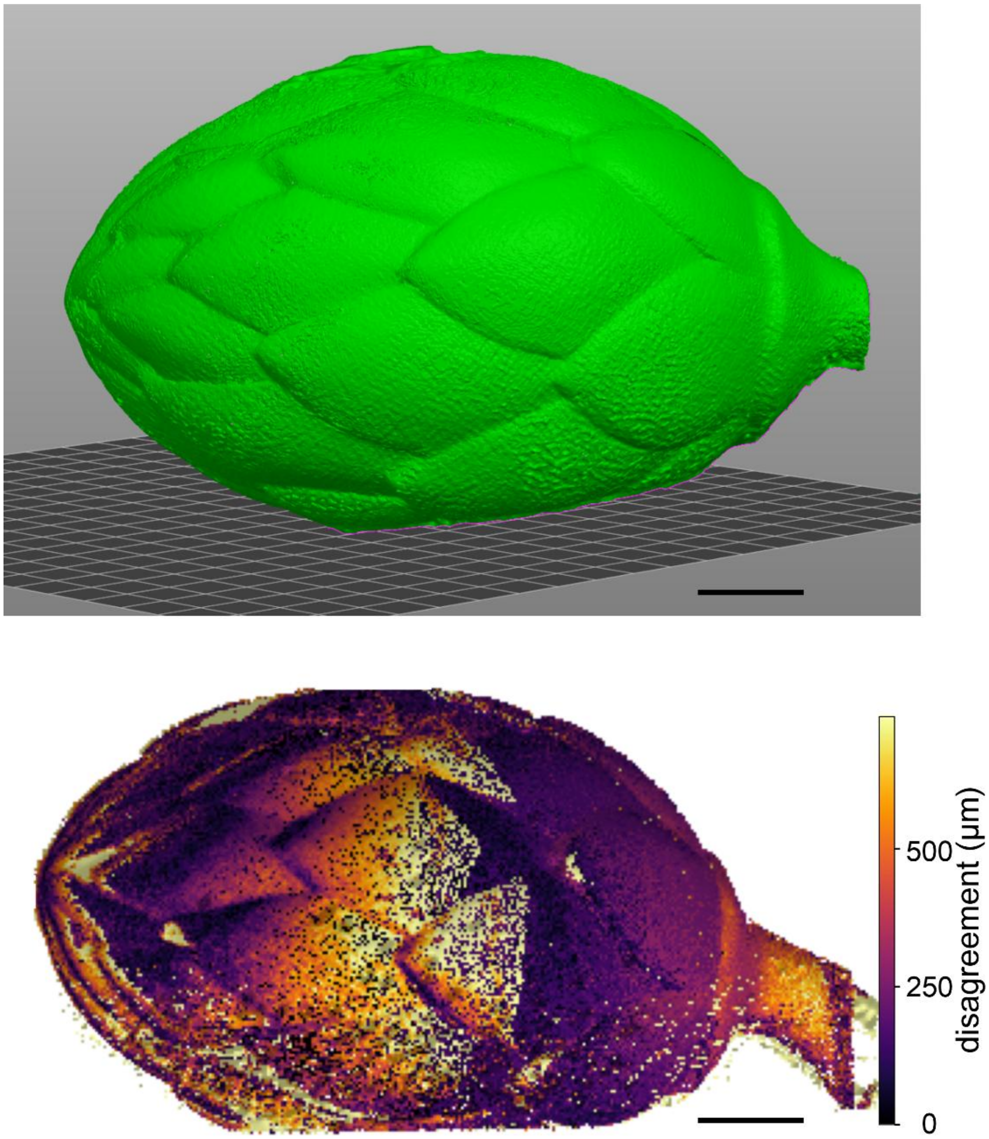


**Supplementary Fig. 2 | Meshed output of the rotational scan.** The merged reconstruction of the hop cone after screened Poisson reconstruction, shown as exported for downstream use, and the same reconstruction color-coded by inter-view disagreement. The mesh is not watertight, because a rotational sequence cannot observe the supported underside of the object. Scale bars, 20 mm.

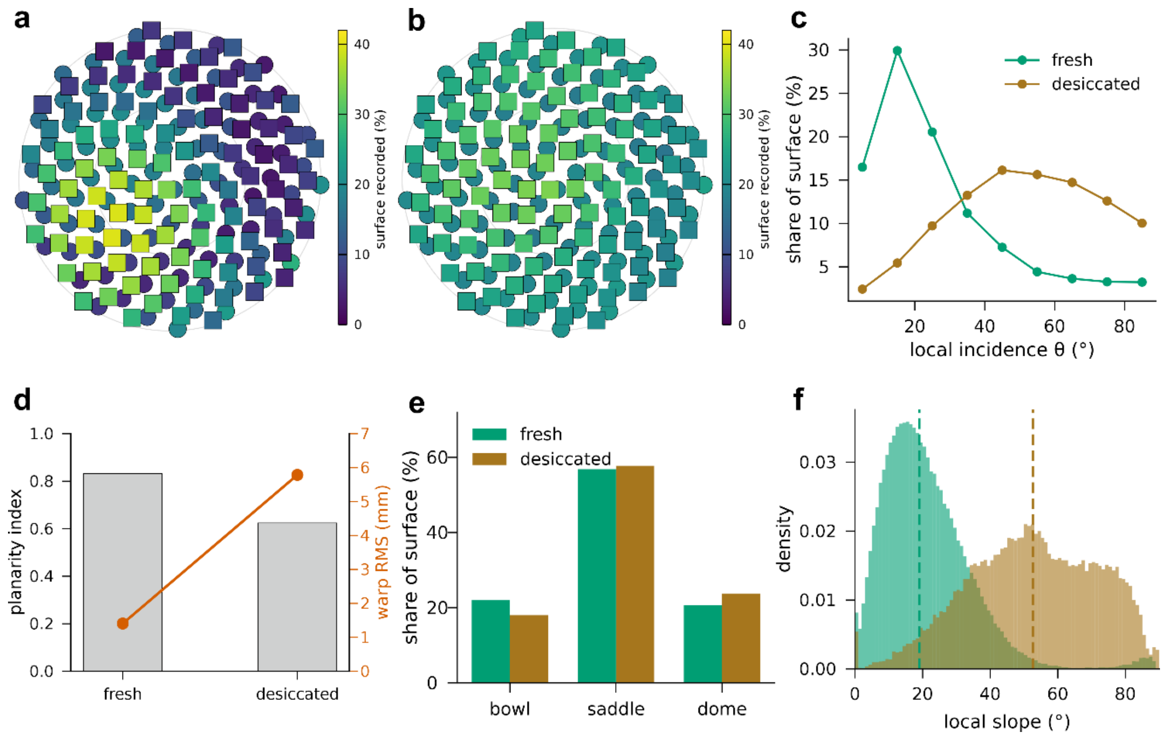


**Supplementary Fig. 3 | Leaf surface, extended descriptors.** Fresh and desiccated states of the same leaf. **a**, **b**, Fraction of the reconstructed surface recorded from each viewing direction, in a Lambert azimuthal equal-area projection of the viewing sphere, with circles for the upper hemisphere and squares for the lower. **c**, Share of the reconstructed surface against local incidence angle. **d**, Planarity index and warp amplitude. **e**, Local shape classified as bowl, saddle or dome from the signs of the principal curvatures. **f**, Distribution of local slope, with the median for each state marked. n = 1 leaf measured in two states.

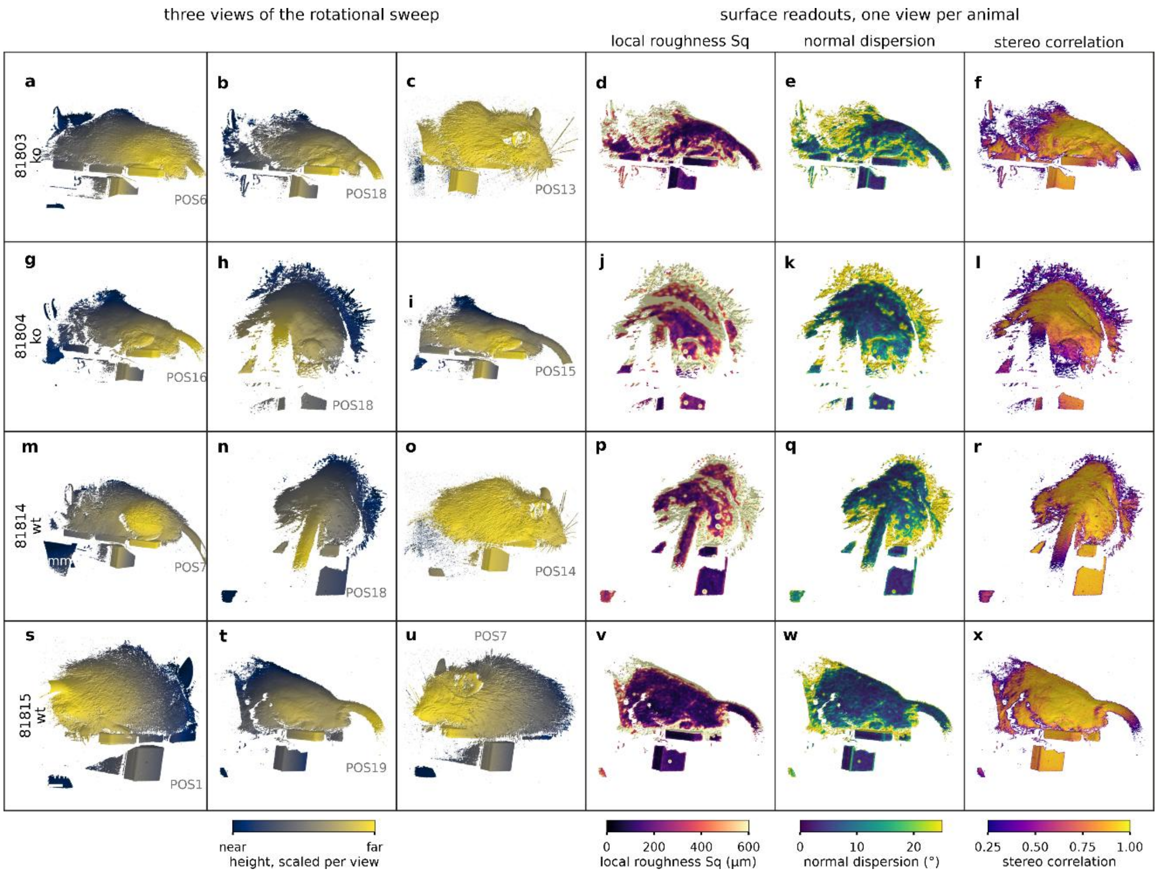


**Supplementary Fig. 4 | Surface readouts across the four murine specimens.** Rows are animals, knockouts above wild types. The first three columns show three views of the rotational sweep for each animal, rendered as shaded surfaces and color-coded by height scaled per view, with the position number given in each panel. The remaining columns show local root-mean-square roughness, dispersion of the surface normal and stereo correlation, computed on the best-conditioned view of each animal and displayed on color scales shared across all four. Exposed skin returns roughness close to the reconstruction noise floor while fur saturates the scale, and the correlation channel marks the same regions as unreliable without reference to the roughness. Scale bars, 20 mm. n = 4 animals.

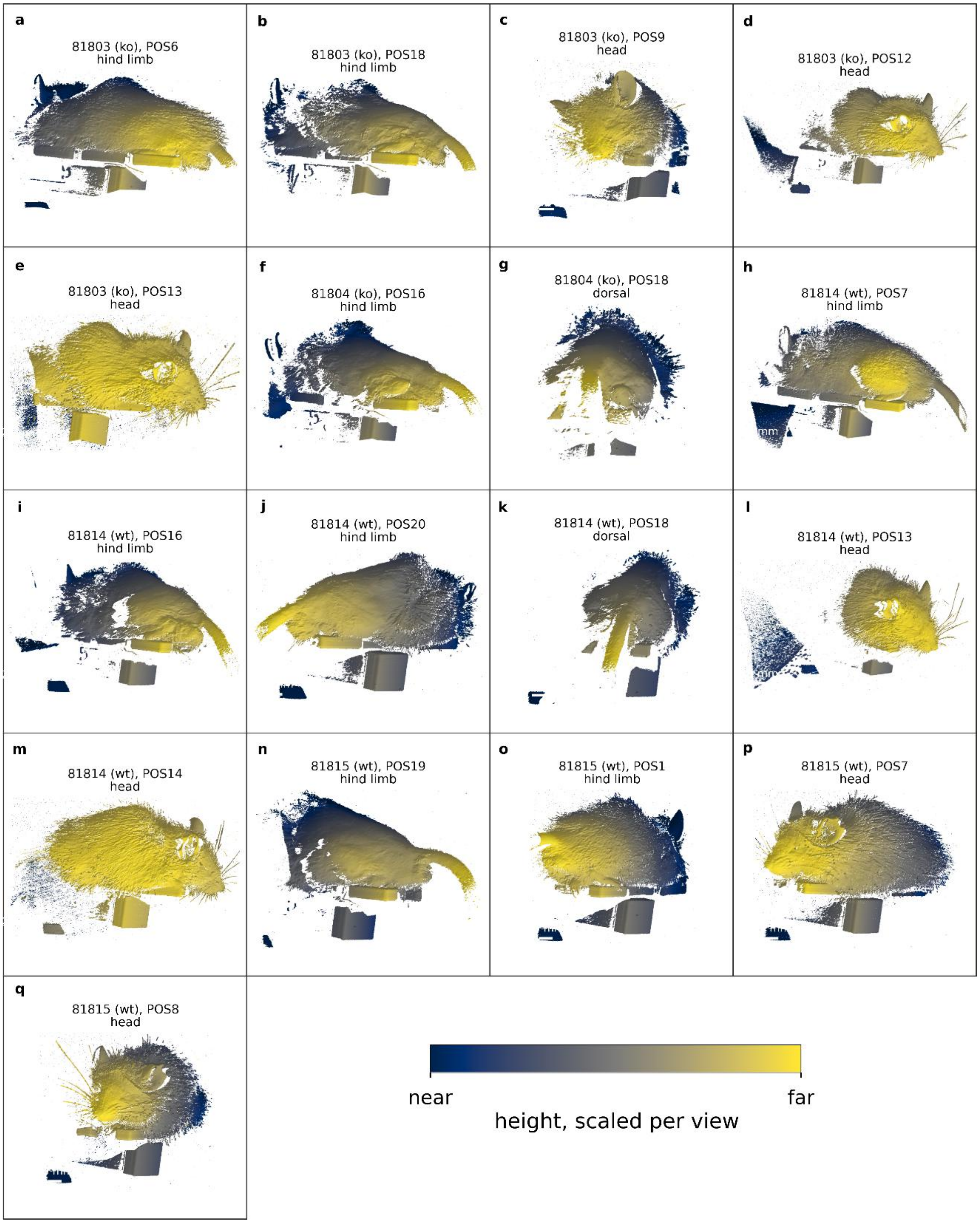


**Supplementary Fig. 5 | Gallery of murine reconstructions.** Seventeen views across the four animals, rendered as shaded surfaces color-coded by height scaled per view and labelled by animal, genotype, position and aspect. Individual vibrissae, the ear pinna and the caudal annuli are resolved from geometry alone, without color or texture information. Scale bars, 20 mm. n = 4 animals.

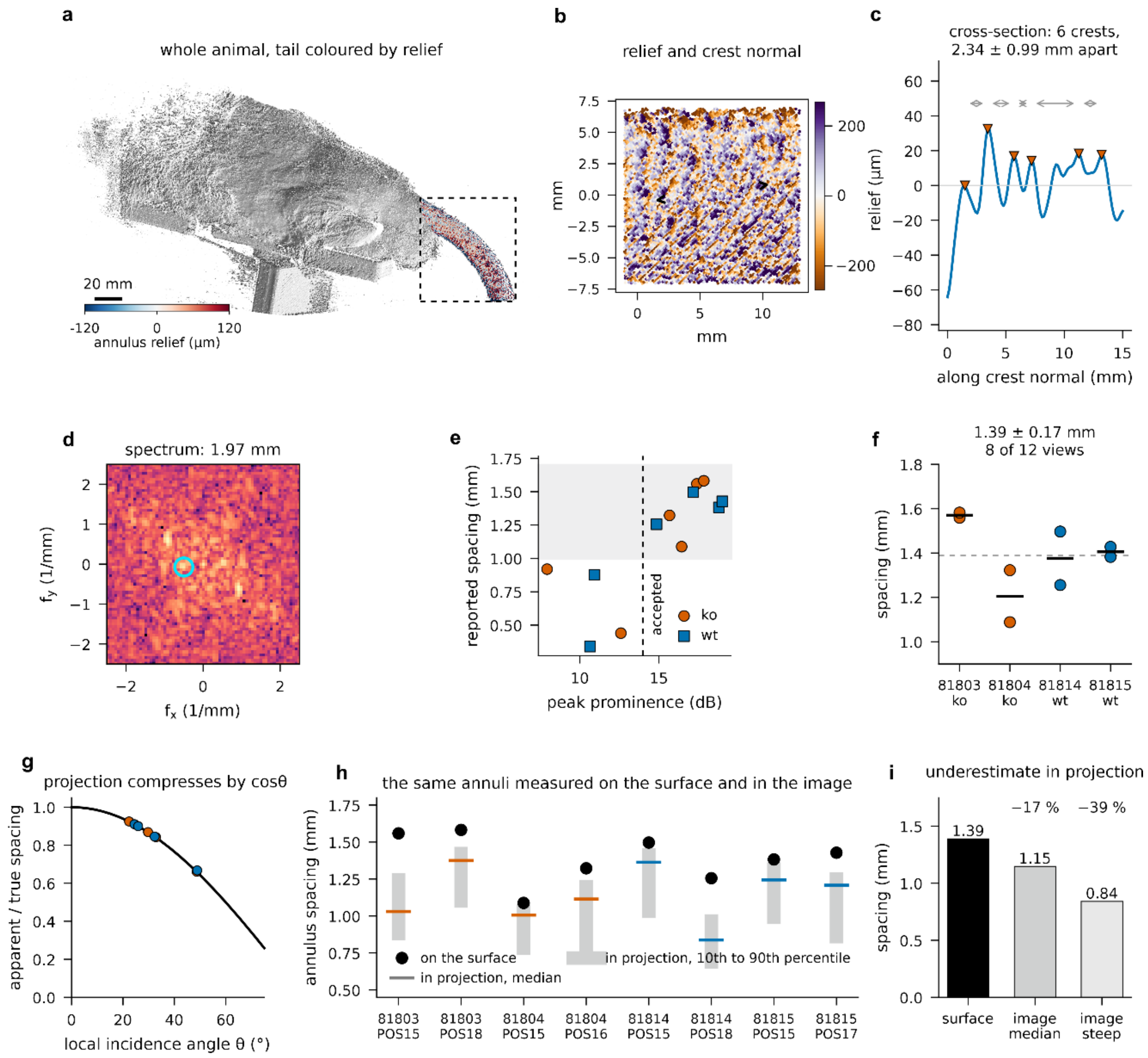


**Supplementary Fig. 6 | Caudal annuli resolved on the tail surface.** **a**, Whole animal rendered as a shaded surface with the segmented tail colored by annulus relief; the dashed box marks the tail. **b**, Detail of the relief within a window, with the crest normal recovered from the local two-dimensional spectrum. **c**, Profile taken perpendicular to the crests, with individual crests marked and their spacings annotated. **d**, Two-dimensional power spectrum of the relief in the same window. **e**, Reported spacing against peak prominence for all twelve tail segments; measurements below the 14 dB criterion return spacings outside the range recovered by the accepted ones. **f**, Accepted spacings by animal, mean 1.39 ± 0.17 mm across eight views. **g**, Compression of an apparent spacing measured in projection, as the cosine of the local incidence angle, with the eight accepted views marked. **h**, Surface spacing against the spacing an image would report, per view, as median and 10th to 90th percentile. **i**, Summary: the same annuli measured on the surface, in projection at the median incidence, and in projection where the surface is steep. Scale bars, 20 mm (**a**) and 5 mm (**b**). n = 4 animals, 12 tail segments of which 8 included in the analysis.

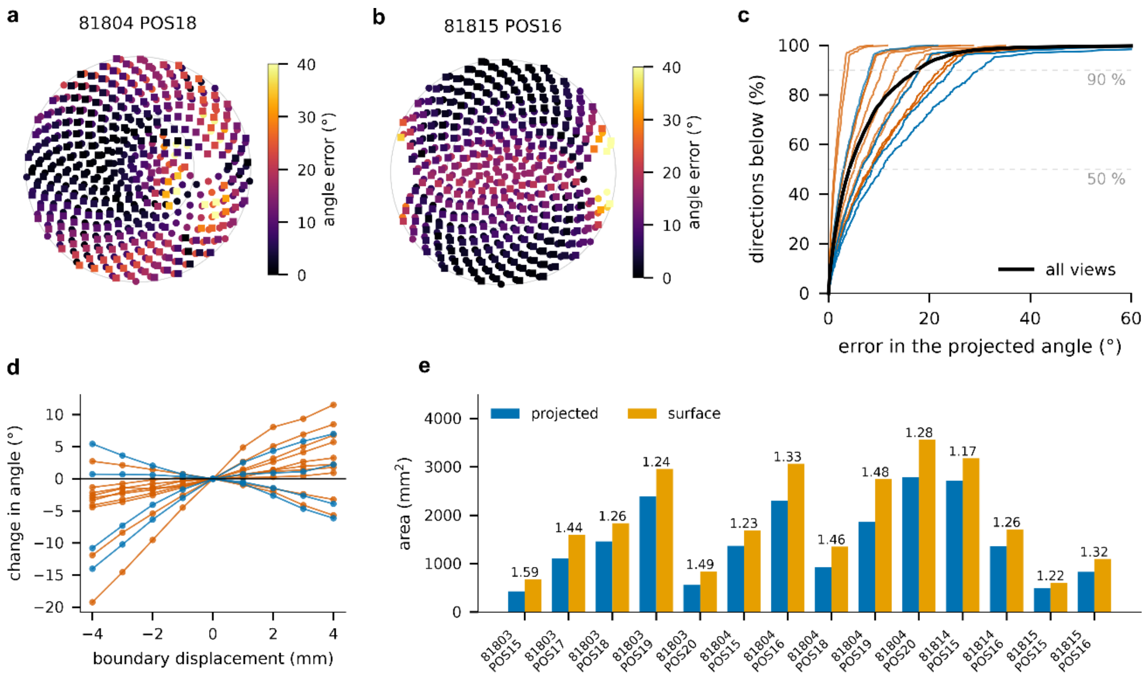


**Supplementary Fig. 7 | Viewpoint dependence of the murine limb angle. a**, **b**, Deviation of the projected inter-segment angle from its three-dimensional value, over 600 viewing directions distributed on the sphere, for two limbs, in a Lambert azimuthal equal-area projection with circles for the upper hemisphere and squares for the lower. **c**, Cumulative distribution of that deviation for all fourteen views, colored by genotype, with the pooled curve in black. **d**, Change in the measured angle when the manually placed segment boundary is displaced along its own normal, relative to the unshifted value, one line per view. **e**, Projected against surface area of the limb per view, with the ratio above each pair. n = 4 animals, 14 views.

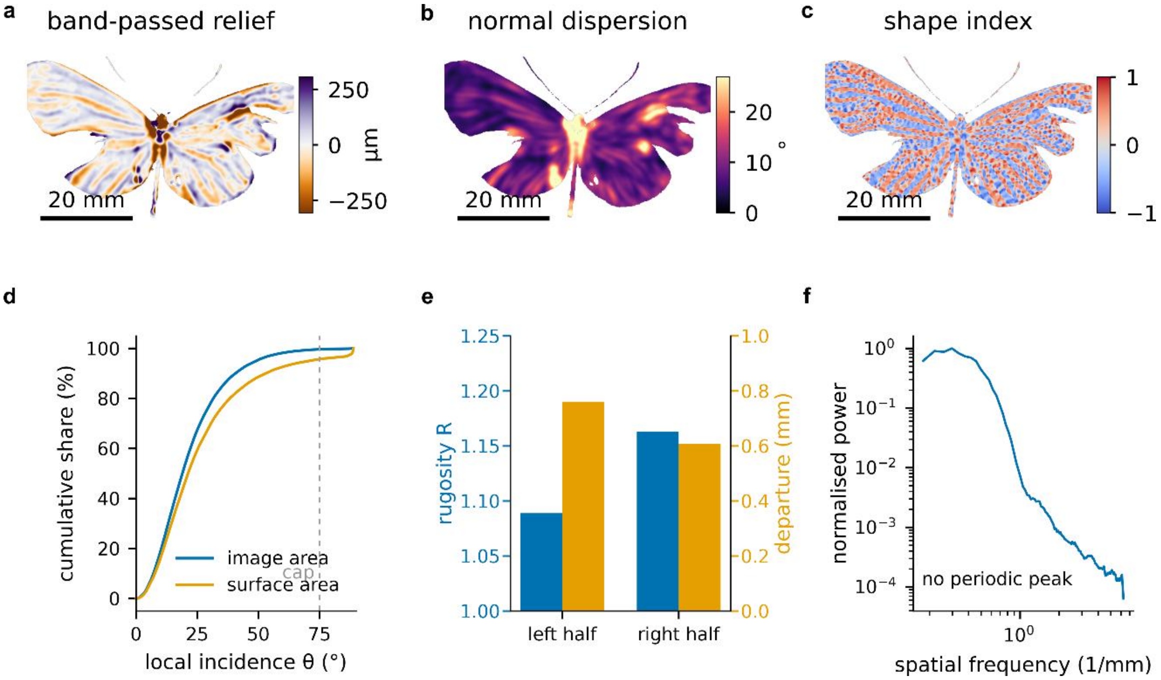


**Supplementary Fig. 8 | Texture and geometry of the lepidopteran specimen. a**, Band-passed relief, resolving venation and wing margins. **b**, Dispersion of the surface normal, a measure of how well ordered the surface is at the evaluation scale. **c**, Shape index, classifying local form between concave and convex. **d**, Cumulative share of the image area and of the reconstructed surface area against local incidence angle, with the 75 degree cap marked. **e**, Rugosity and departure from a fitted plane for the two halves of the specimen. **f**, Radially averaged power spectrum of the band-passed relief, which carries no peak above the local trend and is the reason no characteristic spacing is quoted for the scale rows. Scale bars, 20 mm. n = 1 specimen, single view.

**Supplementary Fig. 9 | Assembly photographs.** All assembly photographs are depicted in line of the **Supplementary Notes 1-4**.

#### Supplementary Data

The reconstructed surfaces supporting this study are available at Zenodo under https://doi.org/10.5281/zenodo.22167250.[54]

Source data for all graph panels are provided with this paper; for panels showing rendered surfaces, the underlying reconstructions are in the same record.

The analysis notebooks, environment specifications, derived data and per-panel source data are available at Zenodo under https://doi.org/10.5281/zenodo.22167598.[55]

**Supplementary Data 1 | Source data for the planar reference and the rotational scan.** Tabulated residual, local flatness and completeness for both regions of interest at every setting of the three parameter sweeps, and the per-pair overlap fraction and agreement for all twenty-one view pairs of the rotational scan.

**Supplementary Data 2 | Leaf surface descriptors.** Full descriptor table for both states, the recorded fraction for every viewing direction sampled, and the venation segmentation statistics.

**Supplementary Data 3 | Murine per-view metrics, limb geometry and tail spectra.** Per-view acquisition geometry and areal parameters, the limb table for all fourteen retained views giving the three-dimensional and projected angles at five misalignment tolerances, the deviation over the sphere of viewing directions, the boundary sensitivity, the reference-plane normals and residuals, the segment extents and the limb surface and projected areas, and the tail table giving spacing, peak prominence, relief and the apparent spacing in projection for all twelve tail segments.

**Supplementary Data 4 | Lepidopteran analysis outputs.** Headline quantities, the incidence-resolved area accounting, the rugosity against evaluation scale, the convention sensitivity, the scale-resolved roughness and curvature, and the apparent area for every viewing direction sampled.
