## supplementary and_figure source data for "Democratizing three-dimensional surface phenotyping: an open structured-light platform reveals and removes the projection bias in biological imaging": README.md

# Supplementary Data and figure source data

Four workbooks, one per Supplementary Data item. Each opens with a README sheet naming
every other sheet and the figure panel it supports.

| File | Item |
|---|---|
| `Supplementary_Data_1_planar_reference_and_rotational_scan.xlsx` | Supplementary Data 1 |
| `Supplementary_Data_2_leaf_surface_descriptors.xlsx` | Supplementary Data 2 |
| `Supplementary_Data_3_murine_metrics.xlsx` | Supplementary Data 3 |
| `Supplementary_Data_4_lepidopteran_analysis.xlsx` | Supplementary Data 4 |

The folder `csv/` carries the same tables as plain semicolon-separated text with a
byte-order mark, for readers who prefer them. File names give the figure and panel they
support, for example `Figure3_panele_exposure_sweep.csv`.

Distributions are given as histogram bin centres and counts matching the figure, because
the underlying arrays run to millions of points and are contained in the deposited
reconstructions. Panels that are renderings or maps have no tabular source data; the
reconstructions are the source for those and the notebooks regenerate them.

Figure numbering follows the final scheme of the manuscript: main Figures 1 to 7 and
Supplementary Figures 1 to 9.

Figure 1 has no separate source data. Panel a carries no plotted quantities beyond the
specimen extents and sampling values stated in the panel itself. Panels b and c replot
values presented in Figures 5 to 7 and in Supplementary Figure 7, whose source data are
in this deposit.
